# Close-kin methods to estimate census size and effective population size

**DOI:** 10.1101/2021.01.19.427337

**Authors:** Robin S. Waples, Pierre Feutry

**Affiliations:** Northwest Fisheries Science Center, National Marine Fisheries Service, National Oceanic and Atmospheric Administration, Seattle, WA, 98112 USA; Commonwealth Scientific and Industrial Research Organisation, Oceans & Atmosphere, GPO Box 1538, Hobart 7001, TAS, Australia

**Keywords:** siblings, parentage analysis, reproductive skew, mark-recapture, genetics, abundance

## Abstract

The last two decades have witnessed rapid developments and increasing interest in use of (1) genetic methods to estimate effective population size (*N*_*e*_) and (2) close-kin mark-recapture (CKMR) methods to estimate abundance based on the incidence of close relatives. Whereas *N*_*e*_ estimation methods have been applied to a wide range of taxa, all CKMR applications to date have been for aquatic species. These two fields of inquiry have developed largely independently, and this is unfortunate because deeper insights can be gained by joint evaluation of eco-evolutionary processes. In this synthesis, we use simple analytical models and simulated pedigree data to illustrate how various factors (life-history traits; patterns of variation in individual reproductive success; experimental design; stochasticity; marker type) can affect performance of the estimators. We show that the *N*_*e*_/*N* ratio and the probability of a close-kin match both depend on a vector of parental weights that specify relative probabilities that different individuals will produce offspring. Although age-specific vital rates are central to both methodologies, for CKMR they can potentially bias abundance estimates unless properly accounted for, whereas they represent the signals of genetic drift that *N*_*e*_ estimation methods depend upon. Coordinating *N*_*e*_ and CKMR estimation methods using the same or overlapping datasets would facilitate joint evaluation of both the ecological and evolutionary consequences of abundance.

## 1 INTRODUCTION

Quantifying abundance is central to many aspects of organismal biology, with two metrics being particularly important. The number of individuals (census size, *N*) informs a wide range of studies in basic and applied ecology, including population growth rates, competition and predation, immigration and emigration, behavior, extinction risk, harvest management, and ecosystem function (Krebs 2009). The evolutionary analogue of census size is effective population size (*N*_*e*_), which determines rates of genetic drift, inbreeding, and loss of genetic variability and mediates the effectiveness of natural selection (Charlesworth 2009). *N*_*e*_ is usually smaller than *N* and sometimes much smaller (Frankham 1995; Hauser and Carvalho 2008; Palstra and Fraser 2012), which means that some abundant populations could nevertheless experience rapid genetic change—a scenario that has been proposed for many marine species with huge populations and high fecundity (Hedgecock and Pudovkin 2011).

Both key population-size metrics are challenging to estimate in nature. Many animal species are vagile, cryptic, or rare, making detection and enumeration difficult (Seber 1986; Taylor and Wade 2000; Holmes 2001; Doak et al. 2005), and this is particularly true in the marine realm (**“**Managing fisheries is hard: it’s like managing a forest, in which the trees are invisible and keep moving around;” J. Shepard; http://jgshepherd.com/thoughts/). Effective population size is defined in terms of key demographic parameters (mean and variance in offspring number; Crow and Denniston 1988, Caballero 1994), but collecting these data often entails formidable logistical challenges, especially for long-lived species. As a consequence, indirect genetic methods have been used to estimate *N*_*e*_ since the early 1970s (Krimbas and Tsakas 1971; Luikart et al. 2010). Similarly, mark-recapture (MR) methods to estimate abundance have been a cornerstone of wildlife ecology for the past half century (Jolly 1965; Seber 1982; Williams et al. 2002). For species that are difficult to capture and/or physically tag, naturally occurring genetic markers can be used to identify “recaptures” of the same individual from passively-collected samples such as feces or hair. For example, Kendall et al. (2008) used genotypes derived from hair samples to obtain a robust estimate of abundance of a threatened population of grizzly bears (*Ursus arctos*) in Glacier National Park, USA, and Buckworth et al. (2012) used ‘genetagging’ to estimate harvest rate and catchability in Spanish mackerel (*Scomberomorus commerson*) from northern Australia.

More recently, researchers have begun to capitalize on the realization that, if naturally-occurring genetic marks are used, it is not necessary to capture individuals more than once, and the single “capture” can simply involve collecting DNA from a dead individual (or non-lethally from a living one). Close kin can also be used in a MR framework because individuals “mark” their relatives with shared genes (Skaug 2001). The two kinship categories that have been used to date with close-kin mark-recapture (CKMR) are parent-offspring pairs (POPs) and siblings. Successful application of CKMR methods to a number of fish species (Rawding et al. 2014; Bravington et al. 2016a, 2019; Hillary et al. 2018; Bradford et al. 2018; Ruzzante et al. 2019; Thomson et al. 2020), together with refinements in statistical methodology (Bravington et al. 2016b; Skaug 2017; Conn et al. 2020), have generated a great deal of interest in using this approach more broadly to study other taxa (e.g., Stewart et al. 2018; Oleksiak and Rajora 2020). Basic principles of CKMR have also been used to place an upper bound on the pre-Columbian human population size in the Caribbean (Fernandes et al. 2020).

Considerable advances in the estimation of effective size have also been made over the past ∼15 years. Prior to the mid-2000s, most genetic estimates of *N*_*e*_ used the temporal method (KIrimbas and Tsakas 1971; Nei and Tajima 1981), which requires at least two samples spaced in time. Most recent applications, however, use one of two single-sample methods (Palstra and Fraser 2012): a bias-adjusted method based on linkage disequilibrium (LD; Waples and Do 2008) or a method based on frequencies of siblings (Wang 2009), and these methods have been widely applied to marine, freshwater, and terrestrial species. Precision and bias of both single-sample methods has been extensively evaluated using simulated data (Waples and Do 2010; Tallmon et al. 2010; England et al. 2011; Gilbert and Whitlock 2015; Wang 2016), as have effects of age structure on LD estimates (Robinson and Moyer 2013; Waples et al. 2014).

Although these recent advances in methods for estimating *N* and *N*_*e*_ have occurred simultaneously, they have also occurred largely independently. That is unfortunate, as CKMR estimates of abundance and related quantities depend on the same key demographic parameters that are needed to calculate *N*_*e*_. Furthermore, genotypes for individuals sampled for CKMR also can be used to generate indirect genetic estimates of effective size, as was recently demonstrated for southern bluefin tuna, *Thunnus maccoyii* (Waples et al. 2018a). Recent work by Akita (2020a,b) has made a good start in this direction for some specialized scenarios, and here we provide a more general treatment. Although POPs-based CKMR has been successfully applied to semelparous species (Rawding et al. 2014), most species are iteroparous with overlapping generations, and that is our focus here.

Bravington et al. (2016b) provide a brief yet authoritative introduction to the statistical underpinnings of CKMR, but not in a format that is very accessible to biologists interested in practical applications. Our synthesis here is motivated by the recognition that (1) a full understanding of the evolutionary ecology of natural populations requires being able to quantify both abundance and effective size; and (2) as a consequence, it is important to maximize the synergistic effects of the two types of analyses. We begin by reviewing basic principles of population demography that are central to both CKMR and *N*_*e*_. We then consider how the same genetic data can be used both to estimate abundance and related quantities via CKMR and to estimate effective size using the popular single-sample estimators. Next, we introduce some simple analytical tools that can be used to allow unbiased, closed-form CKMR estimation of abundance when the number of factors affecting the probability of a close-kin match is limited. Simulated data are then used to model these relatively simple scenarios, and results illustrate how some of the most common covariates (annual mortality; changes in fecundity with age; variance in reproductive success; non-random sampling; intermittent breeding) can affect bias and precision of the estimates and how these factors can be accounted for. We show that the *N*_*e*_/*N* ratio and the probability of a close-kin match both depend on a vector describing the relative probabilities different individuals have of producing offspring. We also show that, although age-specific vital rates are central to both types of analyses, they have different consequences for performance of CKMR estimates than they do for estimates of effective size.

## 2 EVOLUTIONARY DEMOGRAPHY

### 2.1 Vital rates

General features of population demography that underpin models used here are described below (see Table 1 for notation and definitions). The focal population is assumed to be isolated and iteroparous, with separate sexes. Time periods, assumed to be years, are discrete and age is indexed by *x*. Reproduction follows the seasonal birth-pulse model (Caswell 2001). At age *x*, individuals produce on average *b*_*x*_ offspring and then survive to age *x*+1 with probability *s*_*x*_. Both *b*_*x*_ and *s*_*x*_ can differ with age and sex. First reproduction occurs at age α (age at maturity) and maximum age is ω. Often, offspring are not enumerated until they reach a certain age, at which point they are considered “recruits.” We designate *z* as age at recruitment (1≤*z*≤α) and scale *b*_*x*_ to production of offspring that survive to age *z*. Cumulative survival from age *z* through age *x* is 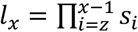, with *l*_z_=1. Letting *N*_*x*_ be the number of individuals of age *x* alive at any given time, common definitions of census size include total abundance, 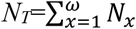 and number of adults, 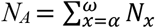. Except as noted we focus on *N*_*A*_.

**Table 1.** Definitions and notation used in this study. Unless specified, variables refer to one time period or year.

|  |  |
| --- | --- |
| $x$ | Age (in years) |
| $z$ | Age at recruitment, when individuals are first inventoried |
| $\alpha$ | Age at maturity |
| $\omega$ | Maximum age |
| $s_x$ | Probability of surviving from age $x$ to age $x+1$ |
| $l_x$ | Cumulative probability of surviving to age $x$ |
| $N_1$ | Cohort size = number of offspring produced per time period that survive to age 1 |
| $N_x$ | Number of individuals of age $x$ alive at any given time; $N_x = N_1 l_x$ |
| $N_T$ | Total number of individuals of all ages alive at any given time; $N_T = \sum N_x$ |
| $N_A (N_f, N_m)$ | Number of adults (females, males) with age $\geq \alpha$ alive at any given time |
| $N_e$ | Effective population size per generation |
| $N_b$ | Effective number of breeders in one year |
| $\hat{N}_A, \hat{N}_e, \hat{N}_b$ | Estimates of adult census size and effective size |
| $n$ | Number of individuals sampled |
| $Y$ | Number of pairwise comparisons to search for close-kin matches |
| $POP$ | Parent-offspring pair |
| $HSP$ | Half-sibling pair |
| $R$ | Number of close-kin matches identified ($R_{POP}$ or $R_{HSP}$) |
| $k_i$ | Number of offspring produced by parent $i$ |
| $\mu_k$ | Mean $k_i$ for all parents |
| $\sigma_k^2$ | Variance of $k_i$ for all parents |
| $b_x$ | Expected number of offspring for parents of age $x$ |
| $\delta_i$ | Difference between expected number of offspring for individual $i$ and the mean for that age and sex ($b_x$) |
| $B_x$ | Total expected number of offspring produced by parents of age $x$: $B_x = N_x b_x$ |
| $\sigma_{k(x)}^2$ | Variance of $k$ for adults of age $x$ |
| $\phi_x$ | Ratio of variance to mean reproductive success for adults of age $x$; $\phi_x = \sigma_{k(x)}^2 / b_x$ |
| $\sigma_{k\bullet}^2$ | Variance in lifetime number of offspring produced by the $N_1$ individuals in a cohort |
| $RO$ | Reproductive output = number of offspring produced. $RO_i = k_i$ for individual $i$ |
| $TRO$ | Total reproductive output of all parents; $E(TRO) = \sum_{x=\alpha}^{\omega} B_x$ |
| $RRO$ | Relative reproductive output of an individual, compared to total reproductive output of all parents; $RRO_i = RO_i / TRO$. |
| $ERRO$ | Expected relative reproductive output, based on a population's vital rates. For an individual of age $x$, $ERRO = E(RO_x) / E(TRO) = b_x / \sum_{x=\alpha}^{\omega} B_x$. |
| $w_i$ | Relative probability that adult $i$ will be the parent of a given offspring |
| $W_i$ | $W_i = w_i / \sum w_i = ERRO_i$ = absolute probability that adult $i$ will be the parent of a given offspring; $\sum W_i = 1$ |
| $CV_W^2$ | Squared coefficient of variation of the parental weights; $CV_W^2 = \sigma_W^2 / \bar{W}^2$, where $\sigma_W^2 = \text{var}(W)$. $CV_W^2$ is the same whether computed using $w$ or $W$. |

### 2.2 Reproduction

Both effective population size and CKMR depend on the population pedigree, which describes genealogical relationships among individuals. Each year, the actual pedigree of a population is one realization of the stochastic process of reproduction and recombination. We use a generalized Wright-Fisher (WF) model of reproduction in which all adults have the potential to breed and produce offspring. Let *k*_*i*_ be the reproductive output (*RO* = number of offspring) produced by the *i*^th^ parent, and let *µ*_*k*_ and 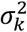 be the mean and variance of *k* across *N*_*A*_ potential parents. In the classical WF model of random reproductive success, each adult has an equal and independent probability of being the parent of each offspring; this produces a multinomial distribution of offspring number, with 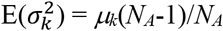 (Crow and Denniston 1988), which is closely approximated by the Poisson variance 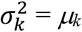. The generalized model (Waples 2020) differs in allowing adults to have unequal probabilities of parenting a given offspring. Define a vector of relative parental weights, ***w***, where *w*_*i*_ is the relative probability that adult *i* will be the parent of a given offspring. These relative weights can be converted to a vector of absolute (standardized) parental weights, ***W***, where *W*_*i*_ = *w*_*i*_/Σ*w*_*i*_ and Σ*W*_*i*_ =1. If all weights are equal, we recover the original WF model, with 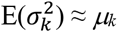; in real populations, however, different individuals generally will have different parentage probabilities, which leads to overdispersed variance in reproductive success 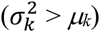 relative to the random expectation. This in turn has important consequences for both *N*_*e*_ and CKMR.

Within a single year of reproduction, two components contribute to variation in individual weights: (1) an among-age effect, which arises from systematic changes in expected fecundity with age; and (2) a within-age effect, which reflects difference in expected reproductive success of individuals of the same age and sex. The among-age effect is determined by the *b*_*x*_ vector from a life table. [As note above, vital rates and other demographic parameters can vary by sex, but to reduce complexity of the notation we omit the subscripts for sex, with the understanding that all analyses involve individuals of a single sex, nominally female.] The within-age effect depends on the relationship between an individual’s expected reproductive success and the *b*_*x*_ value that represents mean fecundity for its age. Thus, the relative parental weight for individual *i* of age *x* can be written as *w*_*i*_ = *b*_*x*_ + *δ*_*i*_, where *δ*_*i*_ is the deviation from the age-specific mean. A positive *δ* means that the individual is expected to be above average for its age; a negative *δ* indicates the reverse. If all *δ*_*i*_=0, the within-age effect disappears, expected age-specific variance in reproductive success equals the mean 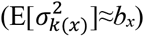, and individuals of the same age effectively behaves like a mini Wright-Fisher population.

Empirical data for 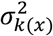 are rarely published, but there are many reasons why individuals of the same age and sex might not have identical expectations of reproductive success. Some might have established breeding territories or acquired harems and others not; some might have phenotypes that favor them in sexual selection. In many species, fecundity depends more on size than age, in which case variation in *w*_*i*_ for same-age individuals will be correlated with variance in size-at-age, leading to (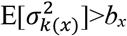. (see Waples et al. 2018a for an example involving southern bluefin tuna). The magnitude of the within-age effect is quantified by the age-specific index 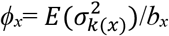, where 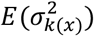 and *b*_*x*_ have been scaled to their expected values in a stable population (Waples et al. 2011; Waples 2016).

Finally, in iteroparous species individual weights can vary across years, for two major reasons: (1) if *b*_*x*_ varies with age; (2) if an individual’s condition factor changes, which in many species depends on whether it has reproduced in a recent year (see Section 3.2.2).

### 2.3 Effective population size

#### 2.3.1 Demographic factors that affect *N*_*e*_

Close-kin methods provide information about abundance in the recent past, so contemporary (rather than long-term) effective size is our focus. Mean and variance in offspring number are the major factors that determine effective size. For seasonal reproduction in an age-structured species, the most relevant effective-size metric is the inbreeding effective number of breeders per year (*N*_*b*_) because it relates directly to the annual number of adults (*N*_*A*_), which also is generally the focus of CKMR analyses. *N*_*b*_ can be calculated using the standard discrete-generation formula for inbreeding effective size (Crow and Denniston 1988; Caballero 1994):

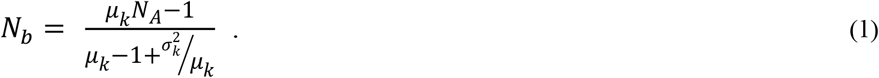

With separate sexes, *N*_*b*_ can be calculated separately for males and females and an overall effective size obtained using Wright’s (1938) sex-ratio adjustment. For simplicity we focus on census size and effective size for females (*N*_*f*_, *N*_*b(f)*_), but exactly analogous equations apply to males.

Equation 1 is exact when based on realized population parameters; it can be reformulated as an expectation using the parental weights described above. Let 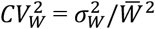 be the squared coefficient of variation (CV) of the individual weights (which is the same whether calculated using raw or standardized weights). Then, the expectation for the effective number of female breeders per year is just (Waples 2020):

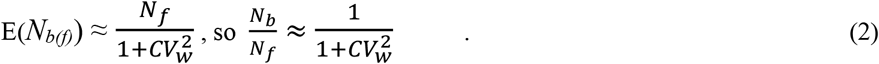

Although *N*_*b*_ relates directly to annual abundance estimated by CKMR, virtually all evolutionary theory depends on effective size per generation (*N*_*e*_) rather than *N*_*b*_ per year. Calculating *N*_*e*_ in iteroparous species is complicated by the necessity of integrating information across multiple episodes of reproduction. All else being equal, *N*_*e*_ per generation is positively correlated with generation length and negatively correlated with lifetime variance in number of offspring, 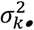 (Hill 1972). The program *AgeNe* (Waples et al. 2011) calculates 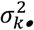 using the species’ age-specific vital rates (*s*_*x*_, *b*_*x*_, *ϕ*_*x*_), under the assumptions (adopted from Hill 1972 and Felsenstein 1971) that vital rates and population size are constant and survival and reproduction are independent over time. Demographic *N*_*e*_ also can be converted from an estimate of *N*_*b*_ based on the *N*_*b*_/*N*_*e*_ ratio, which largely depends on 2-3 life history traits (Waples et al. 2013, 2014).

#### 2.3.2 Genetic methods for estimating *N*_*e*_

The sibship method developed by Wang (2009) considers three types of relationship in the one-generation pedigree: full sibling pairs (FSPs), half sibling pairs (HSPs), and unrelated (U), which nevertheless can be related through previous generations. In a random sample of *n* progeny, *Y* = *n*(*n*-1)/2 ≈ *n*^2^/2 pairwise comparisons are possible, and expected proportions of siblings are simple functions of *N*_*e*_ (from Wang 2009, ignoring the term for selfing):

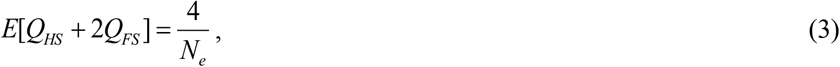

where *Q*_*HS*_ is the fraction of pairs that are half siblings (maternal and paternal HSPs combined) and *Q*_*FS*_ is the fraction that are full siblings. Substituting for *Q* = *R*/*Y*, where *R* is the number of sibling matches, and rearrangement leads to the estimator of effective size:

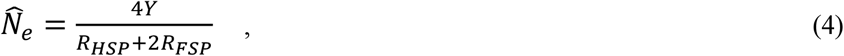

which has the number of pairwise comparisons in the numerator and the total number of sibling doses (one for each HSP and two for each FSP) in the denominator. When applied to offspring from a single cohort, Equation 4 provides an estimate of *N*_*b*_. Wang et al. (2010) described an extension of this sibship method to estimate *N*_*e*_ when generations overlap, but their model assumes that *ϕ*_*x*_ = 1 for each age and sex so is of somewhat limited general applicability.

Linkage disequilibrium (LD) quantifies associations of alleles at different gene loci. Random LD is generated every generation by mating among a finite number of parents, and that provides the basis for estimating effective size. Loci on the same chromosome (“linked”) provide information about historical *N*_*e*_, so unlinked markers are most suitable for estimating contemporary *N*_*e*_. For unlinked loci, *N*_*e*_ can be estimated from *r*^2^ (a measure of LD at pairs of loci) and sample size (*n* = number of individuals) (Hill 1981):

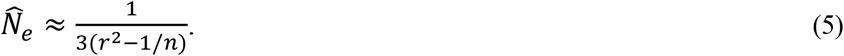

The program LDNe (Waples and Do 2008) adjusts for bias in Equation 5 caused by ignoring second-order terms in 1/*n* and 1/*N*_*e*_.

When applied to offspring from a single cohort, the LD method primarily reflects the signal from parental *N*_*b*_, with some influence from *N*_*e*_ in previous generations (Waples 2005; Waples et al. 2014). When applied to mixed-age samples, LDNe generally underestimates generational *N*_*e*_ by 10-40% (Waples et al. 2014). Blower et al. (2019) developed simulation software for age-structured species that can help researchers interested in using the LD method to estimate effective size to determine sample sizes of individuals and loci that should produce desired levels of precision.

## 3 MARK-RECAPTURE METHODS

Traditional mark-recapture (MR) involves three elements:

- Capture and mark a number (*n*_*1*_) of individuals;
- Release them to mix with the general population;
- Collect a second sample of *n*_*2*_ individuals; those marked are the “recaptures” (*R*).

The fraction of original marks that are recaptured provides information about population abundance, but this metric also can be affected by a variety of other factors, including behavioral effects on sampling probabilities, tag loss, mortality, immigration/emigration, and incomplete mixing between samples. Accordingly, a complex statistical framework for conducting MR analyses has been developed to account for these factors (Lebreton et al. 1992; Pollock 2000; Lindberg 2012). Commonly, MR analyses are conducted using Integrated Population Models (IPMs) that combine information from multiple datasets within a single pseudo-likelihood framework to estimate population abundance, trends, and vital rates. IPMs have been widely used in fisheries stock assessments for several decades (Fournier and Archibald 1982; Punt et al. 2020) and more recently have been applied to wildlife management (Arnold et al. 2018).

In CKMR, close-kin matches are analogous to MR recoveries. Full siblings and parent-offspring pairs on average share half their genes, and half siblings share one quarter. As with MR, close-kin matches provide information about abundance but also are affected by a variety of other factors, which include (but are not limited to) age-specific survival and fecundity, spatial structure, sampling selectivity, variance in offspring number, and correlations of individual reproductive success over time. Most CKMR applications are complicated enough that they are best conducted by building a log-pseudo-likelihood framework, analogous to the IPMs used with MR, that incorporates outcomes of each pairwise comparison (Bravington et al. 2016a,b; Hillary et al. 2018; Conn et al. 2020).

Estimation frameworks for MR and CKMR can be quite complicated and are largely opaque to all but the most dedicated aficionados. In this synthesis, for heuristic purposes we take a different approach. First, by focusing on the simplest versions of the models, we illustrate fundamental similarities between MR and CKMR methods to estimate abundance and sibling-based methods to estimate effective size, and we show how to account for some common factors that can affect the estimates. Second, we suggest a new way to formulate the probability of a sibling match that can be useful in illustrating the influence of various covariates. Finally, we use simulated data to illustrate how factors like age-specific changes in vital rates, adult mortality, intermittent breeding, within-age variance in reproductive success, and experimental design can affect performance of abundance and effective size estimators.

### 3.1 Traditional mark-recapture estimation

In the simplest “cartoon” version of MR (if the probability of a recovery is only influenced by population size), the estimate of abundance is simply 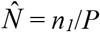, where *P* = *R*/*n*_*2*_ is the fraction of the second sample that is marked. This leads to the abundance estimator

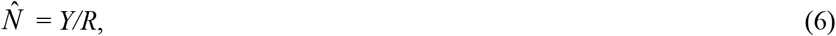

where *Y* = *n*_*1*_*n*_*2*_ is the number of potential comparisons of individuals in the two samples, and the denominator is the number of matches.

### 3.2 Close-kin mark-recapture estimation

In the cartoon version of CKMR, all parents are equivalent in terms of producing offspring and all *Y* pairwise comparisons to search for close kin are independent. Under these conditions, the probability that a random pair of individuals will produce a close-kin match is *P*_*match*_ = 1/*N*_*f*_, and the expected number of matches is just the product of the number of comparisons and the probability of success: (E(*R*) = *P*_*match*_\**Y* = *Y*/*N*_*f*_. This leads to the naive estimator of adult abundance as 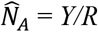, which is identical in form to Equation 6 for MR and Equation 4 for sibship estimation of *N*_*e*_.

More generally, estimation of abundance using the incidence of close-kin matches is based on two basic principles:

A. It is possible to specify the probability that a random pair of individuals will produce a close-kin match, and this probability is inversely related to adult abundance, such that *P*_*match*_ = *f*_*1*_ (1/*N,Z*) = *f*_*2*_(*Z*)/*N*, where *f*_*1*_ and *f*_*2*_ are functions and *Z* represents other covariates than also can affect close-kin probabilities.
B. Pairwise comparisons to search for a match are independent, so the expected number of matches (*R*) is the product of the number of comparisons (*Y*) and the probability that each produces a match: E(*R*) = *Y*\**P*_*match*_ = *Y*\**f*_*2*_(*Z*)/*N*.

Principle B is not generally true in the strict sense, at least for POPs (if A is the mother of X, B cannot be); however, except in very small populations, sampling generally will be sparse enough that this assumption does not lead to appreciable errors (Bravington et al. 2016b; see also Section S2.2 in Supporting Information). Principle A is reasonable if one has a flexible enough concept of ‘function.’ If A and B hold to an acceptable degree, then rearrangement produces

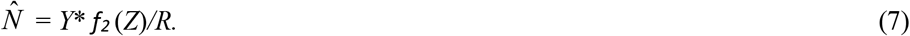

Because *Y* and *R* are empirically determined, estimation of *N* is possible provided one can define the function *f*_*2*_ so that all relevant covariates are accounted for. In certain cases, Equation 7 can be expressed as 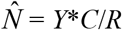;, i.e., the function *f*_*2*_ (*Z*) takes the simple form of a scalar conversion factor *C*, which can be calculated as a function of individual parental weights *W* (see section 2.2) and other life-history parameters. Supporting Information S2 illustrates how to calculate *C* in a number of cases.

The core problem for CKMR, therefore, is to account for all other factors besides *N* that can affect the probability of close-kin matches. Among the most important covariates to consider in calculating probabilities of a close-kin match are mortality, fecundity, sampling selectivity, temporal correlations, population structure, age, and sex. Unless the sex ratio is even and vital rates are the same in both sexes, the distribution of genetic “marks” passed on to their offspring by male and female parents will be different. In general, therefore, it is best to make sex-specific CKMR estimates, for which mtDNA can be useful (see Section 5.3).

#### 3.2.1 Parent-offspring pairs

Notwithstanding the difficulty in developing a general closed formula for CKMR, one key metric is central to all CKMR analyses because it describes the probability that any given pair of individuals will produce a close-kin match: the relative reproductive output (*RRO*) of different potential parents. For POPs, consider the probability [P(*i*→*j*)] that female *i* is the mother of offspring *j*. A general expression for this probability is (Bravington et al. 2016b):

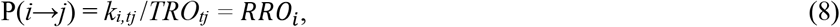

where *k*_*i,tj*_ = *RO*_*i,tj*_ is the number of offspring produced by female *i* in the year of *j*’s birth and *TRO*_*tj*_ = Σ*k*_*i,tj*_ is the total number of offspring by all females in that year. To estimate abundance, it is necessary to combine these probabilities across all sampled offspring and potential parents. POPs provide information about *N*_*f*_ in the year the offspring were born, so if samples are taken from multiple cohorts, a time-series of abundance estimates can be obtained via age-structured population dynamics, as in many fish stock assessments (Quinn and Deriso 1999).

Because actual reproductive output is rarely known, CKMR generally deals with expected relative reproductive output, *E*(*RRO*), based on expectations for different classes of parents (Bravington et al. 2016b). This concept provides a direct link to the parental weighting scheme described previously: parental weights for a given individual *i* are calculated as *W*_*i*_=E(*RO*_*i*_)/E(*TRO*), so E(*RRO*_*i*_) = *W*_*i*_.

CKMR estimates based on POPs are greatly simplified by the fact that each offspring has exactly one mother. This means that variation in offspring number does not, by itself, affect expected number of matches (but it can if there is unmodeled correlation between reproductive output and sampling probability; see Section 5.2.1).

In the simplest POP-based examples, *P*_*POP*_ = 1/*N*_*f*_, which would lead to an estimator of adult female census size, directly analogous to a standard Lincoln-Petersen MR estimate (Seber 1982), as

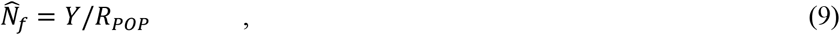

which has the same general form as Equations 4 and 6. Because E(*R*_*POP*_) is not sensitive to variation in offspring number, CKMR estimates based on POPs are not confounded by a signal related to *N*_*e*_ or *N*_*b*_. However, as illustrated in Section 5.2.2, precision of POPs-based estimates *is* inversely related to variation in reproductive success. If one or a few parents dominate reproduction, the number of matches will vary widely depending on whether the prolific parent(s) are sampled.

#### 3.2.2 Siblings

CKMR for siblings is considerably more complicated because (1) parents are not observed directly, and (2) the probability of a sibling match depends heavily on adult mortality and the extent to which reproductive output varies both within and across years. As demonstrated below, only cross-cohort sibling comparisons are suitable for estimating abundance. Because full siblings from different cohorts will be rare in large, randomly mating populations, CKMR analyses generally focus on finding HSPs, and the probability that two offspring share a mother can be taken as a close approximation to the probability that they are a maternal HSP (MHSP). Our examples focus on MHSPs, but exactly equivalent equations apply to paternal HSPs.

Realistic CKMR applications based on siblings must consider two ordered time periods, 0 and *t*. For two samples of *n*_*1*_ and *n*_*2*_ offspring, *Y* = *n*_*1*_*n*_*2*_ across-sample comparisons are possible. A general formula for the probability that two randomly selected offspring, one from time 0 and one from time *t*, have the same female parent is (from Equation 3.9 in Bravington et al. 2016b):

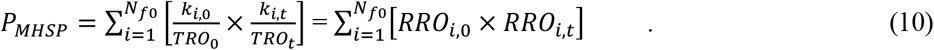

It is necessary to sum across all possible females that could have been alive when potential siblings were born. Females that mature after time 0 cannot produce a sibling match for offspring born in time 0, so the summation is Equation 10 is taken over the *N*_*f* 0_ females that were sexually mature at time 0.

Equation 10 does not generally lead to a closed-form estimator of adult abundance, unless the species’ biology allows a number of simplifying assumptions (see Hillary et al. 2018 for an example involving the white shark, *Carcharodon carcharias*, which also required estimating adult mortality from the data). However, some heuristic insights into factors that influence *P*_*MHSP*_ can be achieved if the equation is reformulated as follows (see Supporting Information Section S1 for details):

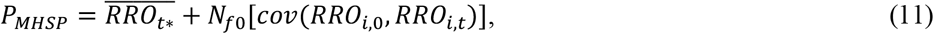

where the “*” in *RRO*_*t\**_ indicates that this vector only includes data for the subset of time-*t* females that were mature at time 0. It might appear from Equation 11 that the term *N*_*f*0_ represents a signal of female population size at time 0, but that is misleading: the denominator of the population covariance is *N*_*f* 0_, so that signal cancels out. Instead, the term 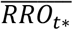 provides information about adult abundance at time *t*. Two general cases can be identified.

##### 3.2.2.1 Case 1. Offspring are from the same cohort (*t* = 0)

If only a single time period is involved, *RRO*_*i,0*_ = *RRO*_*i,t*_ = *RRO*_*i*_, with 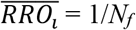. Noting that the covariance of a random variable with itself is its variance, Equation 11 becomes

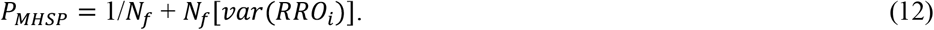

Equation 12 shows that, when offspring from the same cohort are compared, the probability of a sibling match increases as a function of the realized variance in offspring number. This variance is the primary signal for *N*_*b*_, so siblings from the same cohort provide information related to effective size and not directly about adult abundance. For this reason, CKMR estimates using siblings generally attempt to exclude within-cohort comparisons (Bravington et al. 2016b; Hillary et al. 2018; Thomson et al. 2020).

##### 3.2.2.2 Case 2. Offspring are from different cohorts (*t* ≠ 0)

In comparing offspring from different cohorts, it is always important to account for adult mortality: if an adult female dies between times 0 and *t*, it can’t be a parent in time *t*, so *RRO*_*i,t*_ = 0. These null parents are included in the summation in Equation 10 and will reduce 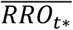 and therefore *P*_*MHSP*_ unless this mortality is accounted for. Other common factors that affect sibling probabilities are considered below.

###### Changes in fecundity with age

All adults that survived to reproduce again at time *t* will be at least one year older. If *b*_*x*_ changes with age (fecundity increases with age in many species, especially fish and other poikilotherms with indeterminate growth), this will affect 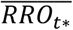 (and perhaps the covariance term as well), unless this effect is accounted for.

###### Persistent individual differences

If within-age individual deviations from mean fecundity (the *δ*_*i*_ defined in Section 2.2) are non-zero and positively correlated over time, such that some individuals are consistently above or below average at producing offspring, this will (all else being equal) lead to positive *cov*(*RRO*_*i*,0_, *RRO*_*i,t*_) and a higher proportion of siblings, as well as reduced *N*_*e*_ (Lee et al. 2011, 2020). Body size is one factor that can easily lead to positive temporal correlations in reproductive success. Fecundity often depends on size more directly than on age, and an individual that is large (or small) for its age at time 1 is also likely to differ in a similar way in subsequent years. Whether an individual is ‘sexy’ or has secured a territory are other factors that could produce persistent individual differences.

###### Skip breeding

Skip breeding leads to a complex pattern of *cov*(*RRO*_*i*,0_, *RRO*_*i,t*_) that can vary both in magnitude and sign, depending on the species’ life history and the time interval for cross-cohort comparisons (Figure 1). For example, consider a species for which females generally skip *t* years after reproducing before doing so again. Comparisons of offspring born <*t* years apart will produce few if any sibling matches, while comparisons exactly *t* years apart will produce sibling matches at a rate determined not by female population size as a whole, but by the ≈ *N*_*f*_ /*t* females that actually reproduce each year. It should be possible to develop an adjustment for skip breeding using the vector (*θ*_*t*_) of the probability that an individual will reproduce in the current year, given that it last reproduced *t* years before (Shaw and Levin 2013). For female loggerhead turtles (*Caretta caretta*), *θ*_*t*_ = [0.025, 0.443, 0.634, 0.743, 1] for *t*=1,5 (Waples and Antao 2014), indicating a substantial effect of prior reproduction lasting four years. With adequate data, CKMR can estimate parameters like *θ*_*t*_ based on age-gap patterns in the data (see Section 6.1).

**Figure 1.**
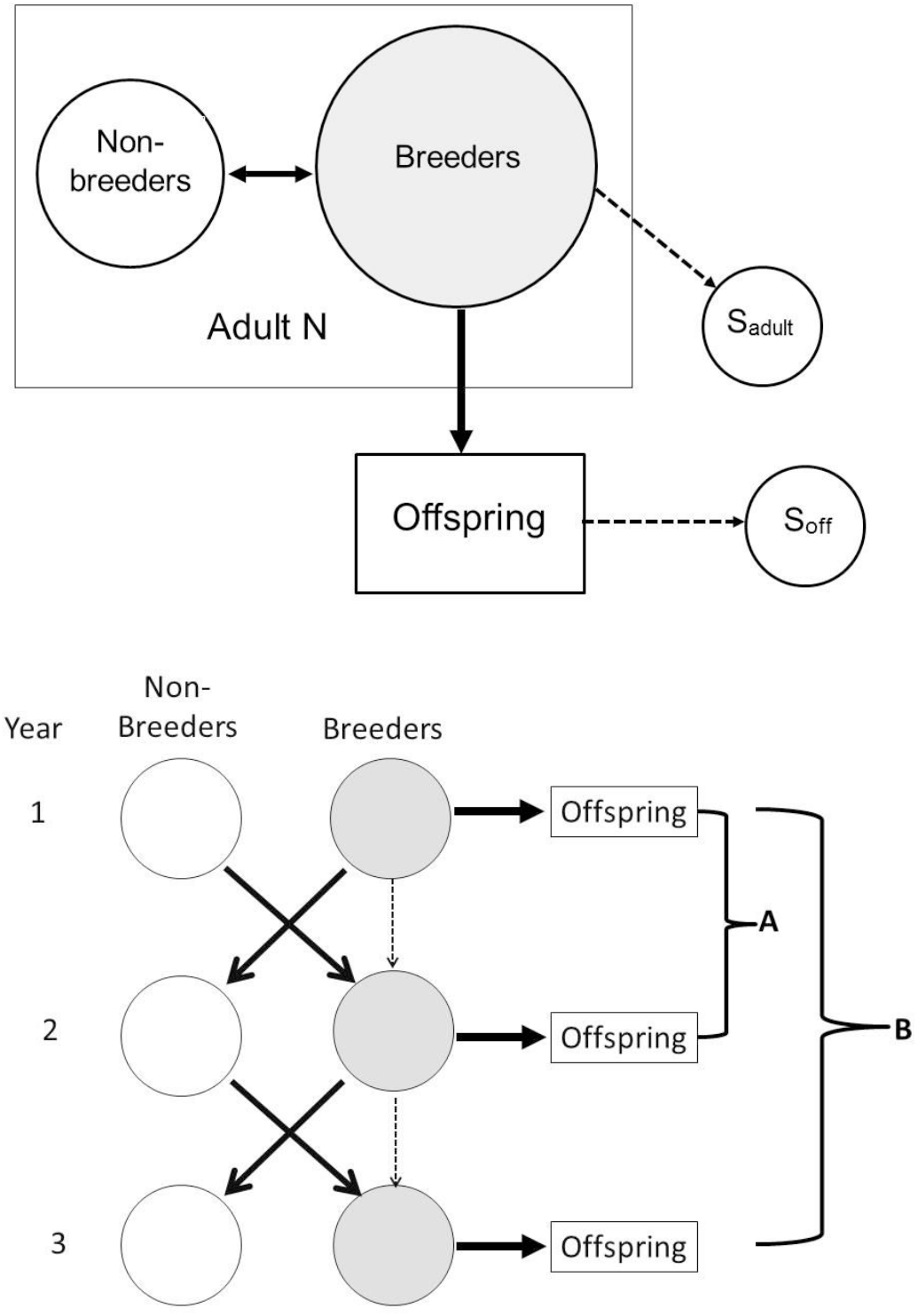
Schematic diagram showing possible effects of skip breeding on CKMR estimates. Top: If non-breeders are not sampled, POPs will provide an abundance estimate only for the breeding population. Bottom: A multi-year study for a population in which most adults alternate between being breeders and non-breeders; the thin dashed arrows indicate that a fraction of adults breed in consecutive years. Two consecutive years of POPs collections will sample the entire adult population. Few if any HSPs will be found for pairs of individuals separated in age by odd numbers of years (sampling scheme A), while the number of HSPs for comparisons of individuals born an even number of years apart (sampling scheme B) will reflect the number of breeders per year more than the adult census size, *N*_*A*_. Genetic methods that estimate *N*_*b*_ based on single-cohort samples also would produce an estimate related to the number of actual breeders each year, rather than the total number of adults.

## 4 SPATIAL STRUCTURE and SAMPLING

Spatial structure, where reproduction and/or sampling is concentrated in certain geographic locations, will *not* bias CKMR estimates if either of the following conditions are met (Conn et al. 2020): (1) offspring (siblings analyses) or parents and offspring (POPs analyses) mix thoroughly before sampling occurs; or (2) sampling is equiprobable, meaning that every individual has an equal chance of being sampled. To illustrate the first criterion, if reproduction is panmictic in area A, where adults are sampled, and offspring randomly migrate to areas B and C, sampling in just one of those areas still produces a random sample of all offspring (see Bravington et al. 2016a for an empirical example involving southern bluefin tuna). If the second criterion (and other CKMR assumptions) are met, abundance estimates will reflect the total number of adults. If spatially-restricted reproduction is persistent and there are independent or semi-independent populations or stocks for which separate abundance estimates are desired, the naive CKMR estimate of a single overall abundance might be unsatisfactory. If estimates are available for key parameters describing the degree of reproductive connectivity and distributional overlap, it might be possible to develop area-specific abundance estimates (Davies et al. 2017). Fortunately, if population stratification is strong enough to substantially affect CKMR estimates, this should be apparent from even a fairly coarse preliminary sampling design (Conn et al. 2020). Furthermore, spatial patterns in distribution of close-kin can provide insights into population subdivision and connectivity (Økland et al. 2009; Palsbøll et al. 2010; Wang 2014; Feutry et al. 2017).

In general, the most important proviso regarding sampling is that whether an individual is sampled should have no effect on the probability that one of its close kin is also sampled. If this criterion is satisfied, conditioned on covariates associated with the sampled individuals, then expected values of *RRO* are multiplicative, which makes the CKMR analyses tractable. Joint sampling of close kin in the same spatio-temporal stratum at higher frequencies than would occur by chance will increase the probability of close-kin matches; this potential source of bias can be reduced by appropriate filtering of the data (as Thomson et al. 2020 did with the school shark, *Galeorhinus galeus*).

Sampling considerations for single-sample genetic estimators are similar to those for sib-based CKMR, except that non-random collection of siblings can be difficult to avoid when sampling within cohorts. Without independent information, it is not possible to reliably compensate for a non-random sample by excluding some or all putative siblings, although some approaches can reduce bias (Waples and Anderson 2017). Biases to genetic estimates of *N*_*b*_ from undetected population structure should be similar to those for CKMR-based estimates of abundance. If the focal population is part of a metapopulation but sampling is local, the LD method primarily estimates local *N*_*b*_ unless migration is very high in genetic terms (>5-10%; Waples and England 2011; Gilbert and Whitlock 2015).

## 5 WORKED EXAMPLES

### 5.1 Simulations

We simulated demographic and genetic data (using the software CKMRPop, Anderson 2021) to illustrate effects of some common factors on CKMR and *N*_*e*_ estimators. Scenarios modeled here were simple enough that the function *f*_*2*_ from Equation 7 could be specified and CKMR estimates could be made using that closed-form equation. Below we briefly describe the model and summarize the main results and conclusions. More details about the simulations and explanations for all the calculations can be found in Supporting Information.

The core scenario involved a single closed population with the following features (notation follows that introduced in Section 2 and defined in Table 1):

- age at maturity (α) = 3
- maximum age (ω) = 10
- equal primary sex ratio and same vital rates in males and females
- annual probability of survival was constant at *s*_*x*_ = 0.7
- relative fecundity was either constant with age (*b*_*x*_ = 1 for all adults) or proportional to age (*b*_*x*_ = *x* for all adults)
- The *N*_*A*_ adults produced a constant number of yearling offspring per year (*N*_*1*_ = cohort size = 2000)
- number of offspring produced by individuals of the same age and sex was drawn from a negative binomial distribution with either Poisson (∼random) variance (*ϕ* = 1) or overdispersed variance (*ϕ* = 10).

The life table for this core population is shown in Table 2. With constant cohort size and random survival, total abundance each year averages 3239 individuals of each sex (*N*_*T*_ = 6478), of which 3400 are juveniles and *N*_*A*_ = 3078 are adults (hence *N*_*f*_ = *N*_*m*_ = 1539). Table 3 defines six Scenarios (A-F) that represent variations on this life table for which we simulated data, in addition to some other scenarios described below.

**Table 2.**
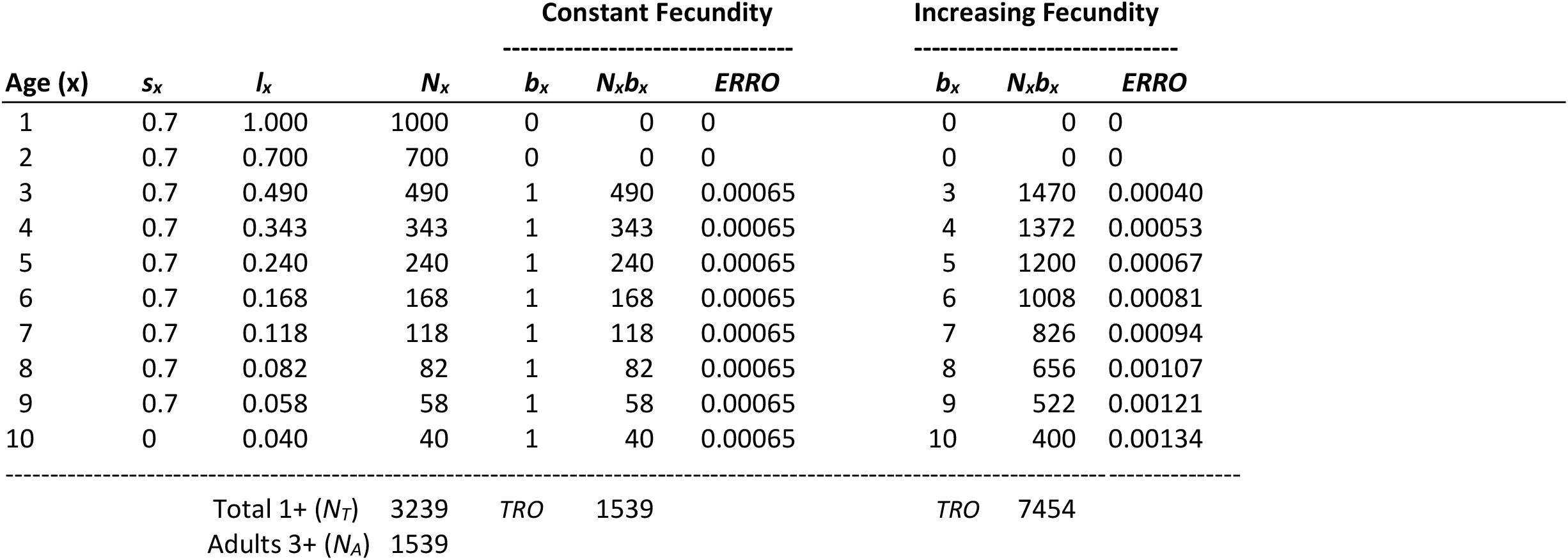
Life table for a hypothetical species used in the simulations. s_*x*_ is the probability of surviving from age *x* to *x*+1, *l*_*x*_ is cumulative survival through age *x*, and *b*_*x*_ is relative fecundity of an individual of age *x*. This example assumes an equal sex ratio and the same vital rates in males and females, but data are shown for only one sex (nominally females). The hypothetical population produces 2000 offspring in each cohort, of which *N*_*1*_ = 1000 are females. Expected numbers in successive age classes are defined by *N*_*x*_ = *l*_*x*_*N*_*1*_. Two fecundity scenarios are considered: constant fecundity and fecundity increasing with age. *TRO* = Σ*N*_*x*_*b*_*x*_ is total reproductive output of the population in one time period, in the same relative units as *b*_*x*_, and *ERRO* is the expected relative reproductive output of an individual of the specified age (all individuals of the same age are assumed to have the same *ERRO*). *ERRO* is identical to the standardized parental weights described in Section 2.2.

**Table 3.** Comparison of expected and estimated parameters for six scenarios evaluated in the simulations. Except for scenario F, for which cohort size and adult *N* are 10 times as large, all scenarios have a constant cohort size of *N*_*1*_ = 2000 newborns, which produces adult *N*_*A*_ = 3078 (1539 of each sex; Table 2). *n*_*off*_ and *n*_*parents*_ are annual sample sizes of offspring and potential parents, respectively. In selective sampling (Scenario D), the relative probability that an adult was sampled was equal to its expected fecundity (*b*_*x*_); in other scenarios, adult sampling was random. Samples of potential parents were drawn only from mature (age 3+) females except in Scenario E, where the sample was collected from the entire population. Sampling of adults has no effect on siblings, so results for Scenarios D and E are only shown for POPs. Exp(*R*) and Obs(*R*) are expected and observed numbers of close-kin matches, respectively, and the conversion factor *C* is the constant that relates the true probability of a match to the naive value 1/*N*_*f*_ (see Eq. S3). For sibship analyses there is a separate *C* value for each age gap between offspring, and values shown here are weighted mean 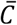 values applicable to the total number of comparisons. Raw (naïve) 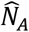 is the estimate from Equation 7, assuming all *P*_*match*_ = 1/*N*_*f*_ ; the adjusted estimator 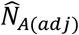 accounts for effects of mortality, changes in fecundity with age, and sampling using the conversion factor *C*.

|  | Scenario | A | B | C | D | E | F |
| --- | --- | --- | --- | --- | --- | --- | --- |
| | $N_I$ | 2000 | 2000 | 2000 | 2000 | 2000 | **20000** |
|  | fecundity | constant | constant | **increasing** | **increasing** | constant | constant |
|  | adult sampling | random | random | random | **selective** | **random<sup>a</sup>** | random |
| | $\phi$ | 1 | **10** | 1 | 1 | 1 | 1 |
| | $N_A$ | 3078 | 3078 | 3078 | 3078 | 3078 | **30783** |
| **POPs** | $n_{off}$ | 300 | 300 | 300 | 300 | 300 | 600 |
| | $n_{parents}$ | 300 | 300 | 300 | 300 | 300 | 600 |
| | $Exp(R)$ | 58.5 | 58.5 | 58.5 | 67.5 | 27.9 | 23.4 |
| | $Obs(R)$ | 59.2 | 59.2 | 58.9 | 66.9 | 27.8 | 23.1 |
| | $C$ | 1.00 | 1.00 | 1.00 | 1.15 | 0.48 | 1.00 |
| | $\hat{N}_A$ raw | 3072 | 3182 | 3135 | 2802 | 6722 | 32556 |
| | $\hat{N}_{A(adj)}$ | 3072 | 3182 | 3135 | 3094 | 3201 | 32556 |
| **Siblings** | $n_{off}$ | 100 | 100 | 100 | - | - | 300 |
| | $Exp(R)$ | 63.7 | 63.7 | 84.3 | - | - | 57.4 |
| | $Obs(R)$ | 63.8 | 67.7 | 82.4 | - | - | 58.8 |
| | $\bar{C}$ | 0.4905 | 0.4905 | 0.648 | | | 0.4905 |
| | $\hat{N}_A$ raw | 6504 | 6577 | 4854 | - | - | 61348 |
| | $\hat{N}_{A(adj)}$ | 3122 | 3157 | 3145 | - | - | 29447 |
|  | **True $N_b$** | 3078 | 388 | 2665 | - | - | 30783 |
| | $\hat{N}_b$ | 3074 | 390 | 2658 | - | - | 28805 |

<sup>a</sup>In Scenario E, sampling of potential parents was random but with respect to the entire population rather than just mature adults

CKMRPop does not do kin-finding based on genetic data. Because we wanted to focus on bias and precision related to experimental design and the species’ life history, we used the true pedigree recorded by CKMRPop to identify POPs and siblings, which is equivalent to assuming that all close-kin matches were made without error.

### 5.2 Results

Bias can arise in CKMR estimates when the model does not fully account for all factors that affect the probability of a close-kin match (*P*_*match*_). In the examples considered here, we accounted for potential biases by calculating a conversion factor *C* that is the ratio of the true *P*_*match*_ to the naïve value of 1/*N*_*f*_. This conversion factor thus plays the role of the function *f*_*2*_ in Equation 7. The value of *C* for any given application is determined by the focal species’ vital rates and the experimental design, but it is independent of population size (see Supporting Information for details about calculation of *C*). *C* can then be used to accurately predict the expected number of close-kin matches and to obtain an unbiased estimate of abundance.

#### 5.2.1 Bias

##### 5.2.1.1 Parent-offspring pairs

In our simulations, POP-based estimates of adult *N* were unbiased using the naive model that assumes *P*_*POP*_ = 1/*N*_*f*_, (Table 3 and Figure 2) except when:

**Figure 2.**
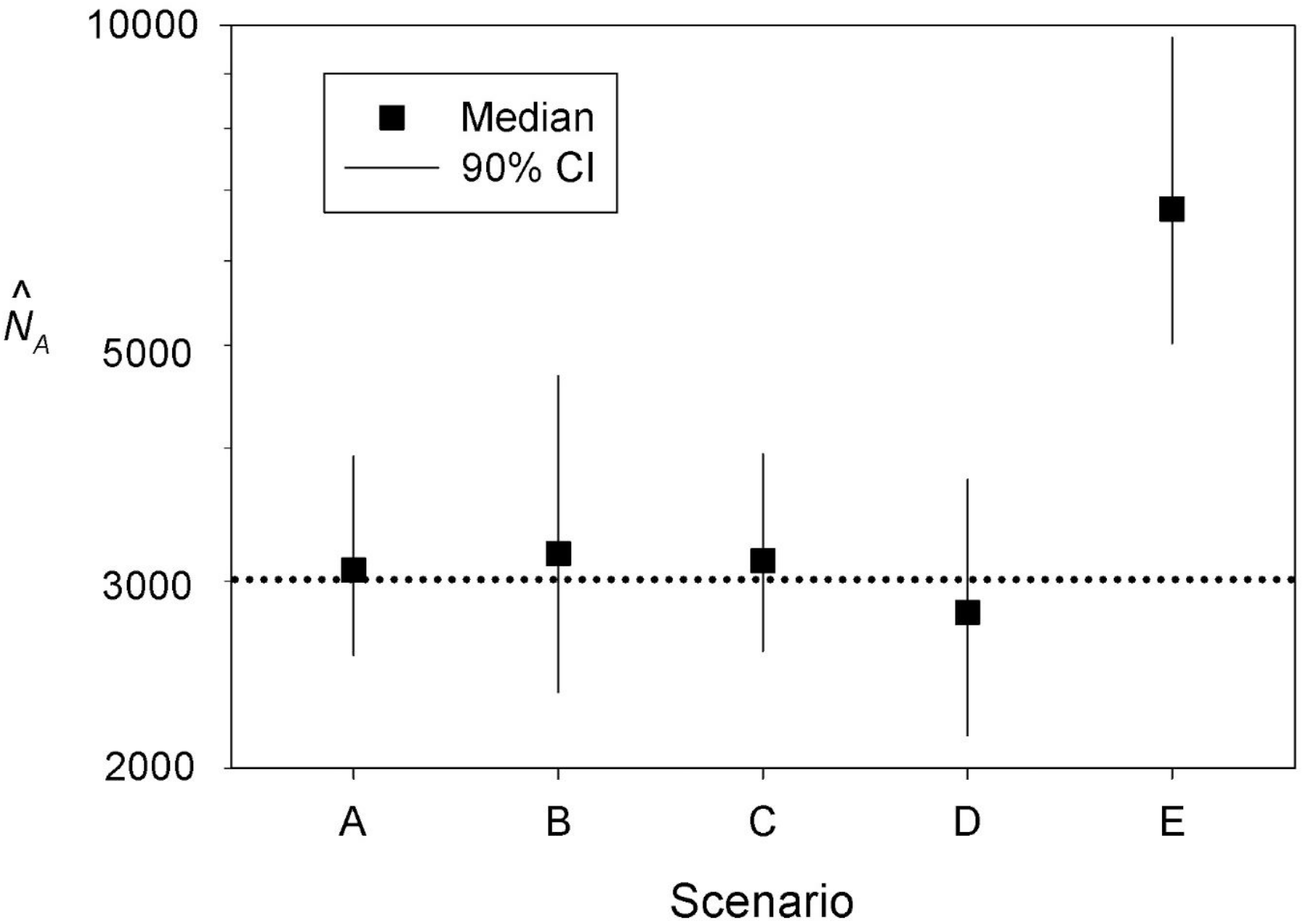
POPs results for simulated scenarios A-E from Table 2. Symbols show medians of 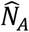 across 500 replicate simulated samples; vertical lines show empirical confidence intervals (CIs) defined by the upper 95% and lower 5% quantiles. All scenarios have the same true adult *N*_*A*_ = 3078 (dotted line). Estimated *N*_*A*_ (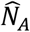: *Y* axis) is the raw estimate based on the observed number of POPs and assuming that *P*_*POP*_ = 1/*N*_*f*_ ; all estimates adjusted using the conversion factor *C* to account for deviations from this assumption were close to true *N*_*A*_ (see Table 3). Note the log scale on the *Y* axis.

1. Fecundity and probability of being sampled both increased with age; or
2. The sample of potential adults also included immature individuals.

In situation 1), joint effects of age-specific changes in selectivity and fecundity increased the number of POP matches beyond the number expected under the naive model, leading to *C*>1 (calculations shown in Table S1). In situation 2), dilution of the ‘adult’ sample by inclusion of juveniles that could not produce offspring reduced the fraction of comparisons that produced a POP match, leading to *C*<1. With equiprobable sampling of the entire population, this produced an unbiased estimate of overall abundance (*N*_*T*_) but would represent a bias if the goal were to estimate *N*_*A*_. After adjusting *P*_*POP*_ by the factor *C*, 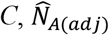 for these two scenarios was unbiased as well (Table 3).

##### 5.2.1.2 Siblings

Because sibling-based estimates of abundance depend on cross-cohort comparisons of offspring to search for a shared parent, in general one of the following conditions must be met for CKMR to be feasible: a) mixed-age samples can be aged precisely and sorted into cohorts, or b) a series of samples from individual cohorts is collected over time. Our simulations followed the second approach; in each replicate for sibling analyses we collected samples of yearlings in 5 consecutive years, which allowed us to compare cohorts born 1-4 years apart (the ‘age gap’). Even when all adults are equally likely to produce offspring, it cannot be the case that the probability of a sibling match is 1/*N*_*f*_, because some parents will die and cannot be the parent of an offspring born in a later year. If all adults are otherwise equally likely to produce offspring, the conversion factor *C* can be calculated from cumulative survival rates over time. If fecundity changes with age, the bias adjustment is more complicated but can be done using Equation 10 to calculate *P*_*MHSP*_ based on the population’s vital rates (as illustrated in Table S2). By making these adjustments, unbiased abundance estimates can be obtained for a wide range of scenarios, for very small to very large populations (Table 3 and Figure S1).

Figure 3 illustrates the effects of intermittent breeding on the number of sibling matches as a function of the age gap. Allowing a random subset of adults to reproduce every year does not affect the total number of cross-cohort comparisons (columns ‘Total 1-4’), but it does reduce *N*_*b*_ per year and hence increases the number of within-cohort siblings. In scenarios where the probability that the same individual reproduces in consecutive years is low or 0 (as in the ‘Alternate’ scenario in Figure 3), the expected number of sibling matches can vary dramatically depending on the age gap—a factor that must be considered to obtain an unbiased estimate of abundance. Fortunately, the empirical distribution of age gaps between sibling matches (or between POP matches in a long-term POP study; see Bravington et al. 2016a) can be used to estimate the nature and magnitude of skip breeding.

**Figure 3.**
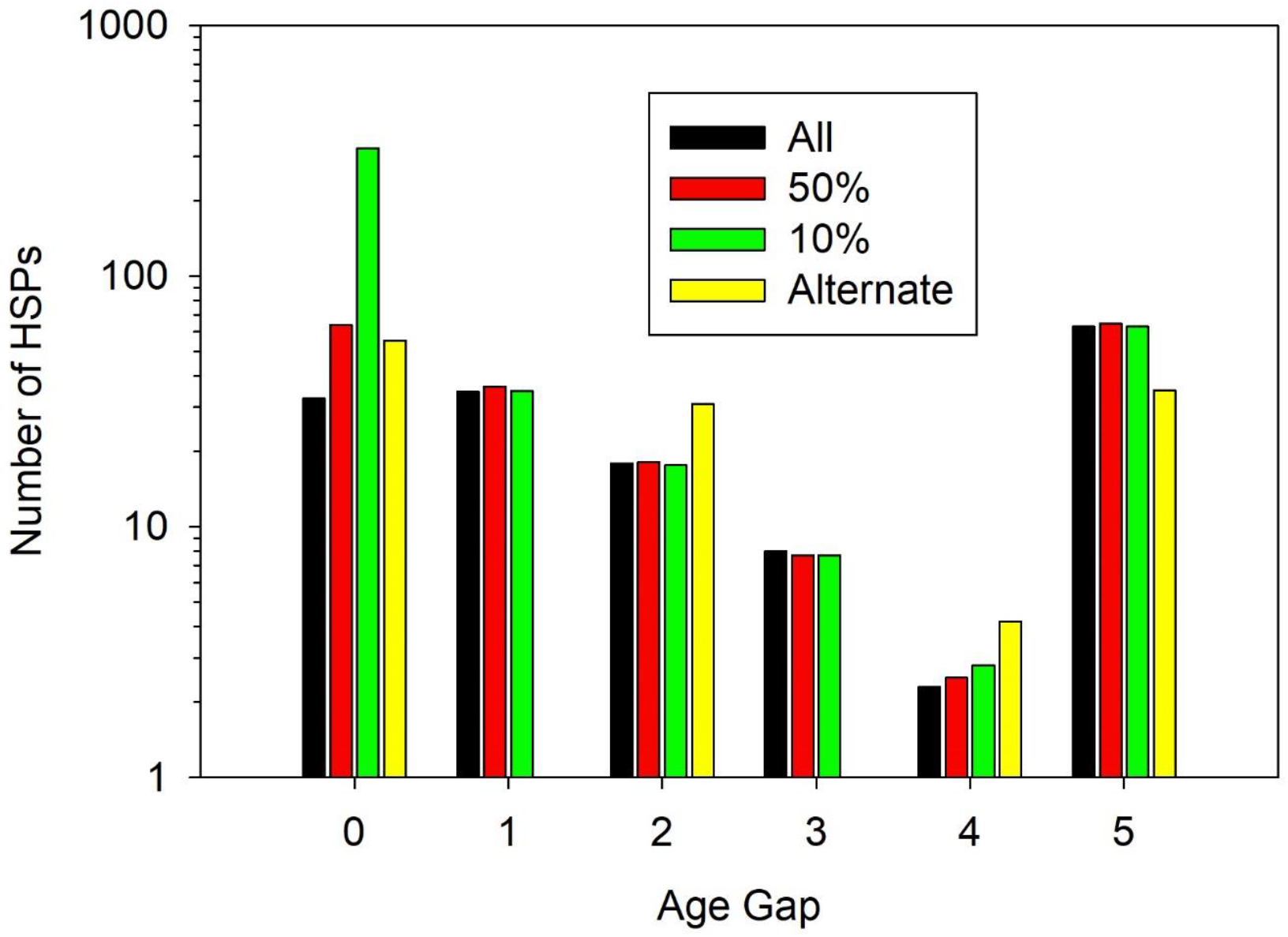
Mean numbers of siblings recorded in the simulations as a function of the age gap between offspring, for scenarios that involved intermittent breeding. The last column shows the total number of cross-cohort siblings found (age gaps 1-4). The scenario labeled ‘Breeders: All’ (black bars) is scenario A from Table 3. In the ‘50%’ and ‘10%’ scenarios (red and green bars), only a random half or 10% of all adults were allowed to reproduce each year; in the ‘Alternate’ scenario (yellow bars) only individuals aged 3, 5, 7, or 9 were allowed to reproduce, so every surviving parent skipped one year after reproduction. Note the log scale on the *Y* axis.

#### 5.2.2 Precision

CKMR methods are categorical (each pairwise comparison is either a POP/HSP or not), so only integer numbers of close-kin matches are possible (Figure 4). [In theory, probabilistic kin assignments could be used, as they commonly are for calling genotypes in genomics-scale datasets (Korneliussen et al. 2014), but this would cause hideously daunting consequences for CKMR likelihood calculation, for little gain in demographic information—M. Bravington, pers com. January 2021.] Precision of CKMR estimates is primarily determined by two key parameters: mean and variance in number of close-kin matches (*R*).

**Figure 4.**
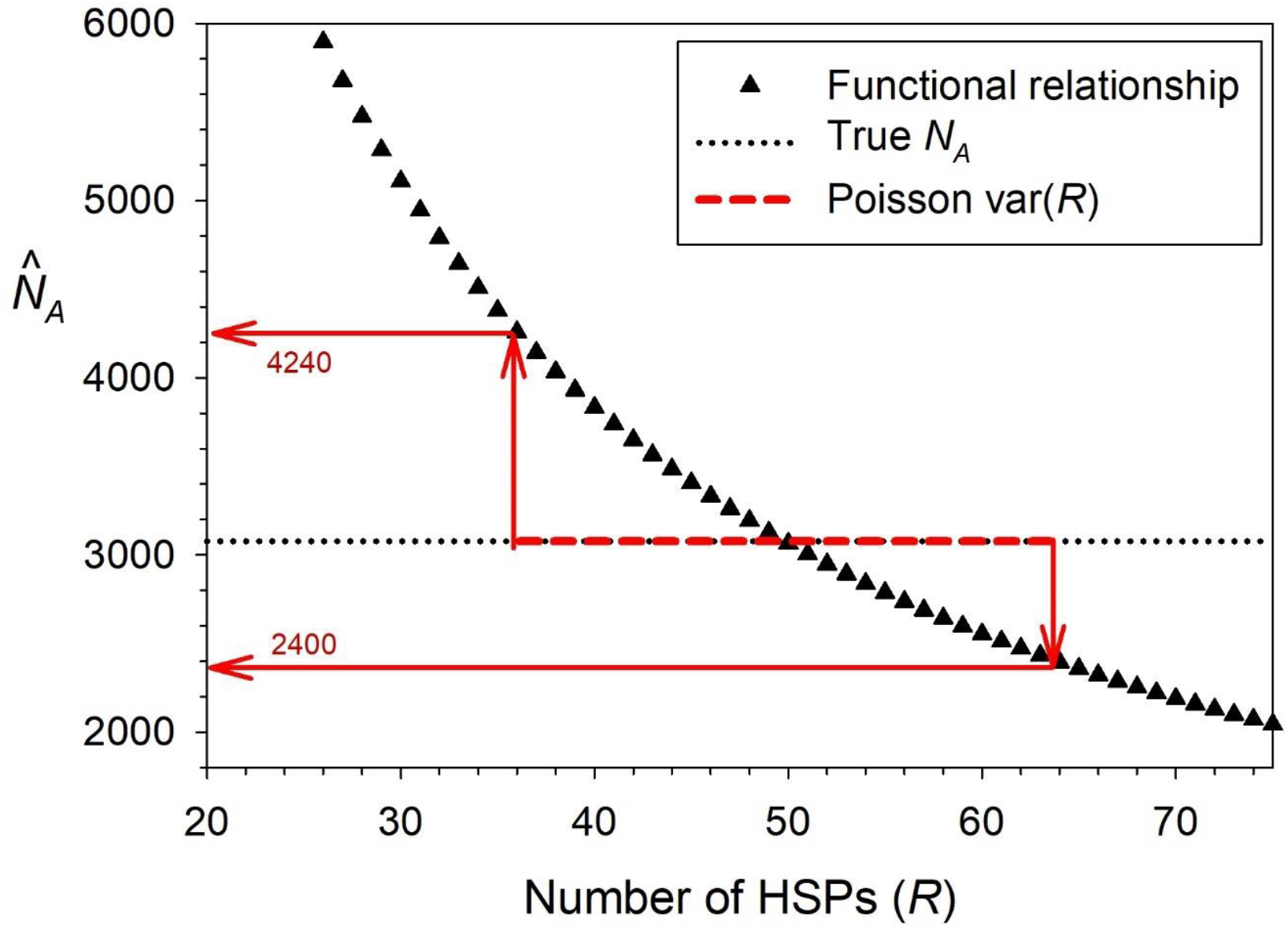
Precision in estimating adult census size (*N*_*A*_) using CKMR analyses based on the number of half sibling pairs (HSPs). Data are for Scenario A from Table 3, with true *N*_*A*_ = 3078 (black dotted line) and 89 offspring sampled from 5 consecutive cohorts. Black triangles are possible estimates of *N*_*A*_ using *R* = integer total numbers of HSPs (maternal and paternal combined) in Equation 7, with the mean conversion factor 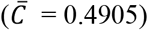 playing the role of the function *f*_*2*_. This level of sampling produces an average of 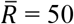 sibling matches which, if var(*R*) were Poisson, would produce a standard error of 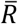 of about 7 and a 95% confidence interval (CI) around 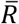 of [36-64] (dashed red line); this in turn would (following red arrows) lead to a 95% CI of 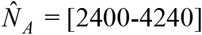. In the simulations, actual var(*R*) among replicates was slightly higher than the Poisson expectation (1.13 times the mean; Figure S2B), which would lead to a slightly wider CI, but the empirical 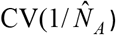 was still close to 0.15 (Figure 5C).

A crude approximation to a minimum CV for 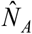 from MR or CKMR is 1/√*R* (Bravington et al. 2016b), in which case *R*=50 would translate into a CV of about 15%. This rule-of-thumb assumes a Poisson variance in *R*, implying that var(*R*) ≈ 50. In our simulations we monitored var(*R*) and compared it to the Poisson expectation. Also, because of the inverse relationship between 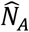 and 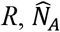 is skewed high and is infinitely large when no recoveries are found. This same issue applies to Wang’s sibship method (see Equation 4), and for this reason it is common for evaluations of precision of effective size estimators to focus on the distribution of 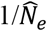 rather than 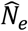, and to use the harmonic mean rather than arithmetic mean as a measure of central tendency (e.g., Wang 2009; Waples and Do 2010). Accordingly, here we calculate empirical CVs of 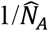 and 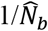 based on variances across replicates, as well as medians or harmonic means.

##### 5.2.2.1 Parent-offspring pairs

We found that var(*R*) among POP replicates was generally close to or slightly smaller than the Poisson variance (Figure S2A), which implies that in the simplest forms of POP-based CKMR the above rule-of-thumb is fairly accurate. An exception occurred for Scenario B, which had substantially overdispersed variance in reproductive success (ϕ=10); in this case, the large within-age effect on 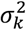 caused var(*R*) to be over twice as large as the mean (Figure S2A), which also considerably widened the distribution of 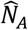 (Figure 2).

##### 5.2.2.1 Siblings

For the same scenarios from Table 3, the ratio var(*R*)/mean(*R*) was generally slightly higher for sibling analyses than for POPs (Figure S2A). For scenarios B (ϕ=10) and C (fecundity increasing with age), var(*R*)/mean(*R*) for siblings was > 1 (2.22 and 1.23, respectively), which presumably reflected the fact that both of these scenarios included unequal parental weights.

For a given life-history scenario, the ratio var(*R*)/mean(*R*) (and hence precision) is sensitive to adult abundance. As illustrated in Figure 5 (Panels A and B), although larger samples of individuals are needed from larger populations to produce a fixed number of close-kin matches, required sample sizes increase more slowly than adult abundance, so in very large populations only a small fraction of the population has to be sampled each year. This occurs because increasing the sample size by a factor *X* increases the number of possible comparisons to search for siblings by the factor *X*^2^. In theory, the effects of overdispersed variance and other factors that cause non-independence of the data should attenuate as sampling becomes sparse compared to overall abundance (Bravington et al. 2016b).

**Figure 5.**
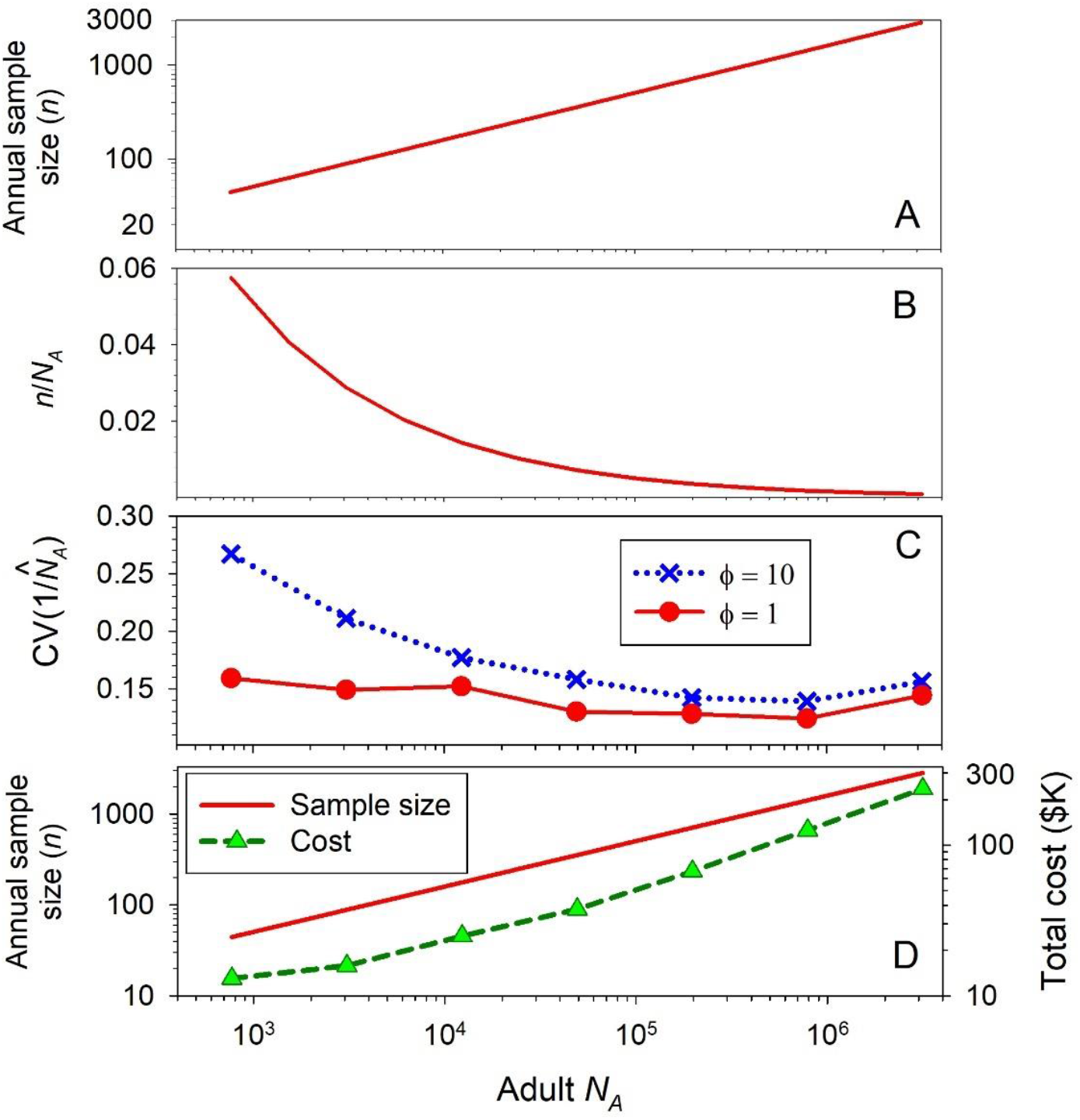
Relationships between annual sample size (*n*), adult abundance (*N*_*A*_), precision, and cost. In panels A and B, the red line represents sample sizes that are expected to produce a total of 50 cross-cohort half siblings, assuming samples are collected for 5 consecutive years. Panel C shows results of simulations based on two reproductive-skew variations described in Table 3: Scenario A (ϕ=1, solid red line and symbols) and Scenario B (ϕ=10, dotted blue line and symbols). Cohort sizes were varied to produce adult abundances that ranged more than three orders of magnitude, and corresponding sample sizes were adjusted as shown in Panel A to produce a constant expected number of 50 total HSPs. In panel D, the green dashed line shows approximate total cost (across all 5 years) for SNP discovery and genotyping for sample sizes expected to produce 50 HSPs. Vertical axes: A) Annual sample size, *n*; B) *n* as a fraction of *N*_*A*_; C) CV of 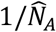: D) left, *n*; right, total cost in thousands of USD. Note the log scale on the *X* axis, and on the *Y* axes in Panels A and D.

Our simulations of populations with large *N*_*A*_ produced results consistent with this theory. For the smallest populations we modeled (*N*_*A*_ = 770), var(*R*)/mean(*R*) was somewhat elevated (1.26) from the Poisson expectation of 1.0 even for ϕ=1, but the variance rapidly dropped to at or below the Poisson variance as population size increased (Figure S2B). With moderately high within-age variance in reproductive success (ϕ=10), departures from the Poisson expectation are greater and the rate of attenuation lower, but even so var(*R*)/mean(*R*) dropped to ∼1.0 for adult abundances > 10^5^ (Figure S2B). All of these scenarios were designed to produce an average of 50 total half-sibling matches, and those that achieved a var(*R*)/mean(*R*) ratio close to 1.0 produced CVs of 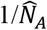 that were close to the expected 0.15 (Figure 5C).

For two scenarios (A and B from Table 3), we compared CKMR precision for estimating *N*_*A*_ with precision to estimate *N*_*b*_ using Wang’s sibship method, as a function of the number of consecutive years of samples that were used (2-5). In Scenario A, ϕ = 1 and fecundity was constant with age, so *N*_*A*_ = *N*_*b*_ = 3078; precision was higher for 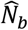 with only 2 years of data, but for longer studies precision of 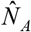 was higher (Figure 6). This latter result occurred because the total number of pairwise comparisons increases faster for cross-cohort comparisons used in CKMR than it does for within-cohort comparisons used to estimate *N*_*b*_. Results for Scenario B were quite different: with ϕ = 10, true *N*_*b*_ (388) is greatly reduced compared to adult *N* (3078), and this produces a stronger drift signal that is relatively easier to estimate precisely.

**Figure 6.**
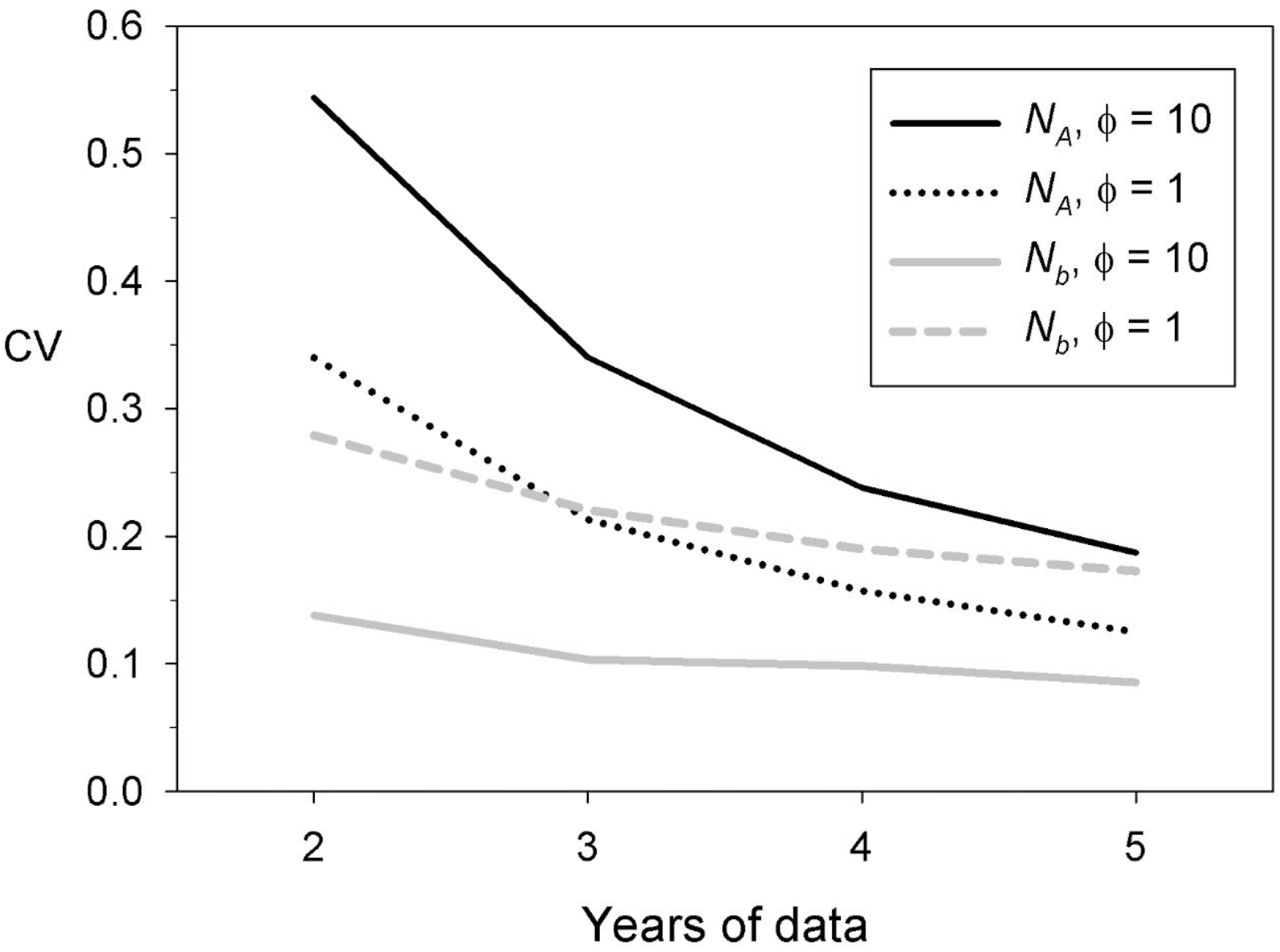
Precision for using close-kin sibling data to estimate adult *N*_*A*_ (black lines) or *N*_*b*_ (gray lines) as a function of the number of consecutive years during which samples are collected. CV (*Y* axis) is the coefficient of variation of 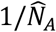 or 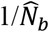, using data for all years. Fecundity was constant with age and true *N*_*A*_ was 3078. Results apply to Scenario A (ϕ = 1, solid lines; true *N*_*b*_= 3078) or Scenario B (ϕ = 10, dashed and dotted lines; true *N*_*b*_ = 388).

Conversely, the high variance in reproductive success substantially increased variability in the number of cross-cohort sibling matches. As a consequence, 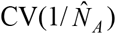 was 4 times as large as 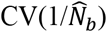 for 2 years of samples and still more than twice as large with 5 years of samples (Figure 6). These results illustrate an important difference between CKMR and estimators of effective size: overdispersed variance in reproductive success reduces precision of 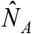 but increases precision of 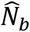 and 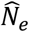.

CKMR and Wang’s sibship method share a limitation in that once enough genetic markers are used to reliably identify all close kin, additional markers cannot further increase precision, unless doing so allows more distant kin relationships to be integrated into the analysis. The LD method does not suffer from this limitation; in fact, the number of pairs of loci that can be used to compute mean *r*^2^ increases with the square of the number of loci, so in theory precision to estimate *N*_*e*_ and *N*_*b*_ could become arbitrarily large with genomics-scale datasets. In practice, limited recombination within chromosomes and lack of independence of overlapping pairs of the same loci limit the information content in large genetic datasets (Thompson 2013; Waples et al. 2021). Despite these limitations, if more than roughly 500-2000 SNPs are used, precision of the LD method for estimating *N*_*b*_ will generally exceed the maximum possible precision of Wang’s sibship method, which occurs when all sibling relationships are correctly identified (Waples 2021).

#### 5.3 Costs

Data shown in Figure 5 provide an opportunity to illustrate the relationship between experimental design and genotyping costs (see also Table S5 for more details). The following parameters apply to prices discussed below:

- Costs are in $USD and apply to SNP discovery and genotyping, but not DNA extraction, using DArTseq/DArTcap technology (Jaccoud et al. 2001) as implemented in Feutry et al. (2020);
- Costs include discovering and genotyping 1,500+ high quality SNPs (low error rate, high call rate, and MAF>0.1);
- Costs assume 94 samples per plate, with 1 positive and 1 negative control;
- Costs include a fixed component for SNP discovery (1 plate of DArTseq @ $3800) + a fixed component for DArTcap SNP panel synthesis + $11.4/individual);
- Costs assume a species with at least moderate levels of genetic diversity; species with low diversity might require more sequencing at extra cost for SNP discovery.

Estimates are based on several CKMR projects conducted by CSIRO Hobart, using methods described in Feutry et al. (2020), which in our experience is sufficient for accurate inference of POPs and siblings in species with *N*_*A*_ up to ∼2×10^6^. These costs represent only one possible commercial option, but a proven one that is globally available. Other methodologies exist, some laboratories might be able to save money by performing analyses in-house, and costs are likely to keep dropping in the future. Nevertheless, estimates shown in Figure 5D can serve as a useful reference point (and upper limit) for planning and experimental design.

Data shown in Figure 5D are total costs for an experimental design that involves 5 consecutive years of samples at a level designed to produce about 50 total HSPs and 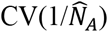 of about 0.15. Costs rise more slowly than population size for two reasons: a) the fraction of the population that has to be sampled to produce a fixed level of precision declines as *N*_*A*_ increases (Figure 5B); and b) in large studies, fixed costs of SNP discovery become relatively less important. Figures above do not include sampling, costs for which can be substantial but will vary widely across applications. On the other hand, resulting genotypes should be suitable for performing all of the analyses discussed in this paper, including identification of POPs and siblings and calculation of mean *r*^2^ for the LD analyses.

Also not included are costs for mtDNA analysis to estimate the fractions of HSPs that are of maternal and paternal origin, which can be modelled using haplotype frequencies (Bravington et al. 2016b; Thomson et al. 2020). Costs are modest (∼5K$ per 50 HSPs) because only the close-kin matches have to be tested but might be relatively important for small studies. For POPs, parental sex is generally determined morphologically or using sex-specific markers.

## 6 DISCUSSION

CKMR estimates of abundance and genetic methods to estimate effective population size share a dependence on core features of population demography: means and variances of age-specific vital rates and covariances in realized reproductive success over time. Effective size can be predicted based on a vector of parental weights, ***W***, that is mathematically equivalent to the expected-relative-reproductive-success (*ERRO*) concept that underpins CKMR. A CKMR study will generate some or all of the demographic data needed to calculate effective size directly, and genotypes for the sampled individuals provide an opportunity to apply indirect genetic methods for estimating *N*_*e*_ or *N*_*b*_. Nevertheless, demographic parameters and other covariates influence CKMR and effective size estimators in different ways.

### 6.1 CKMR

As illustrated in Equations 8 (for POPs) and 10 (for siblings), the probability of a close-kin match is inversely proportional to total reproductive output (*TRO*) of the population. *TRO* is closely related to spawning stock biomass, which typically is an important metric in fisheries stock assessments. In many other applications, however, adult abundance is the parameter of primary interest, in which case it is necessary to disentangle the relationship between *TRO* and the *N*_*A*_ adults responsible for that reproductive output. For some simple scenarios, this can be done using Equation 7 and a scalar conversion factor *C*, calculated as described in detail in Supporting Information S2.2. This approach was adopted for analyzing the simulated data, which illustrated predictable effects of some common factors on performance of CKMR and effective size estimators. Key results that were demonstrated include the following:

- Large population size does not by itself cause bias (Figure S1; Table 3), but large populations require larger sample sizes (*n*) to detect the same number of close kin (*R*), where E(*R*) = *Y*\**P*_*match*_. However, the number of pairwise comparisons (*Y*) increases with *n*^2^ while the probability of a match declines only according to 1/*N*_*A*_. Therefore, the fraction of the population that must be sampled to achieve a fixed level of precision is smaller in large populations, which helps reduce costs for large projects (Figure 5).
- Precision of CKMR estimates primarily depends on two factors: mean and variance in number of close-kin matches. The relationship between 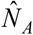 and *R* is inverse and non-linear, so as *R* increases the curve becomes flatter (Figure 4). With respect to precision, therefore, the experimental-design goal should be to make E(*R*) large enough that random sampling error and limited errors in pedigree reconstruction have relatively little influence on 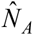. Our simulations showed that the assumption of a Poisson variance to *R* is generally reasonable, at least for large populations. But the empirical var(*R*) can be substantially higher than the Poisson expectation in populations of small to moderate size, and this effect is exacerbated when adults have very unequal probabilities of producing offspring. To compensate for this increased variance and maintain a target level of precision, it might be necessary to boost the sampling effort to increase E(*R*).
- Sibling-based CKMR analyses should exclude same-cohort comparisons, whereas Wang’s method is restricted to single cohorts. Like Jack Sprat and his wife^1^, the two methods neatly carve up the data into non-overlapping layers. Relative precision to estimate *N*_*A*_ and *N*_*b*_ from the same experimental design depends on three factors: the number of years of samples, annual mortality, and variance in reproductive success (Figure 6). In longer studies, the number of cross-cohort comparisons increases faster than those within cohorts, which increases relative precision of 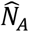; however, this advantage is partially offset by adult mortality, which reduces the effective number of cross-cohort comparisons. Overdispersed variance in reproductive success (ϕ>1) increases var(*R*) and 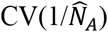 but has the opposite effect on 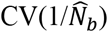 (Figure 6), because large ϕ reduces *N*_*b*_ and increases the drift signal.
- Although ϕ>1 increases var(*R*) and reduces precision for CKMR (except perhaps in very large populations), it does not bias abundance estimates based on either POPs or cross-cohort siblings (Figures 2 and S1).
- *P*_*POP*_ is a simple inverse function of 1/*N*_*A*_, leading to unbiased estimates of adult abundance, provided that there are no unmodelled covariates that affect both an adult’s sampling probability AND its *ERRO* among the sampled offspring. If *ERRO* and sampling probability of adults both increase over time (as in Scenario D, Table 3), this potential source of bias can be accounted for as illustrated in Table S1.
- POPs-based abundance estimates depend heavily on which individuals are included in the sample of potential parents. If non-breeding adults are not sampled, the estimate will apply to the number of breeders in a given year (Figure 1A); conversely, if immature individuals are sampled as potential adults, the naive estimate will be skewed away from *N*_*A*_ toward overall *N* (Figure 2).
- Cross-cohort sibling comparisons provide information about adult abundance in the year the younger sibling was born (Equation 11). POP comparisons provide information about adult abundance in the year(s) offspring were born. Sampling from multiple cohorts thus can provide information about population trend.
- It is always necessary to consider adult mortality for sibling-based CKMR. This source of random noise increases var(*R*) and reduces precision, even when average mortality rates can be accurately estimated (Figure S2).
- Random and uncorrelated individual variation in *ERRO* does not bias sibling-based estimates of abundance (Figure 3). However, factors that cause positive or negative correlations in *ERRO* over time do affect *P*_*MHSP*_ (Equation 11) and must be accounted for to avoid bias. Effects of some of these factors, such as intermittent breeding and persistent individual differences, are relatively straightforward (Figure 1). Less intuitive is the fact that any systematic change in *ERRO* with age also establishes predictable correlations in individual reproductive success over time, and these effects can be accounted for as illustrated in Table S2.

Despite its heuristic value, the simplified approach outlined in Equation 7 and illustrated by the simulated examples has some important limitations. For simplicity and tractability, we assumed that:

- All close kin were correctly identified;
- The single, closed population was constant in size;
- Sex ratio was equal and vital rates were constant over time and known without error;
- All sampled individuals could be aged accurately;
- Sampling was random unless adults were sampled in proportion to relative fecundity, which also was known precisely.

In the sibling simulations, these assumptions allowed us to combine all sibling matches across five years of samples to estimate a single, constant parameter (*N*_*A*_ for CKMR, *N*_*b*_ for Wang’s sibship method).

None of the CKMR applications published to date meet all these assumptions. More generally, CKMR experimental designs involve sampling and testing for POPs and/or siblings from multiple time periods, across which abundance and vital rates can change, perhaps sharply. Incorrectly assuming parameter stationarity can easily lead to bias (Skaug 2001). When values of key covariates are uncertain, the estimation process can integrate over plausible ranges, albeit at a cost with respect to precision. For all these reasons, the most robust implementations of CKMR incorporate information from parent-offspring pairs and siblings into a single, log-pseudo-likelihood framework, within which all demographic parameters can be jointly estimated in a way that allows coherent statements about uncertainty.

Another major benefit of using an integrated modeling framework is that many, if not all, key covariates that affect *P*_*match*_ can also be estimated from the data. For example, with POPs plus ages one can estimate age-specific fecundity. Adult survival can be inferred from the distribution of age gaps between siblings (Hillary et al. 2018; see also Table S3). Intermittent breeding can be evaluated using both POPs and HSPs by examining lag times between repeated offspring assignments to the same parent (Bravington et al. 2016a; Thomson et al. 2020). To the extent that independent information about covariates is available from other sources, this can be included within the overall estimation process.

### 6.2 Effective size

Many of the factors discussed above that complicate CKMR abundance estimates are part of the genetic drift signal and hence create no problems for genetic methods that estimate annual *N*_*b*_. This applies to age-specific changes in vital rates, skewed sex ratios, skewed offspring distributions, and skip breeding. As with CKMR, however, estimates of effective size can be very sensitive to non-random sampling that leads to positive correlations of sampling probabilities among relatives (Goldberg and Waits 2010; Whiteley et al. 2012). Although detailed life-history and age-structure information for the population as a whole is necessary for demographic estimates of *N*_*e*_ or *N*_*b*_, ageing of individuals is primarily important for sorting offspring into single cohorts. Single-sample genetic methods estimate the effective number of breeders that produced the sampled offspring (Waples 2005; Wang 2009). Researchers should keep in mind that, depending on the experimental design and the biology of the target species, this might not represent the entire adult population (e.g., under skip breeding in Figure 1).

### 6.3 Practical applications

Meaningful CKMR estimates require (1) genetic data robust enough to reliably identify close-kin pairs; and (2) sufficient sampling effort to produce the desired level of precision. We have assumed that all close-kin matches are correctly identified, which implies that criterion (1) has been met (see Bravington et al. 2016b for a summary of basic principles of kin identification). In practice, some level of uncertainty is associated with both genotyping and kin identification, so pedigree-reconstruction models should account for potential errors and evaluate their consequences.

To date, CKMR has only been applied to a handful of species, but these cover a wide range of population sizes and life histories, from relatively small populations of salmonids (Rawding et al. 2014; Ruzzante et al. 2019) and fecundity-limited sharks (Hillary et al. 2018) to a large, mobile, highly-fecund tuna with 2 million estimated adults (Bravington et al. 2016a). Most of these applications fall into one of two categories: at-risk species with high conservation value, or high-value fisheries. All published applications to date are for aquatic species, but CKMR methods can also potentially be applied to terrestrial species, as is commonly done with genetic methods to estimate effective size.

Even when various factors constrain the types of analyses that are feasible, it can be possible to extract valuable information using CKMR. Despite the inability to reliably sample adults, juvenile samples of the cryptic and critically endangered speartooth shark, *Glyphis glyphis*, have provided important insights into connectivity (Feutry et al. 2017). Inability to precisely age the school shark, *Galeorhinus galeus*, added considerable uncertainty to CKMR estimates of abundance, which nevertheless were more precise than those from fishery-dependent methods (Thomson et al. 2020; Simpfendorfer et al. 2021). Among the most important factors that affect the feasibility of CKMR methods are the following (see also Tables 4 and 5):

**Table 4.** Summary of major factors that, unless explicitly accounted for, can influence performance of CKMR methods to estimate abundance. ‘RS’ = reproductive success.

| Factor | POPs | HSPs |
| --- | --- | --- |
| Ageing individuals | Necessary if required to separate parents/offspring, account for changes in fecundity with age, or calculate ERRO in the year of offspring’s birth | Required to sort individuals into cohorts and calculate age gaps |
| Sampling of cohorts | Samples from single cohorts can be used for semelparous species. Sampling multiple cohorts is essential for most iteroparous species | Sampling multiple cohorts is essential because only cross-cohort comparisons provide information about census size |
| Age-specific fecundity | Affects precision; needs to be estimated if selectivity varies with age | Affects precision and bias; can be estimated independently or from POPs + ages |
| Adult mortality | Not required unless sampling of adults is non-lethal | Required to adjust probability a parent produces sibs in different years, but can be estimated from HSP data |
| Overdispersed variation in RS ($\phi > 1$) | Does not affect $P_{POP}$ but reduces precision | Reduces precision. For bias, see below |
| HSPs from the same cohort | | Siblings from same cohort estimate $N_b$, not adult $N$ |
| HSPs from different cohorts |  |  |
| Residual RS uncorrelated across years | | No bias in $\hat{N}_A$ |
| Persistent individual differences in RS | | Increases $P_{HSP}$ and will bias $\hat{N}_A$ unless accounted for |
| Skip breeding | Downward bias in $\hat{N}_A$ if skip breeders not sampled. With adequate samples, pattern of skip breeding can be estimated from the data | Complex effects on $P_{HSP}$. With adequate samples, pattern of skip breeding can be estimated from the data |
| Non-random sampling concerns | POPs collected together more often than by chance; Probability of sampling an adult correlated with its RS | Siblings born in different years collected together more often than would occur by chance |
| Precision | In simple scenarios with $\phi \approx 1$, $CV(1/\hat{N}_A) = 0.15$ requires ~50 POPs, which implies sample sizes on the order of $\sqrt{N_A}$ individuals | Somewhat more sensitive than POPs to factors that inflate variance, but $var(R_{HSP})$ generally converges on $\bar{R}_{HSP}$, for large $N$ |
| Benefits of genomics-scale datasets | Scant: performance asymptotes at low numbers of genetic markers after POPs reliably identified | Once all siblings are correctly identified (which generally requires a few thousand SNPs), no benefits from adding more markers |

**Table 5.** Summary of major factors that, unless explicitly accounted for, can influence performance of some genetic methods to estimate effective size. ‘RS’ = reproductive success. 43

| Factor | Sibship method | LD method |
| --- | --- | --- |
| Ageing individuals | Required to sort individuals into cohorts to estimate $N_b$ or to estimate $N_e$ from mixed-age samples | Required to sort individuals into cohorts to estimate $N_b$; not required to estimate $N_e$ from mixed-age samples |
| Sampling of cohorts | A single-cohort sample provides a point estimate of $N_b$. Multiple such samples provide a temporal series of estimates | A single-cohort sample provides a point estimate of $N_b$. Multiple such samples provide a temporal series of estimates. Mixed-age samples can estimate $N_e$ |
| Changes in fecundity or mortality with age; skewed sex ratio; overdispersed variance in RS ($\phi > 1$) | No bias; these are part of the signal of reduced $N_e$ and $N_b$ | No bias; these are part of the signal of reduced $N_e$ and $N_b$ |
| Skip breeding and persistent individual differences in RS | No bias to single-cohort samples estimating $N_b$ | No bias to estimates of $N_b$ or $N_e$, but this affects the ratio $N_b/N_e$ and influences the bias adjustments to account for age structure |
| Non-random sampling issues | Estimate is downwardly biased if siblings are collected together more often than would occur by chance | Estimate is downwardly biased if relatives are collected together more often than would occur by chance |
| Precision | For large populations, sample sizes on the order of $\sqrt{N_e}$ individuals | Precision jointly determined by samples of individuals and loci |
| Benefits of genomics-scale datasets | Once all siblings are correctly identified (which generally requires a few thousand SNPs), no benefits from adding more markers | With more than about 500-2000 SNPs, precision of LD method will meet or exceed maximum precision possible with sibship method |
| Changes in $N$ and/or $N_e$ | No effect on estimates based on single cohorts | Some bias toward recent $N_e$ for 3-5 generations |

#### 6.3.1 Biology

- Long lifespan and/or precise ageing facilitates sampling of multiple cohorts
- Low adult mortality increases precision of HSPs
- For HSPs, it is necessary to be able to either a) assume that fecundity is constant with age, or b) use independent sources of information, or c) estimate *b*_*x*_ from CKMR data, generally using POPs + ages.
- In small populations, the lack of independence of close-kin comparisons can become an issue.

#### 6.3.2 Ageing

- Minimum requirements: must be able to sort into potential parents and offspring (POPs) or into consecutive cohorts (siblings; so age gap is known)
- Need actual ages to estimate age-specific fecundity with POPs
- Uncertainty in ageing can be modeled but can substantially reduce precision

#### 6.3.3 Sampling

- Must be able to sample individuals independent of their kinship, or model lack of independence
- For POPs, if adults cannot be sampled independent of their *ERRO*, must be able to estimate selectivity

### 6.4 Future considerations

We expect to see some or all of the following developments in the future:

- As whole-genome-sequencing of non-model species becomes more common, it will become feasible to extend the array of close-kin categories available to CKMR beyond the one-generation pedigree to include more distant relationships, including cousins and niblings (nieces and nephews).
- Conversely, as long-term studies continue, complications posed by other relationships that have the same kinship probabilities as half siblings (e.g., grandparent – grandchild) will increase and require effective treatments. Current applications often can ignore these complications because of the species’ biology (esp delayed age at maturity) and experimental design.
- Single-sample genetic estimators can be used to provide a continuous time-series of annual estimates of *N*_*b*_, which are generally done independently each year (Ruzzante et al. 2016; Whitely et al. 2015; Waples et al. 2018a,b). One could develop an analytical framework analogous to that used by CKMR that jointly considers multiple years of data to estimate temporal trajectories of *N*_*b*_ and other population parameters, but we are not aware of any attempts to do that. Furthermore, because CKMR and the two single-sample estimators of effective size use the same data, it should be possible to incorporate all of the estimation of annual abundance, *N*_*b*_, and related parameters into a single pseudo-likelihood framework that would help shed light on the important ratio of effective size to census size. An initial attempt to combine CKMR and effective size information, based only on POP data for microsatellites used in the CKMR study of southern bluefin tuna, showed that the range of *N*_*e*_/*N* ratios that were most consistent with the combined data was about 0.1 to 0.5 (Figure 7). This ruled out any substantial effect of sweepstakes reproductive success for southern bluefin tuna, even though the species’ longevity and high fecundity are similar to several marine species for which “tiny” estimates of *N*_*e*_/*N* have been reported (Hedgecock and Pudovkin 2011).
- Tightly linked markers on the same chromosome provide insights into *N*_*e*_ in the more distant past. If detailed information about genomic structure is available, it can be leveraged to provide a temporal series of 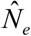 estimates back in time. Applications of this approach include humans (Tenesa et al. 2007), red drum (*Sciaenops ocellatus*; Hollenbeck et al. 2016), and Atlantic salmon (*Salmo salar*; Lehnert et al. 2019), and a recently-developed modification (Santiago et al. 2020) appears to considerably improve performance. Apart from using more distant kin to extend pedigree inference back one or a few more generations, there does not appear to be any comparable potential for CKMR to provide insights into historical abundance.

**Figure 7.**
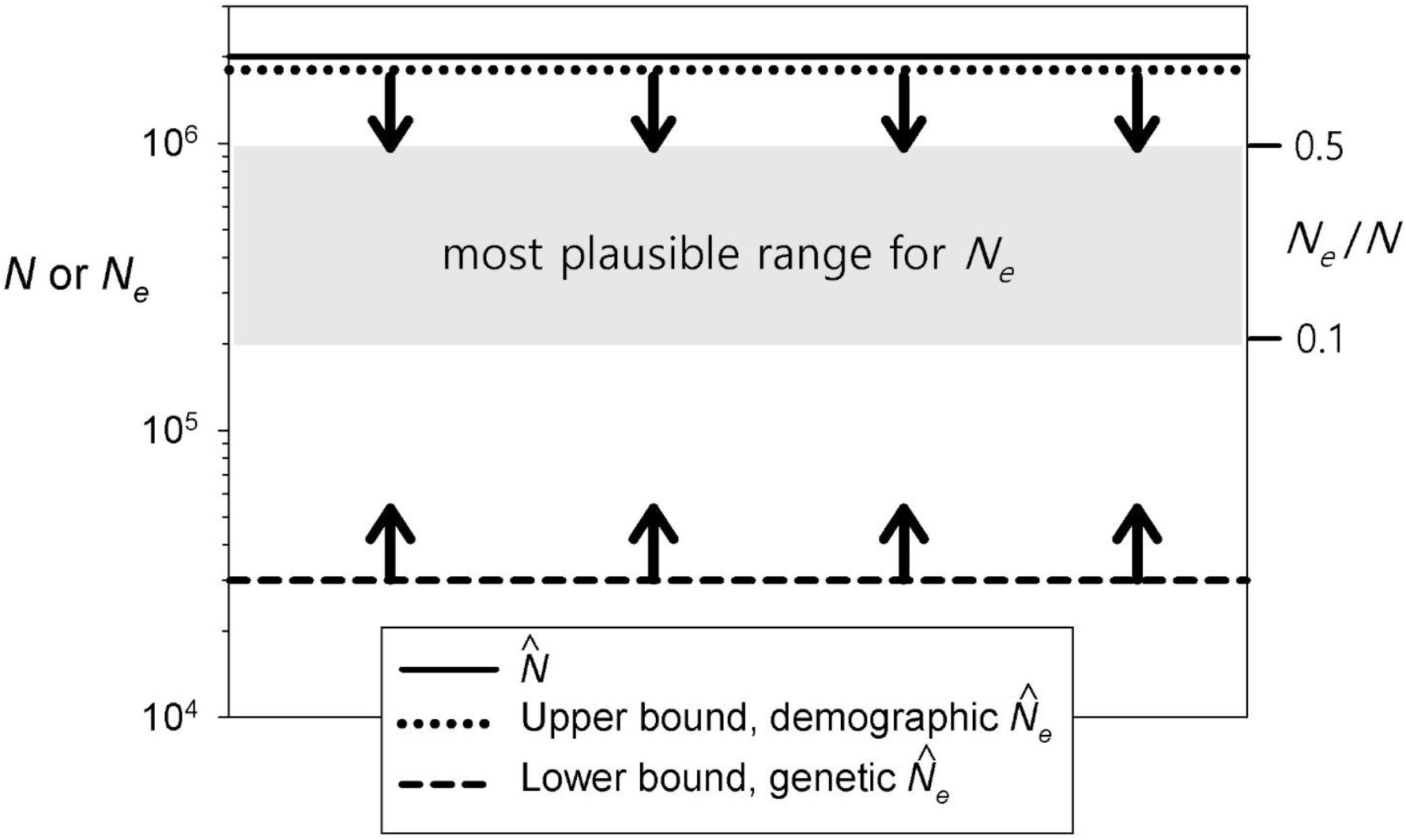
Using complementary properties of genetic and demographic estimators to bracket the plausible range for the ratio *N*_*e*_/*N*_*A*_ in southern bluefin tuna. The solid black line is the CKMR estimate of adult abundance (2×10^6^) from Bravington et al. (2014); Waples et al. (2018a) used the same genetic and demographic data, respectively, to delimit a lower bound (dashed line) and an upper bound (dotted line) to *N*_*e*_. Genetic methods can provide robust lower bounds for *N*_*e*_ but, because the drift signal is so small, have difficulty distinguishing large from very large or infinite effective sizes. Conversely, life history constrains how large *N*_*e*_ can be but, without very detailed demographic information, cannot rule out very small *N*_*e*_/*N*_*A*_ ratios. Collectively, and in conjunction with a robust estimate of *N*_*A*_ from CKMR, the two effective-size methods constrain the plausible range of *N*_*e*_/*N*_*A*_ to about 0.1-0.5.

Finally, an aspirational goal would be to integrate estimation of abundance with CKMR and estimation of generational effective population size, *N*_*e*_. This should uncover some interesting parallels: unlike annual *N*_*b*_ but similar to sib-based CKMR, *N*_*e*_ per generation is sensitive to correlations in individual reproductive success over time. Hill’s (1972) model for *N*_*e*_ when generations overlap is robust to random demographic variation if *N* is not too small (Engen et al. 2005a; Waples et al. 2011). However, allowing for environmental stochasticity requires a different formulation (Engen et al. 2005), and even this becomes problematical under strong density dependence, when vital rates vary over time and generation length is not well defined (Myhre et al. 2016). Successfully integrating generational *N*_*e*_ and CKMR into a single estimation framework thus remains a challenging prospect.

## Acknowledgments

Mark Bravington contributed a great deal to our understanding of the material presented here. We thank Eric Anderson, Peter Grewe, Daniel Ruzzante, and Hans Skaug for useful discussions. Marco Andrello, Mark Bravington, Ryan Kovach, Jann Martinsohn, Todd Seamons, Antonella Zanzi, and two anonymous reviewers provided insightful comments on earlier versions of the manuscript. Eric Anderson kindly modified his CKMRpop simulation package to run under Windows and performed some simulations for population sizes (adult *N* > 10_5_) that were too large to run locally. RW’s work on this project was supported by an International Science Fellowship from the National Marine Fisheries Service and by a Froelich Fellowship from CSIRO. The authors declare no conflict of interest. This manuscript did not generate new empirical data.

## Footnotes

1 Jack Sprat could eat no fat. His wife could eat no lean. But, together both, they licked the platter clean (Opie and Opie 1997, p. 238)

## References

Akita, T., 2020a. Nearly unbiased estimator of contemporary Ne/N based on kinship relationships. Ecology and evolution, 10:10343–10352.

Akita, T., 2020b. Nearly unbiased estimator of contemporary effective mother size using within-cohort maternal sibling pairs incorporating parental and nonparental reproductive variations. Heredity 124:299–312.

Anderson EC. 2021. CKMRpop: Forward-in-time Simulation and Tabulation of Pairwise Relationships in Age-structured Populations v. 0.1.0 (github.com/eriqande/CKMRpop).

Arnold, T.W., Clark, R.G., Koons, D.N. and Schaub, M., 2018. Integrated population models facilitate ecological understanding and improved management decisions. The Journal of Wildlife Management, 82(2), pp.266–274.

Blower, D.C., Riginos, C. and Ovenden, J.R., 2019. NeOGen: A tool to predict genetic effective population size (Ne) for species with generational overlap and to assist empirical Ne study design. Molecular ecology resources, 19:260–271.

Bradford, R, Thomson, R, Bravington, M, Foote, D, Gunasekera, RM, Bruce, B, Harasti, D, Otway, N, Feutry, P. 2018. A close-kin mark-recapture estimate of the population size and trend of east coast grey nurse shark. Report to the National Environmental Science Program, Marine Biodiversity Hub. CSIRO Oceans & Atmosphere, Hobart, Tasmania. available at: https://www.nespmarine.edu.au/document/close-kin-mark-recapture-estimate-population-size-and-trend-east-coast-grey-nurse-shark.

Bravington, M.V., P. G. Grewe, C. R. Davies. 2014. Fishery-independent estimate of spawning biomass of Southern Bluefin Tuna through identification of close-kin using genetic markers. FRDC Report 2007/034, Australia, 2014.

Bravington, M.V., Grewe, P.M. and Davies, C.R., 2016a. Absolute abundance of southern bluefin tuna estimated by close-kin mark-recapture. Nature communications, 7:13162.

Bravington, M.V., Skaug, H.J. and Anderson, E.C., 2016b. Close-kin mark-recapture. Statistical Science, 31:259–274.

Bravington, M., Feutry, P., Pillans, R.D., Hillary, R., Johnson, G., Saunders, T., Gunesekera, R., Bax, N.J. and Kyne, P.M. (2019). Close-Kin Mark-Recapture population size estimate of Glyphis garricki in the Northern Territory. Report to the National Environmental Science Program, Marine Biodiversity Hub. CSIRO Oceans and Atmosphere, Hobart.

Buckworth, R., Ovenden, J., Broderick, D., Macbeth, G., McPherson, G. and Phelan, M., 2012. GENETAG: Genetic mark-recapture for real-time harvest rate monitoring. Pilot studies in northern Australia Spanish mackerel fisheries. Darwin Australia: Northern Territory Government, Department of Resources; 2012. Available at: http://hdl.handle.net/102.100.100/100224?index=1

Caballero, A., 1994 Developments in the prediction of effective population size. Heredity 73: 657–679.

Caswell, H. 2001. Matrix Population Models, Ed. 2. Sinauer, Sunderland, MA.

Charlesworth, B., 2009. Effective population size and patterns of molecular evolution and variation. Nature Reviews Genetics, 10(3), p.195.

Conn, P.B., Bravington, M.V., Baylis, S. and Ver Hoef, J.M., 2020. Robustness of close-kin mark–recapture estimators to dispersal limitation and spatially varying sampling probabilities. Ecology and Evolution 10:5558–5569.

Crow, J.F., and C. Denniston, 1988 Inbreeding and variance effective population numbers. Evolution 42:482–495.

Davies C, Bravington M, and Thomson R. 2017. Advice on Close-Kin Mark-Recapture for estimating abundance of eastern Atlantic bluefin tuna: a scoping study. CSIRO Marine Laboratories, Hobart, Tasmania, Australia.

Doak, D.F., Gross, K. and Morris, W.F., 2005. Understanding and predicting the effects of sparse data on demographic analyses. Ecology, 86(5):1154–1163.

Engen, S., R. Lande, and B.-E. Sæther. 2005. Effective size of a fluctuating age-structured population. Genetics 170:941–954.

England, P.E., G. Luikart, and R.S. Waples. 2010. Early detection of population fragmentation using linkage disequilibrium estimation of effective population size. Conservation Genetics 11:2425–2430.

Felsenstein, J., 1971 Inbreeding and variance effective numbers in populations with overlapping generations. Genetics 68:581–597.

Fernandes, D.M., Sirak, K.A., Ringbauer, H. et al. 2020. A genetic history of the pre-contact Caribbean. Nature (https://doi.org/10.1038/s41586-020-03053-2; published 23 December 2020)

Feutry P, Berry O, Kyne PM, Pillans RD, Hillary RM, Grewe PM, Marthick JR, Johnson G, Gunasekera RM, Bax NJ (2017) Inferring contemporary and historical genetic connectivity from juveniles. Molecular Ecology 26:444–456.

Feutry, P, Devloo-Delva, F, Tran Lu Y A, Mona, S, Gunasekera, RM, Johnson, G, Pillans, RD, Jaccoud, D, Kilian, A, Morgan, DL, Saunders, T, Bax, NJ, and PM Kyne. 2020. One panel to rule them all: DArTcap genotyping for population structure, historical demography, and kinship analyses, and its application to a threatened shark. Molecular Ecology Resources 20(6): 1470–1485.

Fournier, D., Archibald, C.P., 1982. A general theory for analysing catch at age dat a. Can. J. Fish. Aquat. Sci. 39, 1195–1207.

Frankham, R., 1995. Effective population size/adult population size ratios in wildlife: a review. Genetics Research, 66(2):95–107.

Gilbert, K.J. and Whitlock, M.C., 2015. Evaluating methods for estimating local effective population size with and without migration. Evolution, 69(8): 2154–2166.

Goldberg CS, Waits LP (2010) Quantification and reduction of bias from sampling larvae to infer population and landscape genetic structure. Molecular Ecology Resources, 10, 304–313.

Hauser, L., and G. R. Carvalho. 2008. Paradigm shifts in marine fisheries genetics: Ugly hypotheses slain by beautiful facts. Fish Fish. 9:333–362.

Hedgecock, D, A. I. Pudovkin. 2011. Sweepstakes reproductive success in highly fecund marine fish and shellfish: A review and commentary. Bull. Mar. Sci. 87:971–1002.

Hedrick, P., 2005. Large variance in reproductive success and the Ne/N ratio. Evolution 59:1596–1599.

Hill, W.G., 1972 Effective size of population with overlapping generations. Theoretical Population Biology 3:278–289.

Hill, W. G., 1981 Estimation of effective population size from data on linkage disequilibrium. Genet. Res. 38:209–216.

Hillary, R.M., Bravington, M.V., Patterson, T.A., Grewe, P., Bradford, R., Feutry, P., Gunasekera, R., Peddemors, V., Werry, J., Francis, M.P. and Duffy, C.A.J., 2018. Genetic relatedness reveals total population size of white sharks in eastern Australia a nd New Zealand. Scientific reports, 8(1), p.2661.

Hollenbeck CM, Portnoy DS, Gold JR (2016) A method for detecting recent changes in contemporary effective population size from linkage disequilibrium at linked and unlinked loci. Heredity 117:207–16.

Holmes, E.E., 2001. Estimating risks in declining populations with poor data. Proceedings of the National Academy of Sciences, 98(9):5072–5077.

Jaccoud, D., Peng, K., Feinstein, D., & Kilian, A. (2001). Diversity arrays: A solid state technology for sequence information independent genotyping. Nucleic Acids Research, 29(4), e25.

Jolly, G. M. (1965). Explicit estimates from capture –recapture data with both death and immigration-stochastic model. Biometrika 52:225–247.

Kendall KC, Stetz JB, Roon DA, Waits LP, Boulanger JB, Paetkau D. 2008. Grizzly bear density in Glacier National Park, Montana. Journal of Wildlife Management 72:1693–1705.

Korneliussen, T.S., Albrechtsen, A. and Nielsen, R., 2014. ANGSD: analysis of next generation sequencing data. BMC bioinformatics, 15(1), p.356.

Krebs, CJ. 2009. Ecology: The Experimental Analysis of Distribution and Abundance, 6th Edition. Pearson International Edition. San Francisco, USA.

Krimbas, C.B. and Tsakas, S., 1971. The genetics of Dacus oleae. V. Changes of esterase polymorphism in a natural population following insecticide control —selection or drift?. Evolution, 25(3):454–460.

Lebreton, J.D., Burnham, K.P., Clobert, J. and Anderson, D.R., 1992. Modeling survival and testing biological hypotheses using marked animals: a unified approach with case studies. Ecological monographs, 62(1), pp.67–118.

Lee, A.M., S. Engen, and B.-E. Sæther, 2011 The influence of persistent individual differences and age at maturity on effective population size. Proceedings of the Royal Society of London, Series B: Biological Sciences, 278:3303–3312.

Lee, A.M., Myhre, A.M., Markussen, S.S., Engen, S., Solberg, E.J., Haanes, H., Røed, K., Herfindal, I., Heim, M. and Sæther, B.E., 2020. Decomposing demographic contributions to the effective population size with moose as a case study. Molecular ecology, 29(1):56–70.

Lehnert, SJ, T Kess, P Bentzen, MP Kent, S Lien, J Gilbey, M Clément, NW Jeffrey, RS Waples, and IR Bradbury. 2019. Genomic signatures and correlates of widespread population declines in salmon. Nature Communications 10:2996.

Lindberg, M.S., 2012. A review of designs for capture–mark–recapture studies in discrete time. Journal of Ornithology, 152(2):355–370.

Luikart, G., N. Ryman, D. A. Tallmon, M. K. Schwartz, and F. W. Allendorf. 2010. Estimation of census and effective population sizes: the increasing usefulness of DNA-based approaches. Conservation Genetics 11:355–373.

Myhre, A. M., S. Engen, B.-E. Sæther. 2016. Efective size of density-dependent populations in fluctuating environments. Evolution 70:2431–2446

Nei M, Tajima F (1981) Genetic drift and estimation of effective population size. Genetics 98:625–640.

Økland, J.M., Haaland, Ø.A. and Skaug, H.J., 2009. A method for defining management units based on genetically determined close relatives. Ices Journal of Marine Science, 67(3):551–558.

Oleksiak Marjorie F., and Om P. Rajora. 2020. Marine Population Genomics: Challenges and Opportunities. pp 3–35 in Population Genomics: Marine Organisms, Springer Nature, Cham Switzerland.

Opie, I, and Opie, P. 1997. The Oxford Dictionary of Nursery Rhymes (2nd ed.). Oxford University Press, Oxford, UK.

Palsbøll PJ, Zachariah Peery M, Berube M (2010) Detecting populations in the ‘ambiguous’ zone: kinship-based estimation of population structure at low genetic divergence. Molecular Ecology Resources, 10, 797–805.

Palstra, F.P., and D.J. Fraser, 2012 Effective/census population size ratio estimation: a compendium and appraisal. Ecology and Evolution 2:2357–2365.

Pollock, K.H., 2000. Capture-recapture models. Journal of the American Statistical Association, 95(449):293–296.

Punt, A.E., Dunn, A., Elvarsson, B.Þ., Hampton, J., Hoyle, S.D., Maunder, M.N., Methot, R.D. and Nielsen, A., 2020. Essential features of the next-generation integrated fisheries stock assessment package: A perspective. Fisheries Research, 229, p.105617.

Quinn, T.J. and Deriso, R.B., 1999. Quantitative fish dynamics. Oxford University Press.

Rawding DJ, Sharpe CS, Blankenship SM. 2014. Genetic-based estimates of adult Chinook salmon spawner abundance from carcass surveys and juvenile out-migrant traps. Transactions of the American Fisheries Society 143(1):55–67.

Robinson, J. D., and G. R. Moyer, 2013 Linkage disequilibrium and effective population size when generations overlap. Evol. Appl. 6:290–302.

Ruzzante, D.E., McCracken, G.R., Parmelee, S., Hill, K., Corrigan, A., MacMillan, J. and Walde, S.J., 2016. Effective number of breeders, effective population size and their relationship with census size in an iteroparous species, Salvelinus fontinalis. Proceedings of the Royal Society B: Biological Sciences 283:20152601.

Ruzzante, D.E., McCracken, G.R., Førland, B., MacMillan, J., Notte, D., Buhariwalla, C., Flemming, J.M. and Skaug, H. 2019. Validation of close-kin mark-recapture (CKMR) methods for estimating population abundance. Methods Ecol Evol. 10:1445–1453.

Santiago, E., Novo, I., Pardiñas, A.F., Saura, M., Wang, J. and Caballero, A., 2020. Recent demographic history inferred by high-resolution analysis of linkage disequilibrium. Molecular Biology and Evolution, 37(12), pp.3642–3653.

Seber, G. A. F. 1982. The estimation of animal abundance and related parameters. Macmillan, New York.

Seber, G.A., 1986. A review of estimating animal abundance. Biometrics 42:267–292.

Shaw, A. K., and S. A. Levin. 2013. The evolution of intermittent breeding. Journal of Mathematical Biology, 66:685–703.

Simpfendorfer C, Cox S, Stokes K and Waples R (2021) Independent Expert Peer Review of the Close Kin Mark Recapture Assessment for School Shark. Report to AFMA. AFMA Research Project Number 190844.

Skaug, H. J. (2001). Allele-sharing methods for estimation of population size. Biometrics 57 750–756.

Skaug, H.J., 2017. The parent–offspring probability when sampling age-structured populations. Theoretical population biology, 118, pp.20–26.

Stewart, J.D., Jaine, F.R., Armstrong, A.J., Armstrong, A.O., Bennett, M.B., Burgess, K.B., Couturier, L.I., Croll, D.A., Cronin, M.R., Deakos, M.H. and Dudgeon, C.L., 2018. Research priorities to support effective manta and devil ray conservation. Frontiers in Marine Science, 5:314.

Tallmon, D.A, D. Gregovich, R.S. Waples, C.S. Baker, J. Jackson, B. Taylor, E. Archer, K.K. Martien, and M.K. Schwartz. 2010. When are genetic methods useful for estimating contemporary abundance and detecting population trends? Molecular Ecology Resources 10:684–692.

Taylor, B.L. and Wade, P.R., 2000. “Best” Abundance Estimates and Best Management: Why They Are Not the Same. In Ferson S, Burgman M, eds, Quantitative methods for conservation biology (pp. 96–108). Springer, New York, NY.

Tenesa A, Navarro P, Hayes BJ et al. (2007) Recent human effective population size estimated from linkage disequilibrium. Genome Research 17:520–526.

Thompson, E.A., 2013. Identity by descent: variation in meiosis, across genomes, and in populations. Genetics, 194(2):301–326.

Thomson, R.B., Bravington M.V., Feutry, P., Gunasekera, R. and Grewe, P. (2020). Estimating the abundance of School Shark in Australia using close kin genetic methods. Final Report to FRDC Number 2014/024, August 2020. 108 pp. Available at: https://www.frdc.com.au/project/2014-024.

Wang, J., 2009 A new method for estimating effective population sizes from a single sample of multilocus genotypes. Molecular Ecology 18:2148–2164.

Wang, J., 2014. Estimation of migration rates from marker-based parentage analysis. Molecular Ecology, 23(13):3191–3213.

Wang, J., 2016. A comparison of single-sample estimators of effective population sizes from genetic marker data. Molecular Ecology, 25, 4692–4711.

Wang, J., Brekke, P., Huchard, E., Knapp, L.A. and Cowlishaw, G., 2010. Estimation of parameters of inbreeding and genetic drift in populations with overlapping generations. Evolution: International Journal of Organic Evolution, 64(6):1704–1718.

Waples, R.S. 2005. Genetic estimates of contemporary effective population size: To what time periods do the estimates apply? Molecular Ecology 14:3335–3352.

Waples, R.S. 2016. Life history traits and effective population size in species with overlapping generations revisited: the importance of adult mortality. Heredity 117:241–250.

Waples, R.S. 2020. An estimator of the Opportunity for Selection that is independent of mean fitness. Evolution 74:1942–1953.

Waples, RS. 2021. Relative precision of the sibship and LD methods for estimating effective population size with genomics-scale datasets. Journal of Heredity (published online 20 July 2021; DOI: 10.1093/jhered/esab042).

Waples, R.S., and C. Do. 2008. LDNE: A program for estimating effective population size from data on linkage disequilibrium. Molecular Ecology Resources 8:753–756.

Waples, R.S., and C. Do. 2010. Linkage disequilibrium estimates of contemporary Ne using highly variable genetic markers: A largely untapped resource for applied conservation and evolution. Evolutionary Applications 3:244–262.

Waples, R.S., C. Do, and J. Chopelet. 2011. Calculating Ne and Ne/N in age-structured populations: a hybrid Felsenstein-Hill approach. Ecology 92:1513–1522.

Waples, R.S., and P.R. England. 2011. Estimating contemporary effective population size based on linkage disequilibrium in the face of migration. Genetics 189:633–644.

Waples, R.S., G. Luikart, J.R. Faulkner, D.A. Tallmon. 2013. Simple life history traits explain key effective population size ratios across diverse taxa. Proc. Royal Society London, Ser. B. 280:20131339.

Waples, R.S., and T. Antao. 2014. Intermittent breeding and constraints on litter size: consequences for effective population size per generation (Ne) and per reproductive cycle (Nb). Evolution 68:1722–1734.

Waples, R.S., T. Antao, and G. Luikart. 2014. Effects of overlapping generations on linkage disequilibrium estimates of effective population size. Genetics 197:769–780.

Waples, R.S., and E.C. Anderson. 2017. Purging putative siblings from population genetic datasets: A cautionary view. Molecular Ecology 26:1211–1224.

Waples RS, Grewe PG, Bravington MV, Hillary R, Feutry P. 2018a. Robust estimates of a high Ne/N ratio in a top marine predator, southern bluefin tuna. Science Advances 4:eaar7759.

Waples RS, Scribner K, Moore J, Draheim H, Etter D, Boersen, M. 2018b. Accounting for age structure and spatial structure in eco-evolutionary analyses of a large, mobile vertebrate. Journal of Heredity 109:709–723.

Waples, R.S., Waples, R.K., and Ward, E.J., 2021. Pseudoreplication in genomics-scale datasets. Molecular Ecology Resources (published online 4 August 2021; DOI: 10.1111/1755-0998.13482).

Whiteley AR, Coombs JA, Hudy M, Robinson Z, Nislow KH, Letcher BH (2012) Sampling strategies for estimating brook trout effective population size. Conservation Genetics, 13, 625–637.

Whiteley AR, Coombs JA, Cembrola M, O’Donnell MJ, Hudy M, et al. 2015. Effective number of breeders provides a link between interannual variation in stream flow and individual reproductive contribution in a stream salmonid. Molecular Ecology 24:3585–602.

Williams BK, J. D. Nichols, and M. J. Conroy. 2002. Analysis and Management of Animal Populations. Academic Press, San Diego, CA, USA, 2002.

Wright, S., 1938. Size of population and breeding structure in relation to evolution. Science, 87:430–431.

