## Supplemental text, tables, and figures for "Close-kin methods to estimate census size and effective population size"

### Contents

S1 | PROBABILITY of a HALF-SIBLING MATCH
S2 | WORKED EXAMPLES
Tables and Figures
Computer code

### S1 | PROBABILITY of a HALF-SIBLING MATCH

The probability that two randomly selected offspring, one from time 0 and one from time *t*, have the same female parent is (from Equation 10 in main text):

$$P_{MHSP} = \sum_{i=1}^{N_{f0}} \left[\frac{k_{i,0}}{TRO_0} \times \frac{k_{i,t}}{TRO_t}\right] = \sum_{i=1}^{N_{f0}} \left[RRO_{i,0} \times RRO_{i,t}\right] \quad . \qquad \text{(S1)}$$

In a sibling-only design parents are not observed directly, so it is necessary to sum across all possible females that could have been alive when potential siblings were born. Females that mature after time 0 cannot produce a sibling match for offspring born in time 0, so the summation is Equation S1 is taken over the $N_{f0}$ females that were sexually mature at time 0.

Equation S1 can be reformulated by taking advantage of two mathematical identities. Let *X* and *Y* be vectors of real numbers, both of length *n*. Then,

- $\sum_{i=1}^{n}[X_i Y_i] = n * \overline{XY}$, and
- $\overline{XY} = \bar{X} * \bar{Y} + cov(X, Y)$.

Substituting for $X = RRO_0$, $Y = RRO_t$, and $n = N_{f0}$ leads to:

$$P_{MHSP} = N_{f0}[\overline{RRO_0} * \overline{RRO_t} + cov(RRO_{i,0}, RRO_{i,t})] \;.$$

At time 0, $TRO_0$ is computed across all $N_{f0}$ females, and because *RRO* is standardized such that $\Sigma(RRO_i) = 1$, it follows that $\overline{RRO_0} = 1/N_{f0}$. For time period *t*, however, the summation in Equation S1 only includes data for the subset of females that were mature at time 0, so we refer to this partial average as $\overline{RRO_{t*}}$. Making the substitution for $\overline{RRO_0}$ produces this general result that is mathematically equivalent to Equation S1:

$$\begin{aligned} P_{MHSP} &= N_{f0}[(1/N_{f0}) * \overline{RRO_{t*}} + cov(RRO_{i,0}, RRO_{i,t*})] \\ &= \overline{RRO_{t*}} + N_{f0}[cov(RRO_{i,0}, RRO_{i,t*})]. \end{aligned} \qquad \text{(S2)}$$

Equation S2 shows that the probability of a half sibling match can be expressed as a simple linear function of the covariance in individual *RRO* across time, with an intercept that is the partial mean *RRO* of adult females at time *t*. This demonstrates that cross-cohort siblings provide information about adult abundance at the time the younger sibling was born. A hypothetical worked example (Table S4) illustrates calculation of $P_{MHSP}$ using Equations S1 and S2.

### S2 | WORKED EXAMPLES

#### S2.1 | Simulations using CKMRPop

For each scenario in Table 3, we ran replicate simulations using CKMRPop, collected samples from specified years, and averaged results across replicates. To analyze POPs, each replicate ran for 25 years; in year 24 a sample of $n_A$ potential parents was drawn from the entire population of adults, and in year 25 another sample was randomly drawn of $n_{Off}$ yearling offspring produced by the previous year's parents. This process was repeated 500 times to generate replicate sets of data for estimating $N_A$ using CKMR. For siblings, we compared relative precision to estimate $N_A$ using CKMR to precision to estimate $N_b$ using Wang's sibship method. Each generation, true $N_b$ was calculated from the full pedigree using Equation 1 (main text). Each sibling replicate ran for 60 years, and samples of $n_{Off}$ offspring were collected for five consecutive years (years 51-55). Data were averaged over 100 replicates, producing a total of 500 replicate samples for analysis.

In the following examples, for simplicity we track the number of close-kin matches ($R$) involving female parents, but since our modeled populations have (on average) equal sex ratios, the total expected number of matches is twice the value for females. We show how various features of experimental design and species' biology affect probabilities of (and expected numbers of) close-kin matches and how to account for these effects to achieve an unbiased estimate of $N$.

### S2.2 | Calculation of the conversion factor

In the cartoon version of CKMR, development of the estimator of abundance is greatly simplified by two key assumptions: 1) all parents are equivalent in terms of producing offspring, so the probability that a random pair of individuals will produce a close-kin match is just $P_{match} = 1/N_f$; and 2) the pairwise comparisons are independent, so the expected number of matches in a particular study is just the product of the number of comparisons and the probability of success ($E(R) = P_{match}*Y$).

The first assumption is rarely true, so illustrating the consequences of unequal parental weights is the main goal of our simulations. The second assumption is not strictly true for either POPs or siblings, since the outcome of one sampling event affects the probability of subsequent events. For POPs, each offspring has exactly one mother. Strictly speaking, therefore, the series of comparisons of an offspring with different females to search for its mother are not independent Bernoulli trials with a fixed probability of success. Similarly, all sibling groups are of finite size, even in highly fecund species. This means that finding a sibling for individual X affects the probability that another individual will also be its sibling. This issue is clearly important for studying small populations, for which a substantial fraction of the population must be sampled to achieve reasonable precision. For relatively large populations, however, good precision can be achieved by sampling only a small fraction of the population (Figure 3B). Because of the relatively sparse sampling in these cases, the Bernoulli assumption provides a close approximation to the true expected number of successes across multiple trials (Bravington et al. 2016b), and that is the assumption we adopt here.

The fact that it is generally NOT the case that $P_{POP} = 1/N_f$ (and that it is NEVER the case that $P_{HSP} = 1/N_f$ for cross-cohort comparisons of offspring) complicates estimation of abundance. However, this can be dealt with by defining a conversion factor, $C$, which is the ratio of the true probability of a close-kin match to the naïve assumption of $1/N_f$:

$$C = P_{match}/(1/N_f) = N_f * P_{match}, \quad \text{so} \tag{S3}$$

$$P_{match} = C/N_f. \tag{S4}$$

If $Y$ is the total number of pairwise comparisons (which are treated as if they were independent), it follows that the expected number of close-kin matches ($R$) is

$$E(R) = Y * P_{match} = Y * C/N_f. \tag{S5}$$

The abundance estimator is then found by simple re-arrangement:

$\widehat{N}_f = Y*C/R.$ (S6)

The conversion factor thus plays the role of the function $f_2$ in Equation 7. Conceptually, $C$ can be thought of in two mathematically equivalent ways. First, it can be viewed as the factor by which the naïve probability of a match has to be multiplied to get the true $P_{match}$ (cf Equation S3). Alternatively, $C$ can be thought of as the factor by which $Y$ has to be multiplied to get the effective number of comparisons, after excluding those that cannot produce a match (cf Equation S5).

After obtaining the conversion factor $C$, estimating abundance is straightforward. But how does one find the value of $C$? In Equation S3, $C$ is defined as the ratio of the true probability of a close-kin match to the naïve probability = $1/N_f$. But $N_f$ is the quantity one wants to estimate! The solution to this conundrum is that, in our simple examples, the value of $C$ can be calculated independently of $N_f$, provided one knows or can estimate the population's vital rates—age-specific survival and/or fecundity, and (for POPs) age-specific selectivity in sampling adults. In a standard life table such as shown in Table 2, total $N$ and the age-specific values in the vector $N_x$ depend on the cohort size ($N_1$), but the relative numbers in each class depend only on age-specific survivals. As illustrated in the examples below, the vectors of age-specific survivals and fecundities can be used to calculate the conversion factor $C$, regardless how large or small $N_1$ or total $N$ are. Then, $C$ can be used, together with observed numbers of close-kin matches and known numbers of comparisons, to estimate $N_A$ as in Equation S6.

**S2.3 | POPs**

We can generalize POP-based CKMR by jointly considering the probabilities that different adults will a) produce offspring, and b) appear in the adult sample (see Table S1). We use the vital rates in Table 2, which has 1539 adult females aged 3-10. Initially we consider the following scenarios (see Table 3 for details), all of which involve annual random sampling of $n_{Off}$ = 300 offspring, annual sampling (random or selective) of $n_{Adult}$ = 300 adult females, and $\phi = 1$: Scenario A—constant fecundity with age and equiprobable sampling of adults; Scenario C—fecundity that is proportional to age and equiprobable sampling of adults; and Scenario D—fecundity and selective sampling probability that are both proportional to age. In these scenarios, relative fecundities and sampling probabilities of adult females of age $x$ are given by the vectors $\boldsymbol{w_{off}}$ and $\boldsymbol{w_{samp}}$, respectively, with $\boldsymbol{w_{off}}$ being identical to the relative parental weights defined in Section 2.2 and $\boldsymbol{w_{samp}}$ being the analogous index for sampling probability. For constant fecundity and equiprobable sampling, these weights are the same for all individuals; with increasing fecundity and selective sampling, the relative weights are 3:10 for ages 3:10 (Table 2). Within age classes, each individual is assumed to have the same weight. For $\boldsymbol{w_{off}}$ and $\boldsymbol{w_{samp}}$, the notation ($i$) indicates the value applies to an individual and ($x$) indicates the value is the sum across all the $N_x$ individuals within the respective age class.

Now consider a single event in which a randomly-sampled juvenile ($j$) is compared with a randomly-sampled adult female to see if they produce a POP match. The female could be any of the $i$ = 1 … 1539 mature individuals in the population. For each female $i$, we can compute the absolute probability that this female is the mother of offspring $j$ AND it is the one we have sampled. The first probability is the standardized parental weight $W_{off(i)} = w_{off(i)}/\Sigma w_{off(i)}$, which is also equal to $ERRO_i$. The second probability is the analogous standardized sampling weight, $W_{samp(i)} = w_{samp(i)}/\Sigma w_{samp(i)}$. The denominators of these standardized weights can be computed as the sum of the products of $N_x$ and $w_x$ for each age class. The joint probability we are interested in is the product of the two weighted probabilities: $W_{off(i)} * W_{samp(i)}$. The sum of these products across all adult females is the overall probability that, whoever the sampled female *i* turns out to be, she will be the mother of offspring *j*.

The example in Table S1 shows that, for Scenario D where both fecundity and the probability of an adult being sampled increase with age, $P_{POP}$ is higher (0.00075) than it would be under the naïve model (0.00065), so the adjustment factor is > 1 ($C$ = 0.00075/0.00065 = 1.16, accounting for rounding errors in reporting $P_{POP}$). It follows from Equation S5 that under this scenario, the expected number of POP matches is $E(R_{POP}) = C*Y_{POP}/N_f = C*n_{Off}*n_{Adult}/N_f$ = $1.16*300^2/1539 = 67.5$, whereas under either Scenario A or C, $E(R_{POP})$ is simply $300^2/1539$ = 58.5. Because the combination of selective sampling + increasing fecundity increases the expected number of parent-offspring matches by 16%, these factors would cause a corresponding downward bias in the estimate of $N_f$, unless they were accounted for as shown in Equation S6.

Three additional POP scenarios (all variants of Scenario A) are also shown in Table 3. Scenario B is identical to Scenario A except that variance in reproductive success is substantially overdispersed for each age and sex ($\phi$ = 10). In theory this should not affect the expected number of POPs but it should increase the variance among replicates. In Scenario E, the sample of potential parents is drawn from all individuals rather than just mature adults; because none of the juveniles can be the parent in a parent-offspring pair, using some of the 'adult' sample to include juveniles reduces $P_{POP}$ and the expected number of POPs, which in turn would increase the estimate of abundance unless accounted for. Scenario F differs from Scenario A only in having an order of magnitude larger cohort size (which also increases Adult *N* by an order of magnitude) and larger samples of adults and offspring.

### S2.4 | Siblings

**S2.4.1 | Effects of survival.** CKMR estimates based on siblings are generally restricted to comparisons of individuals from different cohorts to avoid the effective-population size signal that drives within-cohort sibling probabilities. It is therefore essential to account for mortality because cross-cohort siblings are impossible for parents that die between times 0 and *t*. In our example, we need to consider the probability that all adults alive at time 0 will still be alive $t$ = 1-4 years later. Two outcomes are possible for our modeled population. If the sum of current age ($x$) and $t$ exceeds $\omega$ (10 in our example), survival probability is 0 because all individuals die after age 10; otherwise, cumulate survival across $t$ years starting at age $x$ is:

$$s_{x\to x+t} = 0.7^t \text{ for } t \leq \omega\text{-}x \text{ and 0 otherwise,} \qquad \text{(S7)}$$

where $0.7^t$ = [0.7, 0.49, 0.343, 0.24] for $t$ = 1-4. Average survival across $t$ = 1-4 years for the entire adult population alive at time 0 is given by $[\sum_{x=t+3}^{10} N_x]/[\sum_{x=3}^{10} N_x]$. For our modeled population, the vector of cumulative survival rates for $t$ = 1-4 years is $\boldsymbol{s_{x\to x+t}}$ = [0.682, 0.459, 0.303. 0.194].

**S2.4.2 | Constant fecundity.** In Scenario A, all adults have the same expected fecundity. With $\phi$ = 1, each year all the adults behave like a single Wright-Fisher population with random variance in offspring number ≈ the mean, so $E(N_b) = N_A = 3078$. With random mating in this large a population, full siblings are rare, and almost all those that do occur will be found within a single cohort. With this simple life history, we can use the cartoon version of sibling CKMR (in which $P_{MHSP} = 1/N_f$), after accounting for mortality. Since all females are reproductively equivalent, the mortality-adjusted probability that a pair of randomly-chosen offspring born $t$ years apart share the same mother is just the product of $1/N_f$ and cumulative survival:

$$\boldsymbol{P_{MHSP(t)}} = \left(\frac{1}{N_f}\right)[0.682, 0.459, 0.303. 0.194] \qquad \text{for } t = 1\text{-}4$$
$$= [0.000443, 0.000298, 0.000197, 0.000126]. \qquad (S8)$$

For Scenario A, we can compute a vector of $C$ values for HSPs, one for each gap in years between cohorts, as $\boldsymbol{C_t} = \boldsymbol{P_{MHSP(t)}}/(1/N_f)$ = [0.000443, 0.000298, 0.000197, 0.000126]/0.00065 = [0.682, 0.459, 0.303. 0.194], which is just the vector of cumulative survivals for all adults.

With $n_{Off}$ offspring sampled each year there are $n_{Off}^2$ cross-cohort comparisons of individuals for each pair of yearly samples. In five years of samples, the total number of pairwise comparisons of samples is [4,3,2,1] for samples collected 1-4 years apart. In Scenario A with $n_{Off}$ = 100, the total number of pairwise comparisons of individuals to search for siblings is $\boldsymbol{Y_t}$ = [40000,30000,20000,10000] for age gaps of $t$ = 1-4 years, or 100K total. From Equation S5 we can calculate the expected numbers of HSP matches for age gaps of 1-4 years as $E(R_{MHSP(t)}) = \boldsymbol{C_t}*\boldsymbol{Y_t}/N_f$ = [40000,30000,20000,10000] [0.682, 0.459, 0.303, 0.194]/1539= [17.7, 8.9, 3.9, 1.3]. These are expected numbers of sibling matches for females; we expect an equal number for males, for a total of $E(R_{HSP(t)})$ = [35.4, 17.9, 7.9, 2.5] for $t$ = 1-4 years. One way to think about this is that adult mortality reduces the expected number of HSPs about 32%, 54%, 70%, and 81% for comparisons of offspring born 1-4 years apart.

We can also compute a weighted overall mean $C$ value for our experimental design, for which the weights (the relative number of among-year comparisons) are $\boldsymbol{y_t}$ = [4,3,2,1]. This weighted $C$ value is computed as $\bar{C} = \Sigma(\boldsymbol{C_t}*\boldsymbol{y_t}/\Sigma \boldsymbol{y_t}) = 0.4905$. For each replicate of 5 consecutive years of samples, therefore, we expect to find 49% as many HSPs as we would if mortality were not a factor. With 100K total comparisons possible to search for female HSPs, and an equal number to search for male HSPs, we expect to find a total of 200K*0.49/1539 = 63.7 HSPs.

#### S2.4.3 | Changes in fecundity with age

Fecundity increases with age in scenarios C and D, so it is no longer the case that $P_{MHSP}$ is the same for all adult females. Scenarios C and D differ only in selectivity of sampling adults, which does not affect the occurrence of siblings, so we focus on Scenario C. We start by computing $ERRO_i$ for each adult female alive at time 0. This is obtained from the $W_{off(i)}$ column in Table S1 (second and third sets of data), since those standardized parental weights are identical to $ERRO_i$. These values range from 0.00040 for age 3 to 0.00134 for age 10. Next, we can construct vectors of $ERRO_i$ values for the same individuals in future years that account for both mortality and changes in fecundity with age. Given our 5-year experimental design, we want to calculate $ERRO_{i(t)}$ for $t$ = 1-4 years in the future for the $N_f$ females alive at time 0. To illustrate, consider an individual of age 8 at time 0, whose *ERRO* is 0.00107 (Table S1); we want to find its *ERRO* $t$ years in the future, to calculate the probability that such an individual could produce siblings separated in birth by $t$ years. If $t$>2, the female would be beyond the maximum allowed age and its *ERRO* would be 0, so let's assume that $t$ = 2. The conditional *ERRO* for individuals aged 8+2 = 10 is 0.00134, conditional on surviving to that age. Because our focal individual has not yet survived to age 10, this conditional *ERRO* has to be discounted to account for mortality. Using the 2-year survival probability of $0.7^2 = 0.49$, the expected future relative reproductive output for the focal individual at age 10 is 0.001342*0.49 = 0.00066.

To calculate $P_{MHSP}$ as in Equation S1, the summation of products has $N_f$ terms (one for each female alive at time 0). Like Table S1, Table S2 is condensed to have one row per adult age class. The first three columns in Table S1 give age, number of females in each age class, and

$b_x$ for each age class. Column 4 has age-specific *ERRO* values (taken from Table 2) for each individual. The next 4 columns have the adjusted *ERRO* values for each individual 1-4 years into the future. From this matrix, the probability that a randomly selected pair of offspring, one born in year 0 and one born in year $t$, will share the same mother can be calculated from Equation S1 by pairwise combinations of column 4 with the next 4 columns. The last column in Table S1 shows how to calculate $P_{MHSP}$ for siblings separated by one birth year, using the sums-of-products approach in Equation S1. For example, at time 0 there were $N_5$ = 240 age-5 females, each with *ERRO* = 0.00067, and their conditional *ERRO* a year later was 0.00056. Across these females, the sum of products of *ERRO* now and one year in the future is 240*0.00067*0.00056 = 0.000091, as shown in the last column. The sum of these age-class products across all adult ages is 0.000564, which is the overall probability of a MHSP match for offspring born in consecutive years. The same result can be obtained using the modified Equation S2. $\overline{ERRO}_{t=1}$ (that is, the age-weighted mean of the 5th column in Table S2) is 0.000522 and the covariance of the full columns 4 and 5 across all 1539 females alive at time 0 is $2.705x10^{-8}$, so $P_{MHSP(t=1)}$ = 0.000522 + $1539*2.7x10^{-8}$ = 0.000564.

This probability is lower than $1/N_f$ = 0.00065, but it is higher than $P_{MHSP}$ that only accounts for mortality (0.000443 for $t$=1 from Equation S8), which indicates that consistent patterns in age-specific variation in fecundity also affect $P_{MHSP}$. In this example, these patterns increase $P_{MHSP}$, but not enough to fully offset the reductions associated with mortality. This value applies to comparisons 1 year apart. Similar calculations for the other 'next year' columns in Table S2 (data not shown) produce this vector of $\boldsymbol{P_{MHSP(t)}}$ that accounts for both mortality and changes in fecundity with age:

$$\boldsymbol{P_{MHSP(t)}}[t=1\text{-}4] = [0.000564, 0.000406, 0.000280, 0.000182] \;. \qquad \text{(S9)}$$

From this vector and the vector of relative numbers of comparisons for each gap in years ($\boldsymbol{y_t}$ = [4,3,2,1]), we can calculate an overall $\bar{P}_{MHSP}$ for our experimental design that involves comparisons across 5 years of samples as overall $\bar{P}_{MHSP}$ = [4*0.000564 + 3*0.000406 + 2*0.000280 + 0.000182]/10 = 0.000421.

Given $P_{HSP}$, it is straightforward to calculate the expected number of sibling matches. As with Scenario A and constant fecundity, we define a vector of $C$ values such that $\boldsymbol{C_t}$ = $\boldsymbol{P_{HSP(t)}}/(1/N_f) = N_f * \boldsymbol{P_{HSP(t)}}$. For example, in Scenario C (increasing fecundity with age) and for cohorts sampled one year apart, $\boldsymbol{P_{HSP(1)}}$ = 0.000564 (Table S2) and $1/N_f$ = 0.00065, so $C_1$ = 0.000564/0.00065 = 1539*0.000564 = 0.868. This indicates that, under this scenario, comparing random pairs of offspring born 1 year apart is expected to produce 86.8% as many sibling pairs as would occur if the overall match probability were $1/N_f$. Each comparison across years involves $(n_{off})^2 = 100^2$ = 10K pairwise comparisons of individuals and there are 4 comparisons of samples collected one year apart, so the total number of comparisons for $t$ = 1 is 40K and the expected number of sibling matches is $(C_1/N_f))$*40K = 0.868*40K/1539 = 22.56 pairs that share a female parent, or 45.1 that share either or both parents. Similar calculations for other age gaps lead to expected numbers of HSPs (maternal and paternal combined) for gaps of 2-4 years as [24.4, 11.2, 3.6], for a total of 84.3 expected matches across all comparisons. We can check this against the weighted mean $\bar{P}_{MHSP}$ = 0.000421; with 100K comparisons, the expected number of matches is 42.1 for females or 84.2 for both sexes (with difference from 84.3 due to rounding). The overall $\bar{C}$ value for HSPs that accounts for both survival and changes in fecundity with age is 0.000421/0.00065 = 0.648—that is, the average comparison of two offspring is just under 65% as likely to produce a sibling match as it would be if all parents were equally likely to produce offspring and there were no mortality between years.

### S2.4.4 | Intermittent breeding

We conducted additional simulations in which only subsets of mature adults were allowed to reproduce each year, and we monitored the consequences for the distribution of siblings as a function of their age gap. Except for the intermittent breeding, these simulations followed Scenario A. In the '50%' and '10%' scenarios, only half or 10% of the 3078 mature adults, respectively, were allowed to reproduce, and selection of the lucky parents was done randomly and independently across years. In the 'Alternate' scenario, only adults aged 3,5,7, or 9 were allowed to reproduce each year, with the consequence that surviving adults in a given year were allowed to reproduce only if they did not the year before.

### S2.4.5 | Estimation of $N_b$

For iteroparous species, Wang's sibship method for estimating $N_b$ is restricted to comparisons among individuals from a single cohort (age gap = $t$ = 0). For $t$ = 0, and assuming all $\phi = 1$, Equation S4 reduces to $P_{MHSP(within)} = 1/N_f$. Each of the five samples can be used to estimate $N_b$; with $n_{Off}$ offspring there are $n_{Off}\,(n_{Off}-1)/2 \approx n_{Off}^2/2$ pairwise comparisons of individuals possible each year, and each comparison has two chances to yield a sibling match: maternal and/or paternal. In our example $N_f = N_m = 1539$, so with $n_{Off} = 100$ the expected total number of sibling matches per year is $2*[n_{Off}\,(n_{Off}-1)/2]*(1/N_f) = 9900/1539 = 6.4$, or 32.2 combined over 5 years. When variance in reproductive success is substantially overdispersed ($\phi$ = 10 for each age and sex), overall $N_b$ is reduced almost 90% to 388 (194 each for males and females; AgeNe). It is straightforward to recalculate sibling probabilities using $P_{HSP(within)} = 1/N_{bf}$ and $1/N_{bm}$, and doing this greatly increases the expected number of sibling matches to 51 per year or 255 across 5 years (Table S3).

Changes in fecundity with age increase the overall variance in reproductive success among all adults in a single year, and this reduces $N_b$ and increases $P_{HSP}$. For Scenario C with $\phi$ = 1, age-specific variation in fecundity reduces $N_b$ to 2665 (1333 for each sex), and the resulting expected number of sibling matches is 3.7 per sex per year, or 37.1 total over the five years. With $\phi$ = 10 $N_b$ again is greatly reduced, but only slightly (to 381) compared to Scenario A with $\phi$ = 10 ($N_b$ = 388). For scenario B with overdispered RS, the expected numbers of within cohort sibs are 52 per year and 260 total across 5 years.

### S2.5 | Precision

Because of the inverse relationship between $\widehat{N}$ and $R$, even if the observed number of close-kin matches is approximately normally distributed (as will often be the case), $\widehat{N}$ will not be; instead, $\widehat{N}$ is skewed high and is infinitely large when no recoveries are found. This same issue applies to Wang's sibship method (see Equation 4), and for this reason it is common for evaluations of precision of effective size estimators to focus on the distribution of $1/\widehat{N}_e$ rather than $\widehat{N}_e$ (e.g., Wang 2001; 2009; Waples and Do 2010). Accordingly, here we report values for the CV of $1/\widehat{N}$ and $1/\widehat{N}_b$ rather than $\widehat{N}$ and $\widehat{N}_b$. When precision is high, CV($\widehat{N}$) and CV($1/\widehat{N}$) are generally very similar, but CV($\widehat{N}$) can be much larger when precision is low and is undefined if any $\widehat{N} = \infty$. In contrast, $1/\infty = 0$ so these outcomes pose no problem for computing CV($1/\widehat{N}$). Mathematically, using Equation S6, CV($1/\widehat{N}$) is identical to CV($R$) = the coefficient of variation of the number of close-kin matches.

### S3 | Simulation Results

Simulations for POPs and siblings had roughly equivalent amounts of replication. For POPs, samples of adults (in year 24) and their yearling offspring (in year 25) occurred once in each of 500 replicates. For siblings, in each of 100 replicates samples of offspring were taken from 5 consecutive years, for 500 total samples. Under all scenarios considered in Table 3, the mean total numbers of POPs and HSPs found agreed closely with the expected numbers, which were computed using adjusted $P_{POP}$ and $P_{HSP}$ values that accounted for effects of covariates (Table 3). In addition, the numbers of sibling matches found for different age gaps between samples agreed with our predictions (Table S3). Below we consider in more detail results for POPs and siblings separately.

#### S3.1 | POPs

For POPs, raw harmonic mean estimates of adult $N$ (based on the naïve estimator $\widehat{N}_A = Y/R$) were unbiased only when the conversion factor $C = N_f * P_{POP}$ was exactly 1.0, indicating that the true probability of a close-kin match was $1/N_f$ (as in POP scenarios A, B, C, and F). In scenario D the fact that fecundity and adult selectivity both increased with age increased $P_{POP}$ from the naïve expectation of $1/N_f$, and this produced an excess of parent-offspring matches compared to the naïve expectation, with the consequence that the naïve estimator $\widehat{N}_A = Y/R$ was downwardly biased by 9% (2802 vs 3078). Using the adjusted estimator $\widehat{N}_{A(adj)} = C*Y/R$ brought the adjusted estimate to 3094, within 0.5% of the true value (Table 3).

As expected, overdispersed variance in reproductive success ($\phi = 10$) for Scenario B did not bias the POP-based abundance estimate, but it did increase the variance (Figure 2). When the entire population was sampled for potential parents (Scenario E), the dilution effect of including comparisons with nonreproducing individuals reduced the number of POPs and greatly increased the raw abundance estimate (by over 100%, to 6722). If the objective was to estimate total population size of age 1+ individuals ($N_T = 6478$), this approach would be appropriate. However, if the goal was to estimate adult abundance, this would represent a substantial upward bias, which could be largely eliminated by adjusting the raw $\widehat{N}_A$ using the conversion factor ($C = 0.48$), leading to $\widehat{N}_{A(adj)} = 3201$. Using an order of magnitude larger cohort size (Scenario F) in a simulation that otherwise was identical to Scenario A produced a ten-fold larger estimate ($\widehat{N}_A = 32256$) that was within a few percent of the true value.

We analyzed POPs for one additional scenario not described in Table 3: Scenario G followed Scenario A except that only age-10 individuals reproduced each year, and only age-10 individuals were sampled as potential parents. Scenario G thus represented an extreme form of selectivity for both reproduction and sampling: on average only 80 total 10-year olds reproduced randomly each year, and the adult sample was also drawn from that group (sample size of potential parents was reduced from 300 to 50 to reflect the reduction in available targets for sampling). With $1/N_f = 1/N_m = 1/40$ rather than 1/1539, $P_{POP}$ (calculated using the method illustrated in Table S1) was much higher, leading to an extreme conversion factor $C = 38.1$. For the modified sampling design, the expected number of POPs (from Equation S5) was 371.8 and an average of 375.7 were observed in the simulations. Using the number of matches and comparisons in the naïve estimator $\widehat{N}_A = Y/R$ would lead to $\widehat{N}_A = 81$, which in this case accurately reflected the number of reproducing adults each year, but the adjusted estimator which uses the conversion factor (Equation S6) leads to $\widehat{N}_{A(adj)} = 3072$, very close to the true number of age 3+ adults in the population.

Because only 10-year olds reproduce, no cross-cohort siblings are possible in this effectively-semelparous example, so sib-based estimates of adult *N* are not feasible. Within-cohort siblings were used to estimate $N_b$ each year using Wang's method, and the harmonic mean $\widehat{N}_b$ across replicates was 81, in close agreement with the mean annual number of randomly-reproducing parents

**S 3.2 | Siblings**

Because sib-based CKMR estimates of *N* are restricted to cross-cohort comparisons, it is always necessary to account for mortality. This means that the conversion factor will never be exactly 1.0 (unless by chance other factors cause an adjustment in the opposite direction that exactly cancels the effects of mortality). In our experimental design, the age gap for CKMR-based sibling estimates ranged from 1 to 4 years; cumulative survivals differ across those time intervals, which means that separate conversion factors apply to each age gap. Table 3 shows the weighted mean $\bar{C}$ values that account for all comparisons involving 5 consecutive years of samples, and these mean $\bar{C}$ values ranged from 0.49 to 0.65, depending on which factors affected the probability of a close-kin match. After accounting for effects of mortality and (as needed, for Scenario C) changes in fecundity with age, all adjusted estimates of *N* were within a few percent of the true values (Table 3).

The core scenarios evaluated by our simulations considered adult population sizes of ~3000, with one scenario (F) being ten times as large. Many marine (and some terrestrial) populations can be substantially larger, so we also evaluated the scale of sampling effort that would be required to produce precise estimates of *N* for a wide range of population sizes. For a given value of *N* (and given the life history features of our Scenario A), we calculated the annual sample size that, when used in a 5-year sampling regime as modeled here, could be expected to produce either 50 total cross-cohort siblings. These annual sample sizes are shown in the top panel of Figure 3 (and in Table S5) and range from ~45 per year when adult *N* is 770 to about 2800 when adult *N* was ~ $3x10^6$. Because the sample size required to produce a fixed level of precision increases only as the square root of $\widehat{N}_A$, the fraction of the population that has to be sampled is much lower when $\widehat{N}_A$ is large (Figure 3B).

Figure S1 plots the mean (+/- 1 sd) observed number of sibling matches for these simulations using a wide range of adult *N* values. Sample sizes were chosen to produce expected numbers of 50 siblings, and means across all replicates were very close to those predicted values. Although most of our other examples used a relatively small population (adult *N* = 3078), these results illustrate that unbiased estimates can also be obtained for large populations. Notably, all of these predicted numbers of siblings were made based on the conversion factor *C* calculated using the method illustrated in Table S2, for which adult *N* = 3078. This illustrates the point that the value of *C* depends on age-specific vital rates but not on cohort size or total population size.

For the scenarios considered in Figure S2A, the ratio var(*R*)/mean(*R*) was generally slightly higher for sibling analyses than for POPs. For scenarios B ($\phi$=10) and C (fecundity increasing with age), var(*R*)/mean(*R*) for siblings was > 1 (2.22 and 1.23, respectively), which presumably reflected the fact that both of these scenarios included unequal parental weights.

Our simulations of populations with large $N_A$ produced results consistent with the expectation (Bravington et al. 2016b) that var(*R*) should converge on the Poisson variance as abundance increases and sampling becomes more sparse compared to population size. For the smallest populations we modeled ($N_A$ = 770), var(*R*)/mean(*R*) was somewhat elevated (1.26) from the Poisson expectation of 1.0 even for $\phi$=1, but the variance rapidly dropped to at or below the Poisson variance as population size increased (Figure S2B). With moderately high within-age variance in reproductive success (ϕ=10), departures from the Poisson expectation are greater and the rate of attenuation lower, but even so var($R$)/mean($R$) dropped to ~1.0 for adult abundances > $10^5$ (Figure S2B). All of these scenarios were designed to produce an average of 50 total half-sibling matches, and those that achieved a var($R$)/mean($R$) ratio close to 1.0 produced CVs of $1/\hat{N}_A$ that were close to the expected 0.15 (Figure 3C).

For two scenarios (A and B), we compared precision for estimating $N_A$ using sib-based CKMR with precision to estimate $N_b$ using Wang's sibship method, as a function of the number of consecutive years of samples that were used (2-5). In Scenario A, ϕ = 1 and fecundity was constant with age, so $N_A = N_b = 3078$; precision was higher for $\hat{N}_b$ with only 2 years of data, but for longer studies precision of $\hat{N}_A$ was higher (Figure 6). This latter result occurred because the total number of pairwise comparisons increases faster for cross-cohort comparisons used in CKMR than it does for within-cohort comparisons used to estimate $N_b$. Results for Scenario B were quite different: with ϕ = 10, $N_b$ (388) is greatly reduced compared to adult $N$, and this produces a stronger drift signal that is relatively easier to estimate precisely. Conversely, the high variance in reproductive success substantially increased variability in the number of cross-cohort sibling matches. As a consequence, CV($\widehat{1/\hat{N}_A}$) was more than 5 times as large as CV($1/\hat{N}_b$) for 2 years of samples and still more than twice as large with 5 years of samples (Figure 6).

In the skip-breeding scenarios in which only a random portion of adults were allowed to reproduce each year, the resulting distributions of HSPs by age gap was essentially unchanged from that found for Scenario A (Figure 4). We believe this occurred because the lucky parents were chosen independently each year, so any correlations of individual ERRO across years were random. This would cause the covariance term in Equation S2 to vary randomly around 0 and hence have no systematic effect on the number of HSPs, except to increase the variance. Results for the 'Alternate' scenario were very different: in this case, no individuals reproduce in two consecutive years, and (apart from data errors) it is impossible to find any siblings with odd-numbered age gaps. This creates dramatic differences in the incidence of siblings as a function of the age gap, which means that the pattern of skip breeding must be accounted for to achieve an unbiased estimate of abundance.

Because sibship estimates of $N_b$ rely only on within-cohort comparisons of individuals, the pattern of intermittent breeding does not bias the estimates. Estimates of $N_b$ reflect the effective number of parents that actually reproduce each year, which (for the three skip-breeding scenarios depicted in Figure 4) are 50%, 10%, and 59% of the total of 3078 adults (data not shown).

Table S1. Example illustrating computation of the probability of a parent-offspring match ($P_{POP}$), using variations of the vital rates in Table 2. For simplicity, we consider only adult females and ignore age classes 1&2, which do not produce offspring. Columns with a small '*w*' show relative weights for producing offspring ('*off*') and sampling probability ('*samp*'); columns with an upper-case '*W*' show absolute weights [$W_i = w_i/\Sigma w_i$]. Subscripts '*i*' show weights for individuals and subscripts '*x*' show overall weights for each age class (e.g., $w_{off(x)} = N_x*w_{off(i)} = N_x*b_x$). Within age classes, all individuals have the same weight. The last two columns show the absolute joint probabilities of parenthood and being sampled for individuals ($W*W(i) = W_{off(i)}*W_{samp(i)}$) and for each age class ($W*W(x) = N_x*W*W(i)$). $P_{POP}$ is then calculated as $\Sigma(W*W(x))$. Calculations are shown for three reproductive/sampling scenarios described in Table 3: A = constant fecundity & equiprobable sampling; C = increasing fecundity with age and equiprobable sampling; D = fecundity and sampling probability both increase with age. Note that under the naïve 'cartoon' model, $P_{POP} = 1/N_f = 1/1539 = 0.00065$.

| | **Age (x)** | $N_x$ | $w_{off(i)}$ | $w_{off(x)}$ | $W_{off(i)}$ | $w_{samp(i)}$ | $w_{samp(x)}$ | $W_{samp(i)}$ | $W*W(i)$ | $W*W(x)$ |
|---|---|---|---|---|---|---|---|---|---|---|
| | 3 | 490 | 1 | 490 | 0.00065 | 1 | 490 | 0.00065 | 4.2E-07 | 0.00021 |
| | 4 | 343 | 1 | 343 | 0.00065 | 1 | 343 | 0.00065 | 4.2E-07 | 0.00014 |
| **A** | 5 | 240 | 1 | 240 | 0.00065 | 1 | 240 | 0.00065 | 4.2E-07 | 0.00010 |
| | 6 | 168 | 1 | 168 | 0.00065 | 1 | 168 | 0.00065 | 4.2E-07 | 0.00007 |
| | 7 | 118 | 1 | 118 | 0.00065 | 1 | 118 | 0.00065 | 4.2E-07 | 0.00005 |
| | 8 | 82 | 1 | 82 | 0.00065 | 1 | 82 | 0.00065 | 4.2E-07 | 0.00003 |
| | 9 | 58 | 1 | 58 | 0.00065 | 1 | 58 | 0.00065 | 4.2E-07 | 0.00002 |
| | 10 | 40 | 1 | 40 | 0.00065 | 1 | 40 | 0.00065 | 4.2E-07 | 0.00002 |
| | $N_A$ | 1539 | *TRO* | 1539 | | $\Sigma w_{samp(x)}$ | 1539 | | $P_{POP}$ | 0.00065 |
| | 3 | 490 | 3 | 1470 | 0.00040 | 1 | 490 | 0.00065 | 2.6E-07 | 0.00013 |
| | 4 | 343 | 4 | 1372 | 0.00054 | 1 | 343 | 0.00065 | 3.5E-07 | 0.00012 |
| **C** | 5 | 240 | 5 | 1201 | 0.00067 | 1 | 240 | 0.00065 | 4.4E-07 | 0.00010 |
| | 6 | 168 | 6 | 1008 | 0.00080 | 1 | 168 | 0.00065 | 5.2E-07 | 0.00009 |
| | 7 | 118 | 7 | 824 | 0.00094 | 1 | 118 | 0.00065 | 6.1E-07 | 0.00007 |
| | 8 | 82 | 8 | 659 | 0.00107 | 1 | 82 | 0.00065 | 7.0E-07 | 0.00006 |
| | 9 | 58 | 9 | 519 | 0.00121 | 1 | 58 | 0.00065 | 7.8E-07 | 0.00005 |
| | 10 | 40 | 10 | 404 | 0.00134 | 1 | 40 | 0.00065 | 8.7E-07 | 0.00004 |
| | | | *TRO* | 7456 | | $\Sigma w_{samp(x)}$ | 1539 | | $P_{POP}$ | 0.00065 |
| | 3 | 490 | 3 | 1470 | 0.00040 | 3 | 1470 | 0.00040 | 1.6E-07 | 0.00008 |
| | 4 | 343 | 4 | 1372 | 0.00054 | 4 | 1372 | 0.00054 | 2.9E-07 | 0.00010 |
| **D** | 5 | 240 | 5 | 1201 | 0.00067 | 5 | 1201 | 0.00067 | 4.5E-07 | 0.00011 |
| | 6 | 168 | 6 | 1008 | 0.00080 | 6 | 1008 | 0.00080 | 6.5E-07 | 0.00011 |
| | 7 | 118 | 7 | 824 | 0.00094 | 7 | 824 | 0.00094 | 8.8E-07 | 0.00010 |
| | 8 | 82 | 8 | 659 | 0.00107 | 8 | 659 | 0.00107 | 1.2E-06 | 0.00009 |
| | 9 | 58 | 9 | 519 | 0.00121 | 9 | 519 | 0.00121 | 1.5E-06 | 0.00008 |
| | 10 | 40 | 10 | 404 | 0.00134 | 10 | 404 | 0.00134 | 1.8E-06 | 0.00007 |
| | | | *TRO* | 7456 | | $\Sigma w_{samp(x)}$ | 7456 | | $P_{POP}$ | 0.00075 |

Table S2. Calculation of the probability of obtaining a maternal half-sibling pair ($P_{MHSP}$) by accounting for both mortality and changes in fecundity with age. This example applies to Scenario C from Table 3. *ERRO* for individuals by age is taken from Table 2, and juveniles (age classes 1-2) are omitted. For an individual of age *x* alive at time 0, *ERRO* $t$=1-4 years in the future is the product of two numbers: 1) *ERRO* for individuals that survive to age $x + t$ (from the appropriate column labeled 'ERRO in future years'); and 2) probability of surviving *t* years ($0.7^t$ or 0, depending on whether $x+t > 10$). From Equation S1, $P_{MHSP(t)} = \Sigma[ERRO_0 * ERRO_t]$. The last column 'Product' shows this summation for each age class and the total is the overall probability of a MHSP when comparing offspring born one year apart.

| | | | | ERRO in future years (*t*) | | | | |
|---|---|---|---|---|---|---|---|---|
| **Age(*x*)** | $N_x$ | $b_x$ | **ERRO** | **1** | **2** | **3** | **4** | **Product$_{t=1}$** |
| 3 | 490 | 3 | 0.00040 | 0.00038 | 0.00033 | 0.00028 | 0.00023 | 0.000074 |
| 4 | 343 | 4 | 0.00054 | 0.00047 | 0.00039 | 0.00032 | 0.00026 | 0.000086 |
| 5 | 240 | 5 | 0.00067 | 0.00056 | 0.00046 | 0.00037 | 0.00029 | 0.000091 |
| 6 | 168 | 6 | 0.00080 | 0.00066 | 0.00053 | 0.00041 | 0.00032 | 0.000089 |
| 7 | 118 | 7 | 0.00094 | 0.00075 | 0.00059 | 0.00046 | 0 | 0.000083 |
| 8 | 82 | 8 | 0.00107 | 0.00084 | 0.00066 | 0 | 0 | 0.000075 |
| 9 | 58 | 9 | 0.00121 | 0.00094 | 0 | 0 | 0 | 0.000065 |
| 10 | 40 | 10 | 0.00134 | 0 | 0 | 0 | 0 | 0.000000 |
| | | | | | | | $P_{MHSP(1)}$ | 0.000564 |

Table S3. Predicted number of HSPs (maternal and paternal combined) as a function of the gap in ages between offspring and the magnitude of variation in reproductive success ($\phi = 1$ or 10). Scenarios are described in Table 3 except Scenario C*, which is Scenario C with $\phi = 10$. The column for age gap = 0 applies to within-cohort comparisons. The last column ('Total') is the sum for all cross-cohort comparisons (age gaps 1-4). Predicted numbers of HSPs do not depend on $\phi$ for cross-cohort comparisons; for within-cohort comparisons, the first predicted number applies to $\phi = 1$ and the second to $\phi = 10$. Observed values are means across replicate simulations.

| | | | | **Age Gap** | | | |
|---|---|---|---|---|---|---|---|
| **Scenario** | | **0** | **1** | **2** | **3** | **4** | **Total** |
| **A** | **Observed, $\phi = 1$** | 29.9 | 34.8 | 17.4 | 8.6 | 3.0 | 63.8 |
| **B** | **Observed, $\phi = 10$** | 251.3 | 37.4 | 18.4 | 9.2 | 2.7 | 67.7 |
| | **Predicted** | 32.2/255 | 35.4 | 17.9 | 7.9 | 2.5 | 63.7 |
| **C** | **Observed, $\phi = 1$** | 36.8 | 46.2 | 24.6 | 10.5 | 3.7 | 85.0 |
| **C*** | **Observed, $\phi = 10$** | 256.4 | 43.4 | 25.2 | 11.8 | 2.8 | 83.2 |
| | **Predicted** | 37.1/260 | 45.1 | 24.4 | 11.2 | 3.6 | 84.3 |

Table S4. Hypothetical example illustrating calculation of the probability that comparing two randomly-selected offspring will produce a maternal half-sibling pair [$P_{MHSP}$]. Sibling probabilities are calculated using Equations S1 and S2 and data on the number of offspring ($k$) produced by females in two consecutive years (0, 1). The ten females alive at time 0 (1-10) are divided into four age classes (1-4), and four of the females die before year 1 (indicated by an ‘X’ for Age). In year 1, four new age-1 females recruit to the population (shaded cells). In each year, total reproductive output (*TRO*) is the sum of all offspring produced, and this is used to calculate relative reproductive output of each individual (for each time period, $RRO_i = k_i/TRO$). Although offspring produced by the new recruits in year 1 contribute to $TRO_1$, calculation of $P_{MHSP}$ only uses *RRO* data for females alive at year 0. The last column shows the products ($\Pi_i$) of $RRO_i$ for the two time periods, and the sum of these products for the 10 females alive at year 0 is the result obtained using Equation S1 [$P_{MHSP(S1)}$ = 0.0731]. Equation S2 uses the partial mean $RRO_i$ in year 1 for the 10 females alive in year 0 (0.089) and the covariance of $RRO_i$ in the two time periods ($cov$ = -0.0015) to obtain the identical result [$P_{MHSP(S2)} = 0.089 + 10{*}cov$ = 0.0731].

| | **Year 0** | | | **Year 1** | | | |
|---|---|---|---|---|---|---|---|
| **Female** | **Age** | ***$k_i$*** | ***$RRO_i$*** | **Age** | ***$k_i$*** | ***$RRO_i$*** | $\Pi_i$ |
| 1 | 1 | 2 | 0.059 | X | 0 | 0 | 0 |
| 2 | 1 | 1 | 0.029 | 2 | 5 | 0.143 | 0.0042 |
| 3 | 1 | 0 | 0.000 | 2 | 2 | 0.057 | 0.0000 |
| 4 | 1 | 3 | 0.088 | 2 | 3 | 0.086 | 0.0076 |
| 5 | 2 | 4 | 0.118 | X | 0 | 0 | 0 |
| 6 | 2 | 1 | 0.029 | 3 | 4 | 0.114 | 0.0034 |
| 7 | 2 | 3 | 0.088 | 3 | 8 | 0.229 | 0.0202 |
| 8 | 3 | 7 | 0.206 | X | 0 | 0 | 0 |
| 9 | 3 | 5 | 0.147 | 4 | 9 | 0.257 | 0.0378 |
| 10 | 4 | 8 | 0.235 | X | 0 | 0 | 0 |
| *11* | X | NA | NA | 1 | 2 | | |
| *12* | X | NA | NA | 1 | 0 | Sum($\Pi$) | 0.0731 |
| *13* | X | NA | NA | 1 | 1 | $P_{MHSP(S1)}$ | 0.0731 |
| *14* | X | NA | NA | 1 | 1 | | |
| ***TRO*** | | 34 | | | 35 | | |
| | | | **Year 0** | **Year 1** | | ***cov*** | -0.0015 |
| | **Mean** | ***$RRO_i$*** | 0.100 | 0.089 | | | |

Table S5. Detailed information regarding costs of CKMR shown in Figure 5D. Costs are in $USD and are shown for two options: 1) Dartseq only; 2) SNP discovery + dartseq. n = annual sample size of individuals; 5*n is total sample size for a 5-year experimental design. Each plate holds 94 samples + 2 controls; n' is the total number of samples that can be analyzed using all available wells on all plates and is the basis for cost estimates. Option 1 has a constant cost of $35/individual. In Option 2, SNP discovery costs a fixed $3290, after which costs per individual are reduced. Bold font highlights the option with smaller costs for each annual sample size.

| | | | | Option 1 | Option 2 | | | |
|---|---|---|---|---|---|---|---|---|
| n | 5*n | Plates | n' | Dartseq only | SNP discovery (dartseq) | panel synthesis | genotyping | Total |
| 44 | 220 | 3 | 282 | **9870** | 3290 | 5500 | 4230 | 13020 |
| 89 | 445 | 5 | 470 | 16450 | 3290 | 5500 | 7050 | **15840** |
| 177 | 885 | 10 | 940 | 32900 | 3290 | 7500 | 14100 | **24890** |
| 354 | 1770 | 19 | 1786 | 62510 | 3290 | 7500 | 26790 | **37580** |
| 709 | 3545 | 38 | 3572 | 125020 | 3290 | 10000 | 53580 | **66870** |
| 1418 | 7090 | 76 | 7144 | 250040 | 3290 | 15000 | 107160 | **125450** |
| 2836 | 14180 | 151 | 14194 | 496790 | 3290 | 20000 | 212910 | **236200** |

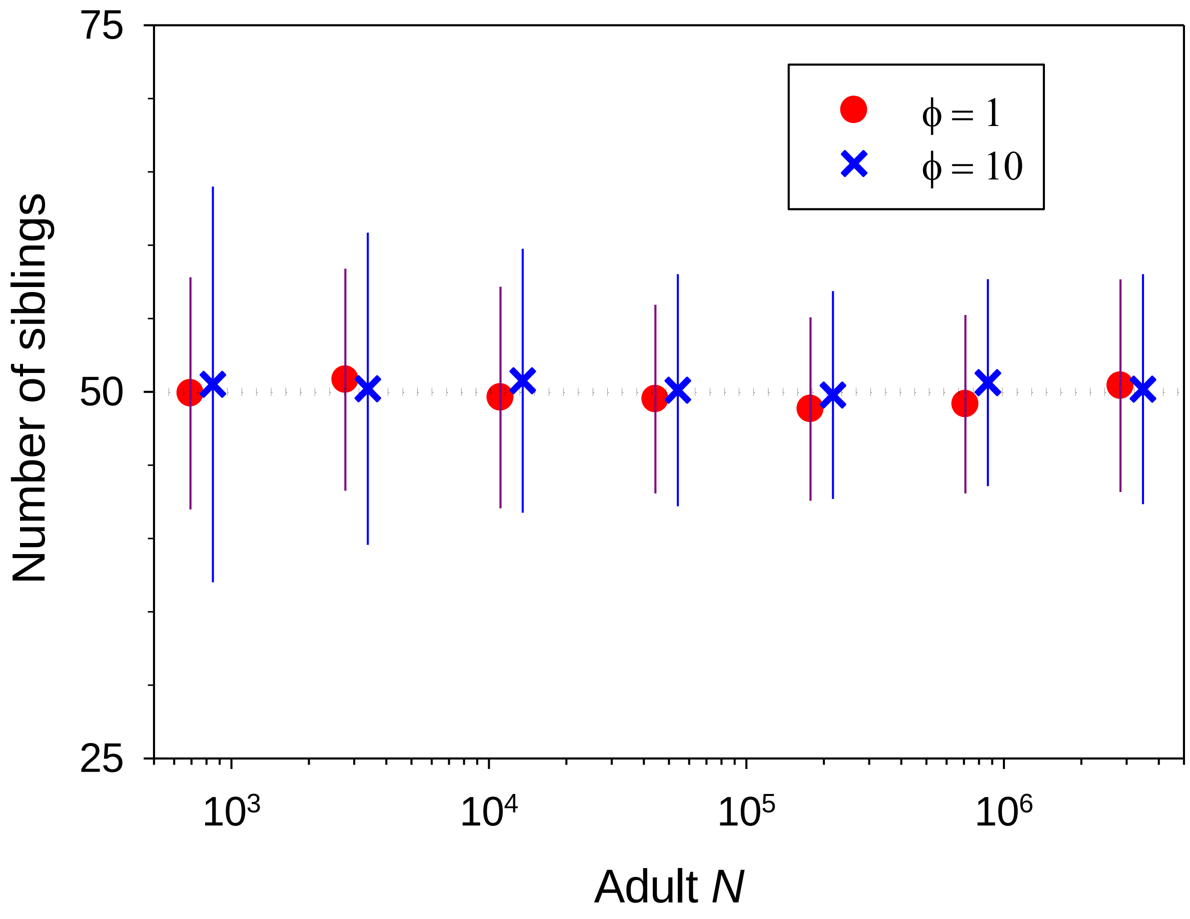

Figure S1. Mean (filled symbols) +/- one sd (vertical lines) of the number of cross-cohort siblings found in 100 replicate simulations for a range of adult abundances. For each replicate, the total sibling matches include all cross-cohort HSPs among 5 years of samples. Annual sample sizes were chosen such that the total expected number of sibling matches was 50. The simulations followed Scenario A ($\phi$=1, red lines and circles) and Scenario B ($\phi$=10, blue lines and Xs) as defined in Table 3.

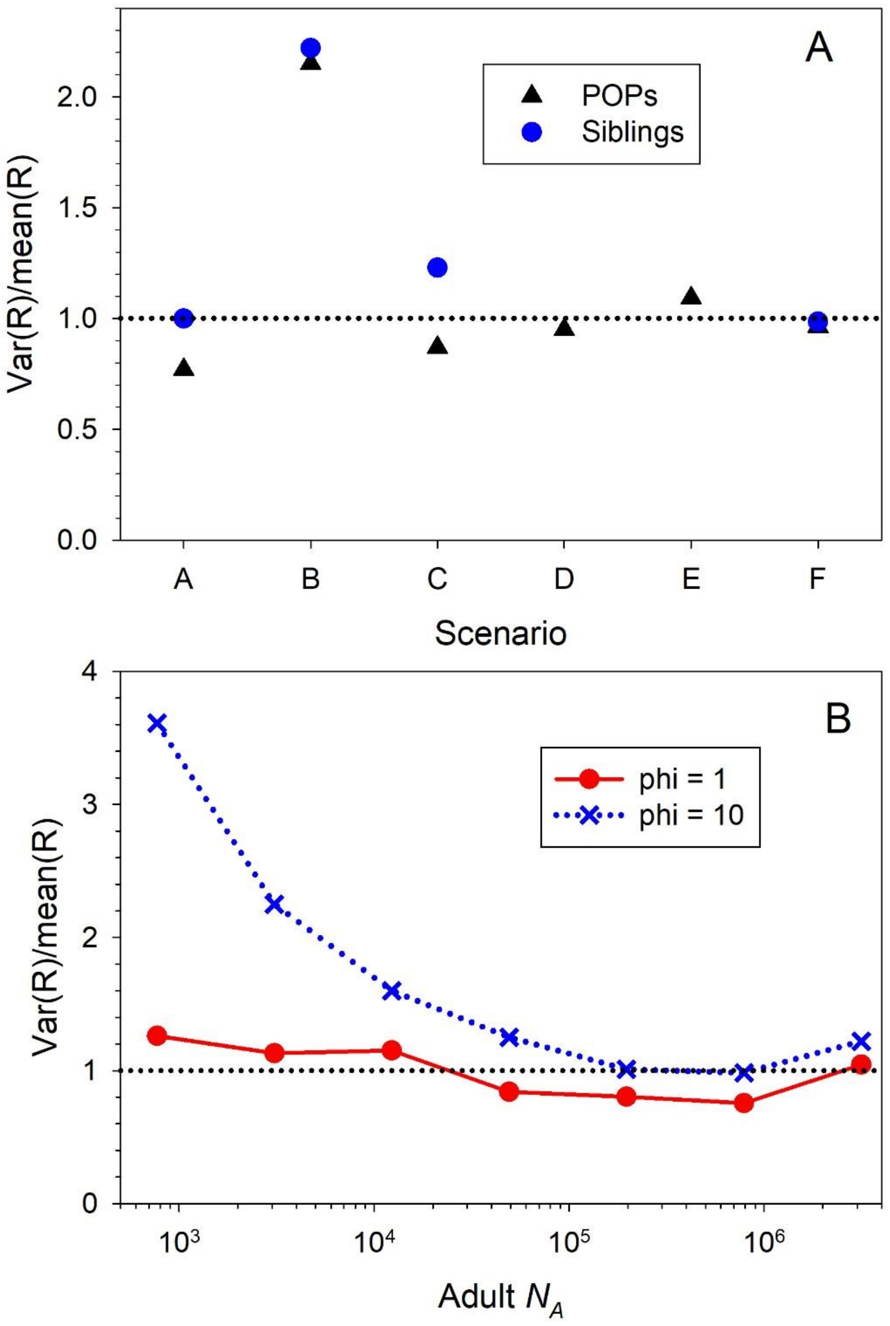

Figure S2. The variance-to-mean ratio for the number of close-kin matches (*R*) found in the simulations. For siblings, results apply to the total number of cross-cohort HSPs found using all 5 years of data. If the number of close-kin matches is Poisson distributed, the expected ratio is 1 (horizontal dotted lines). Panel A: results for scenarios shown in Table 3 for both POPs and siblings. Changes to adult sampling that define Scenarios D and E for POPs do not affect siblings, so no results are shown in those cases. Panel B: results for HSPs only for large $N_A$ simulations shown in Figures 5C and S1. These simulations followed Scenario A (ϕ=1, red lines and circles) and Scenario B (ϕ=10, dotted blue lines and Xs) as defined in Table 3.

### Computer code

The following R code can be used with the Windows version of CKMRPop to simulate the kind of data used in this study. Instructions can be found at https://eriqande.github.io/CKMRpop/index.html. We used R version 4.0.4 with Rtools 4.0. The following initial steps might be required:

```
install.packages("remotes")
remotes::install_github("eriqande/CKMRpop")
library(CKMRpop)
install_spip(Dir = system.file(package = "CKMRpop"))
```

The code can be tested it by running through the code in this vignette:
https://eriqande.github.io/CKMRpop/articles/species_1_simulation.html

++++++++++++++++++++++++++++++++++++++++++++++++++++++++++++++++

**## R code for doing POP simulations**

```
library(tidyverse)
library(hexbin)
library(rdist)
library(CKMRpop)

SPD <- species_1_life_history

### These options are set to run Scenario A from Table 3; options can be toggled/modified for other Scenarios

### define the scenario
survival = 0.7
alpha = 3  ## age at maturity
omega = 10 ## maximum age
adultlifespan = omega-alpha+1
phi = 1 ## ratio of Vk to kbar
##femalefecundity = c(0,0,alpha:omega)  ## fecundity proportional to age
femalefecundity = c(0,0,rep(1,adultlifespan))  ## constant fecundity
##femalefecundity = c(rep(0,9),1)  ## sweepstakes RS; only BOFFFs reproduce
cohort_size <- 2000
SampleSize = 300 ## fixed number of offspring to subsample from each cohort in year t+1
SampleSizeParents = SampleSize  ## fixed number of potential parents to subsample
##SampleSizeParents = 50  ## for sweepstakes because there aren't many age 10 individuals
samp_frac <- 1
sampspace = paste(samp_frac," ")
SPD$`number-of-years` <- 30  # run the sim forward for 30 years
samp_start_year <- 24
samp_stop_year <- 25
range = paste(samp_start_year,"-",samp_stop_year,sep="")
```

```
NReps = 50  ## used 500 in reported results
SPD[[4]] = c(0,0,1,1,1,1,1,1,1,1)  ## prob of reproducing at each age
##SPD[[4]] = c(0,0,1,0,1,0,1,0,1,0)
##SPD[[4]] = c(0,0,rep(0.1,8))
SPD[[6]] = femalefecundity
samplingplan <- paste(rep(sampspace, omega), collapse = "")
selective = alpha:omega/10
bit = "1 0 "
##sweepstakes <- "1 0 0 0 0 0 0 0 0 1"
##samplingpost <- noquote(c(range,sweepstakes))
##samplingpost <- noquote(c(range,samplingplan))
samplingpost <- noquote(c(range,bit,selective))

  a=as.numeric(Sys.time())
  set.seed(a)   ### gives a different result each time

SPD[[1]] = omega
SPD[[2]] = c(1,rep(survival,(SPD[[1]]-1)))
SPD[[3]] = SPD[[2]]
SPD[[5]] = SPD[[4]]
SPD[[7]] = SPD[[6]]
SPD[[8]] = "negbin"
SPD[[9]] = 1/phi
SPD[[10]] = SPD[[9]]
SPD[[11]] = -1
SPD[[12]] = 0.5

L <- leslie_from_spip(SPD, cohort_size)

# then we add those to the spip parameters
SPD$`initial-males` <- floor(L$stable_age_distro_fem)
SPD$`initial-females` <- floor(L$stable_age_distro_male)

# tell spip to use the cohort size
SPD$`cohort-size` <- paste("const", cohort_size, collapse = " ")
SPD$`fixed-cohort-size` <- ""  # define this flag and give it an empty string as an argument

SPD$`discard-all` <- 0
#SPD$`gtyp-ppn-fem-pre` <- paste(range,samplingplan)
#SPD$`gtyp-ppn-male-pre` <- SPD$`gtyp-ppn-fem-pre`
SPD$`gtyp-ppn-fem-post` <- samplingpost
SPD$`gtyp-ppn-male-post` <- SPD$`gtyp-ppn-fem-post`

Born = as.integer((samp_start_year:samp_stop_year)-2)
YearsSamp = length(Born)
BigCohorts = matrix(NA,NReps,YearsSamp)
```

```
colnames(BigCohorts) = Born
BigParents = matrix(NA,NReps,2)
colnames(BigParents) = c("Dads","Moms")
BigPOPs = BigParents
BigNbDads = rep(NA,NReps)
BigNbMoms = rep(NA,NReps)
TrueNbDads = rep(NA,NReps)
TrueNbMoms = rep(NA,NReps)
BigSampledParents = matrix(NA, NReps,(1+ omega-alpha))
BigSexes = matrix(NA,NReps,2)
colnames(BigSexes) = c("Dads","Moms")
colnames(BigSampledParents) = alpha:omega

######### start simulation
for (R in 1:NReps)  {

print(paste0("Replicate = ",R))
flush.console()

spip_dir <- run_spip(
  pars = SPD
)
# now read that in and find relatives within the one-generation pedigree
slurped <- slurp_spip(spip_dir, 1)

# First, get the non-genotype info for each individual all together
non_geno_stuff <- slurped$samples %>%
  mutate(YearSampled = map_int(samp_years_list, 1))  %>%
  select(ID, born_year, YearSampled, sex) %>%
  left_join(slurped$pedigree %>% select(kid, ma, pa), by = c("ID" = "kid"))  %>%
  rename(
    sampleID = ID,
    YearBorn = born_year,
    Sex = sex,
    Mom = ma,
    Dad = pa
  )   # change the column names
Pedigree = non_geno_stuff

## get samplesize each year
a = table(Pedigree$YearSampled)
##BigCohorts[R,] = a$freq

## add ages
Ages = Pedigree$YearSampled - (Pedigree$YearBorn)
table(Ages)
```

```
PedigreePlus = cbind(Pedigree,Ages)

### separate the offspring and parents
Offspring = subset(PedigreePlus,PedigreePlus$YearSampled == samp_stop_year &
PedigreePlus$Ages == 1)
PotentialParents = subset(PedigreePlus,PedigreePlus$YearSampled != samp_stop_year)  ## all
individual alive when Offspring were produced
Parents = subset(PotentialParents,PotentialParents$Ages >= alpha )  ## restrict to those who
were mature when Offspring were produced
BigParents[R,1] = length(Parents$Sex[Parents$Sex == "M"])
BigParents[R,2] = length(Parents$Sex[Parents$Sex == "F"])
## get True Nb

RSmoms = table(Offspring$Mom)
SSmoms = sum(RSmoms^2)
  if(SSmoms > sum(RSmoms))  { TrueNbMoms[R] = (sum(RSmoms)-
1)/(SSmoms/sum(RSmoms)-1) } else {TrueNbMoms[R] = 99999}
RSdads = table(Offspring$Dad)
SSdads = sum(RSdads^2)
  if(SSdads > sum(RSdads))  {TrueNbDads[R] = (sum(RSdads)-1)/(SSdads/sum(RSdads)-1) }
else {TrueNbDads[R] = 99999}

######### subsample offspring and parents
SampledOffspring = Offspring[sample(nrow(Offspring),SampleSize,replace=F),]
SampledParents = Parents[sample(nrow(Parents),SampleSizeParents,replace=F),]
MaleParents = subset(SampledParents,SampledParents$Sex == "M")
FemaleParents = subset(SampledParents,SampledParents$Sex == "F")
BigSexes[R,1] = nrow(MaleParents)
BigSexes[R,2] = nrow(FemaleParents)

b = table(SampledParents$Ages)
BigSampledParents[R,1:length(b)] = b

### Get distribution of RS of sampled parents
POPHitsMom = 1:nrow(FemaleParents)
POPHitsDad = 1:nrow(MaleParents)
for (j in 1:nrow(MaleParents)) {
  hits = SampledOffspring$Dad[SampledOffspring$Dad == MaleParents$sampleID[j]]
  POPHitsDad[j] = length(hits)  }
for (j in 1:nrow(FemaleParents)) {
  hits = SampledOffspring$Mom[SampledOffspring$Mom == FemaleParents$sampleID[j]]
  POPHitsMom[j] = length(hits)  }

BigPOPs[R,1] = sum(POPHitsDad)
BigPOPs[R,2] = sum(POPHitsMom)
```

```
  }  # end for R

##BigSampledParents
Comparisons = SampleSize*BigSexes
Nhat = Comparisons/BigPOPs
BigN = rowSums(Nhat)
IN = 1/Nhat[,1] + 1/Nhat[,2]

TrueNb = 4*TrueNbMoms*TrueNbDads/(TrueNbMoms + TrueNbDads)
##cbind(TrueNbMoms,TrueNbDads,TrueNb)
##c(Nhat,sum(Nhat))
1/mean(1/BigN)  ## harmonic mean naive estimate of adult abundance

NData = matrix(NA,5,1)
rownames(NData) = c("5%","Median","95%","CV1","CV2")
NData[1:3] = quantile(BigN,probs = c(0.05, 0.5, 0.95))
NData[4] = sd(BigN)/mean(BigN)
NData[5] = sd(1/BigN)/mean(1/BigN)

2*mean(BigPOPs)  ## average number of POPS
1/mean(1/TrueNb) ## harmonic mean true Nb

NData  ## distribution of N^
## CV1 = CV(N^)
## CV2 = CV(1/N^)

cohort_size
RPOPs = rowSums(BigPOPs)
var(RPOPs)/mean(RPOPs)
```

++++++++++++++++++++++++++++++++++++++++++++++++++++++++++++++++

**###R code for doing sibling simulations**

```
library(tidyverse)
library(hexbin)
library(rdist)
library(CKMRpop)

### These options are set to run Scenario A from Table 3; options can be toggled/modified for other Scenarios

SPD <- species_1_life_history

### define the scenario
survival = 0.7
alpha = 3  ## age at maturity
omega = 10 ## maximum age
adultlifespan = omega-alpha+1
phi = 10 ## ratio of Vk to kbar
##femalefecundity = c(0,0,alpha:omega)  ## fecundity proportional to age
femalefecundity = c(0,0,rep(1,adultlifespan))  ## constant fecundity
##femalefecundity = c(rep(0,9),1)  ## sweepstakes RS; only BOFFFs reproduce
cohort_size <- 32000
SampleSize = 354 ## fixed number to subsample from each cohort
samp_frac <- 2*SampleSize/cohort_size  ## twice as large as target for subsampling
SPD$`number-of-years` <- 56  # run the sim forward for 56 years
samp_start_year <- 51
samp_stop_year <- 55
NReps = 100
SPD[[4]] = c(0,0,1,1,1,1,1,1,1,1)  ## prob of reproducing at each age
##SPD[[4]] = c(0,0,1,0,1,0,1,0,1,0)
##SPD[[4]] = c(0,0,rep(0.1,8))
SPD[[6]] = femalefecundity

  a=as.numeric(Sys.time())
  set.seed(a)

##Scenario B
SPD[[1]] = omega
SPD[[2]] = c(1,rep(survival,(SPD[[1]]-1)))
SPD[[3]] = SPD[[2]]

SPD[[5]] = SPD[[4]]

SPD[[7]] = SPD[[6]]
SPD[[8]] = "negbin"
SPD[[9]] = 1/phi
SPD[[10]] = SPD[[9]]
```

```
SPD[[11]] = -1
SPD[[12]] = 0.5

L <- leslie_from_spip(SPD, cohort_size)

# then we add those to the spip parameters
SPD$`initial-males` <- floor(L$stable_age_distro_fem)
SPD$`initial-females` <- floor(L$stable_age_distro_male)

# tell spip to use the cohort size
SPD$`cohort-size` <- paste("const", cohort_size, collapse = " ")
SPD$`fixed-cohort-size` <- ""  # define this flag and give it an empty string as an argument

sfspace = paste("0 ")
range = paste(samp_start_year,"-",samp_stop_year,sep="")
SPD$`discard-all` <- 0
SPD$`gtyp-ppn-fem-pre` <- paste(range, "0 ", samp_frac, paste(rep(sfspace, SPD$'max-age' - 2),
collapse = ""))
SPD$`gtyp-ppn-male-pre` <- SPD$`gtyp-ppn-fem-pre`

Born = as.integer((samp_start_year:samp_stop_year)-2)
YearsSamp = length(Born)
BigGapHalf = array(data=0, dim = c(YearsSamp,YearsSamp,NReps))
colnames(BigGapHalf) = c("Year1","Year2","Year3","Year4","Year5")
BigGapFull = BigGapHalf
BigCohorts = matrix(NA,NReps,YearsSamp)
colnames(BigCohorts) = Born
BigNb = matrix(NA,NReps,YearsSamp)
BigNbdads = BigNb
BigNbmoms = BigNb
BigSibsSame = array(data = 0, dim = c(YearsSamp,4,NReps))
dimnames(BigSibsSame)[[2]] = c("cohort","FSP","PHSP","MHSP")
BigSibsSame[,1,] = Born

######### This function provided by Ryan Waples
GetSibs <- function(Pedigree2)  {
# remove duplicate rows in the pedigree
Pedigree2 = Pedigree2[!duplicated(Pedigree2), ]

# convert the parent IDs to unique integers (faster to compare)
unique_moms = sort(unique(Pedigree2$Mom))
unique_dads = sort(unique(Pedigree2$Dad))
mom_range = 1:length(unique_moms)
dad_range = 1:length(unique_dads)
names(mom_range) = unique_moms
names(dad_range) = unique_dads
```

```
Pedigree2$Mom_id = mom_range[Pedigree2$Mom]
Pedigree2$Dad_id = dad_range[Pedigree2$Dad]

# make n x n matrix (n=number of offspring) values are zero if the pair of offspring shares a
parent, positive otherwise
mom_matrix = pdist(Pedigree2$Mom_id)
dad_matrix = pdist(Pedigree2$Dad_id)
# extract the inds sharing parents, don't double count, and sort
mom_matches = which(mom_matrix==0,arr.ind = T)
mom_matches = mom_matches[mom_matches[,1] < mom_matches[,2], ]
dad_matches = which(dad_matrix==0,arr.ind = T)
dad_matches = dad_matches[dad_matches[,1] < dad_matches[,2], ]
mom_matches = mom_matches[order(mom_matches[,1], mom_matches[,2]),]
dad_matches = dad_matches[order(dad_matches[,1], dad_matches[,2]),]

# convert to data.frame - i1, i2 gives the individual - row number in the original pedigree file
mom_df = data.frame(mom_matches)
names(mom_df) = c('i1', 'i2')
mom_df$parent = 'mom'
dad_df = data.frame(dad_matches)
names(dad_df) = c('i1', 'i2')
dad_df$parent = 'dad'

# merge the dfs of the mom and dad matches to find full sibs
sibs = merge(mom_df, dad_df, by = c('i1', 'i2'), all=T)
# arrays below are boolean indexes into the sibs df, telling us how the pair shares parents
share_both = !(is.na(sibs$parent.x) | is.na(sibs$parent.y))
share_mom = is.na(sibs$parent.x)
share_dad = is.na(sibs$parent.y)
# we can count the number of pairs in each category
##sum(share_both)
##sum(share_mom)
##sum(share_dad)

# construct the output file
ms = cbind(Pedigree2[sibs[share_mom,]$i1,][, c('sampleID', 'YearBorn')],
  Pedigree2[sibs[share_mom,]$i2,][, c('sampleID', 'YearBorn')])
names(ms) = c('ID1', 'Born1', 'ID2', 'Born2')
ms$Parent = 'mom'

ds = cbind(Pedigree2[sibs[share_dad,]$i1,][, c('sampleID', 'YearBorn')],
          Pedigree2[sibs[share_dad,]$i2,][, c('sampleID', 'YearBorn')])
names(ds) = c('ID1', 'Born1', 'ID2', 'Born2')
ds$Parent = 'dad'

if (sum(share_both)>0)  {
```

```
  fs = cbind(Pedigree2[sibs[share_both,]$i1,][, c('sampleID', 'YearBorn')],
    Pedigree2[sibs[share_both,]$i2,][, c('sampleID', 'YearBorn')])
  names(fs) = c('ID1', 'Born1', 'ID2', 'Born2')
  fs$Parent = 'both'
  all_sibs = rbind(fs, ms, ds)  }
else {  all_sibs = rbind(ms, ds)  }

return(all_sibs)  }  # end function

######### start simulation

for (R in 1:NReps)  {

print(paste0("Replicate = ",R))
flush.console()

BigSibsHalf = matrix(0,YearsSamp,YearsSamp)
BigSibsFull = BigSibsHalf

spip_dir <- run_spip(
  pars = SPD
)
# now read that in and find relatives within the one-generation pedigree
slurped <- slurp_spip(spip_dir, 1)

# First, get the non-genotype info for each individual all together
non_geno_stuff <- slurped$samples %>%
  mutate(YearSampled = map_int(samp_years_list, 1))  %>% # this gets the sample year out of
the samp_years_list
  select(ID, born_year, YearSampled, sex) %>%  # pick out column in the order desired
  left_join(slurped$pedigree %>% select(kid, ma, pa), by = c("ID" = "kid"))  %>%  # add mom
and dad on there
  rename(
    sampleID = ID,
    YearBorn = born_year,
    Sex = sex,
    Mom = ma,
    Dad = pa
  )   # change the column names to what Robin wants
Pedigree = non_geno_stuff

## get cohort size each year
a = table(Pedigree$YearBorn)
BigCohorts[R,] = a

## get total Nb each year
```

```
Nbmoms = 1:YearsSamp
Nbdads = 1:YearsSamp

for (j in 1:YearsSamp)  {
year = Born[j]
cohort = subset(Pedigree,Pedigree$YearBorn == year)
RSmoms = table(cohort$Mom)
SSmoms = sum(RSmoms^2)
  if(SSmoms > sum(RSmoms))  { Nbmoms[j] = (sum(RSmoms)-1)/(SSmoms/sum(RSmoms)-1)
}
  else {Nbmoms[j] = 99999}
RSdads = table(cohort$Dad)
SSdads = sum(RSdads^2)
  if(SSdads > sum(RSdads))  {Nbdads[j] = (sum(RSdads)-1)/(SSdads/sum(RSdads)-1) }
  else {Nbdads[j] = 99999}
}

BigNbmoms[R,] = Nbmoms
BigNbdads[R,] = Nbdads
BigNb[R,] = 4*Nbmoms*Nbdads/(Nbmoms+Nbdads)

######### subsample cohorts of offspring
Pedigree2 = Pedigree[1,]

for (j in 1:YearsSamp)  {
  year = Born[j]
  cohort = subset(Pedigree,Pedigree$YearBorn == year)
  sampled = cohort[sample(nrow(cohort),SampleSize,replace=F),]
Pedigree2 = rbind(Pedigree2,sampled)
}  # end for j
Pedigree2 = Pedigree2[-1,]
Pegigree2 = as.data.frame(Pedigree2)

sibs = GetSibs(Pedigree2)

## get age gaps between sibs
Gap = abs(sibs$Born2 - sibs$Born1)

MHSP = subset(sibs,sibs$Parent == "mom")
PHSP = subset(sibs,sibs$Parent == "dad")
Halfs = rbind(MHSP,PHSP)
FSP = subset(sibs,sibs$Parent == "both")

##get within cohort sibs
withinhalf = subset(Halfs,Halfs$Born1 == Halfs$Born2)
```

```
 for(j in 1:YearsSamp)  {
      bit = subset(withinhalf,withinhalf$Born1 == Born[j])
      BigSibsHalf[j,j] = nrow(bit)
   }  # end for j
withinfull = subset(FSP,FSP$Born1 == FSP$Born2)
    for(j in 1:YearsSamp)  {
      bit2 = subset(withinfull,withinfull$Born1 == Born[j])
      BigSibsFull[j,j] = nrow(bit2)
      }## end for j

##get arcoss cohort sibs
acrosshalf = subset(Halfs,Halfs$Born1 != Halfs$Born2)
 for(j in 1:(YearsSamp-1))  {
 for(k in (j+1):YearsSamp)  {
    bit = subset(acrosshalf,acrosshalf$Born1 == Born[j] & acrosshalf$Born2 == Born[k])
    BigSibsHalf[j,k] = nrow(bit)
    }} # end for j,k
acrossfull = subset(FSP,FSP$Born1 != FSP$Born2)
 for(j in 1:(YearsSamp-1))  {
 for(k in (j+1):YearsSamp)  {
    bit = subset(acrossfull,acrossfull$Born1 == Born[j] & acrossfull$Born2 == Born[k])
    BigSibsFull[j,k] = nrow(bit)
    }} # end for j,k

BigGapHalf[,,R] = BigSibsHalf
BigGapFull[,,R] = BigSibsFull

  }  # end for R

########################

ComparisonsWithin = SampleSize*(SampleSize-1)/2

WangR = matrix(0,NReps,YearsSamp)
for (R in 1:NReps)  {
  matches = 1:YearsSamp
  for (k in 1:YearsSamp)  {
  matches[k] = 2*BigGapFull[k,k,R] + BigGapHalf[k,k,R] }
  WangR[R,] = matches
  }  ## end for R
WangNb = 4*ComparisonsWithin/WangR

TotWangR = matrix(0,NReps,YearsSamp-1)
colnames(TotWangR) = c("2Years","3Years","4Years","5Years")
for (R in 1:NReps)  {
for (j in 1:(YearsSamp-1))  {
```

```
  TotWangR[R,j] = sum(WangR[R,1:(1+j)])
  }  # end for j
  }  # end for R

ComboWangNb = matrix(0,NReps,YearsSamp-1)
for (j in 2:YearsSamp)  {
Top = 4*ComparisonsWithin*j
ComboWangNb[,j-1] = Top/TotWangR[,j-1]  }

WangData = matrix(NA,5,YearsSamp-1)
colnames(WangData) = c("2Years","3Years","4Years","5Years")
rownames(WangData) = c("5%","Median","95%","CV1","CV2")
for (j in 1:(YearsSamp-1))  {
WangData[1:3,j] = quantile(ComboWangNb[,j],probs = c(0.05, 0.5, 0.95))
WangData[4,j] = sd(ComboWangNb[,j])/mean(ComboWangNb[,j])
WangData[5,j] = sd(1/ComboWangNb[,j])/mean(1/ComboWangNb[,j])
}

AcrossHalfs = matrix(0,NReps,(YearsSamp-1))
colnames(AcrossHalfs) = c("2Years","3Years","4Years","5Years")
for (R in 1:NReps)  {
f=BigGapHalf[,,R]
f[!upper.tri(f)] <- 0
  for (j in 1:(YearsSamp-1))  {
    temp = f[1:(j+1),1:(j+1)]
  AcrossHalfs[R,j] = sum(temp)
  }  # end for j
  }  # end for R

gaps = 1:(YearsSamp-1)

ComparisonsBetween = matrix(2*SampleSize^2,YearsSamp-1,YearsSamp-1)
colnames(ComparisonsBetween) = c("Year2","Year3","Year4","Year5")
rownames(ComparisonsBetween) = c("Year1","Year2","Year3","Year4")
ComparisonsBetween[lower.tri(ComparisonsBetween)] <- NA
AdjustedComparisonsBetween = ComparisonsBetween

K = c(1,rep(survival,omega-1))
Lx = cumprod(K)
Nx = Lx*cohort_size
AdultN = sum(Nx[alpha:omega])
effectivesurvival = 1:4
for (j in 1:4)  {
effectivesurvival[j] = sum(Nx[(alpha+j):omega])/AdultN
}  # end for j
```

```
for (j in 1:(YearsSamp-1))  {
for (k in j:(YearsSamp-1)) {
  q = 1+k-j
  AdjustedComparisonsBetween[j,k] = ComparisonsBetween[j,k]*effectivesurvival[q]
  }}
TotalAdjustedComparisons = 1:(YearsSamp-1)
for (j in 1:(YearsSamp-1)) {
  temp = AdjustedComparisonsBetween[1:j,1:j]
  TotalAdjustedComparisons[j] = sum(temp,na.rm=T)
  }

X = t(2/AcrossHalfs)*TotalAdjustedComparisons
Nhat = t(X)

NData = matrix(NA,5,YearsSamp-1)
colnames(NData) = c("2Years","3Years","4Years","5Years")
rownames(NData) = c("5%","Median","95%","CV1","CV2")
for (j in 1:(YearsSamp-1))  {
NData[1:3,j] = quantile(Nhat[,j],probs = c(0.05, 0.5, 0.95))
NData[4,j] = sd(Nhat[,j])/mean(Nhat[,j])
NData[5,j] = sd(1/Nhat[,j])/mean(1/Nhat[,j])
}

SplitSibs = matrix(NA,NReps,YearsSamp)
TotAcross = rep(NA,NReps)
for (R in 1:NReps)  {
temp = rep(0,YearsSamp)
A = BigGapHalf[,,R]
##A[lower.tri(A,diag=FALSE)] <- 0
for (j in 1:YearsSamp)  {
for (k in j:YearsSamp)  {
  X = 1 + k-j
  temp[X] = temp[X] + A[j,k]
  }}  ## end for j,k
SplitSibs[R,] = temp
TotAcross[R] = sum(temp[2:YearsSamp])
} ## end for R

BigSplit = cbind(SplitSibs,TotAcross)
MeanGaps = colMeans(BigSplit)
VarGaps = 1:(YearsSamp+1)
for (j in 1:(YearsSamp+1))  {
VarGaps[j] = var(BigSplit[,j])
}  # end for j

GapCompare = rowSums(ComparisonsBetween,na.rm=T)
```

```
ESibs = GapCompare*effectivesurvival*2/AdultN
GapData = cbind(MeanGaps,VarGaps,VarGaps/MeanGaps,sqrt(VarGaps)/MeanGaps)
colnames(GapData) = c("MeanSibs","Var","Phi","CV")
rownames(GapData) = c(0:(YearsSamp-1),"TotAcross")

### harmonic means
1/mean(1/WangNb)
1/mean(1/BigNb)
1/mean(1/Nhat)

sum(BigGapFull,na.rm=T)  ## total full sibs
t(NData)
t(WangData)

GapData
ESibs

cohort_size
SampleSize
phi
```
